# A metabolic model to investigate the evolution of chemodiversity

**DOI:** 10.64898/2026.08.21.746203

**Authors:** Frans M. Thon, Meike J. Wittmann

## Abstract

1. Plants produce a great chemodiversity, which is the diversity of specialized metabolites (SMs). These SMs are produced in complex metabolic pathways and play an important role in inter-species interactions. There are numerous hypotheses about the evolutionary processes which brought about and maintain chemodiversity. Some have been partially tested in lab and field studies. However, some of their assumptions and predictions are better tested by quantitative modeling, and so far no quantitative model has investigated the role of metabolic pathways.
2. To close this gap, we developed an individual-based model for metabolic pathway evolution. It models enzymes creating metabolites with various modifications. Enzymes undergo inheritance and mutation. We used the model to compare the screening and interaction diversity hypotheses.
3. The screening hypothesis predicts promiscuous enzymes, genetic drift, the presence of many non-beneficial metabolites, and high metabolite richness. The interaction diversity hypothesis predicts specialized enzymes, selection, the almost exclusive presence of beneficial metabolites, and situation-dependent metabolite richness. We found that the patterns predicted by the screening hypothesis did not occur, while those predicted by the interaction diversity hypothesis did.
4. This provides reason to favor the interaction diversity hypothesis over the screening hypothesis when connecting empirical results to their evolutionary context.

## 1 Introduction

Chemodiversity, the diversity of specialized metabolites (SMs) plants produce (see Wetzel & Whitehead, 2020, for a review), is an important part of biodiversity. Chemodiversity exists at different scales, for example between individuals, within a population, or between populations (Thon *et al*., 2024).

Many SMs affect interactions between plants and herbivores. Whether they increase or decrease herbivory depends on the specifics of the interaction. Higher levels of chemodiversity may increase herbivore resistance (Richards *et al*., 2016; Ziaja & Müller, 2023), increase resistance against specialists while decreasing resistance against generalists (Wang *et al*., 2023), or the exact opposite (Richards *et al*., 2016). Additionally, a highly chemodiverse community ill-adapted to local conditions may be less herbivore-resistant than less diverse but more locally adapted communities (Fernandez-Conradi *et al*., 2022). We therefore need to understand the exact ecological and evolutionary mechanisms that underpin chemodiversity to effectively utilize it in agricultural settings or conservation efforts.

Quantitative modeling in biology can be used to investigate such evolutionary mechanisms, particularly when empirical experiments are impractical or even impossible (Servedio *et al*., 2014). There currently is more verbal theory on chemodiversity evolution than theoretical modeling (Thon *et al*., 2024). Existing models have found, for example, that between-plant trait diversity could result both in associational resistance and susceptibility depending on the exact trait (Hambäck *et al*., 2014) and that coevolution could make different defense chemicals beneficial at different points in time (Speed *et al*., 2015; Ashby & Boots, 2017). But none of these models, nor any other model we are aware of, model the metabolic pathways that underpin chemodiversity.

Plants produce their chemodiversity in complex metabolic pathways, often using promiscuous enzymes: enzymes that can catalyze reactions using multiple substrates (Jones & Firn, 1991; Bar-Even & Salah Tawfik, 2013). When every enzyme uses substrates produced by other enzymes, and also produces substrates for these other enzymes, a grid-like pathway emerges where the number of metabolites produced increases exponentially with every additional enzyme that can make a new modification. This can bring about a great chemodiversity within a plant using only a few enzymes (Jones & Firn, 1991).

All this complexity has led to numerous verbal hypotheses on the ecological and evolutionary mechanisms underpinning chemodiversity. Two such hypotheses are the screening hypothesis and the interaction diversity hypothesis.

**The screening hypothesis** (Jones & Firn, 1991) has often been used to put empirical findings in an evolutionary context (Owen & Peñuelas, 2006; Weng *et al*., 2012; Pais *et al*., 2018; Fernandez-Conradi *et al*., 2022). It makes two controversial assumptions (Pichersky *et al*., 2006) about how chemodiversity is maintained. First, it assumes that only a small proportion of all possible metabolites are biologically active and benefit the plant. Second, Jones & Firn (1991) argue that the costs of producing these metabolites is low enough for many of them to exist, but that producing numerous enzymes is sufficiently costly that they discuss several ways in which a high diversity can be produced with few enzymes. This implies that producing enzymes is substantially more costly than producing metabolites (Jones & Firn, 1991). Therefore, the screening hypothesis predicts that chemodiversity is produced by a system of constantly evolving and highly promiscuous enzymes. These enzymes are proposed to have evolved from enzymes involved with the production of secondary metabolites that have important properties, but with a high tolerance for variance, such as the large, substitutable variation of lipids found in cell membranes (Firn, 2009). The key idea is that high chemodiversity is favored because it allows plants to ‘stumble upon’ the rare useful metabolites by chance.

In contrast, the **interaction diversity hypothesis** assumes that different metabolites are involved with different interactions plants have with other organisms. Therefore, the sheer number of these interactions can explain the large number of SMs plants produce, even if the exact function of every metabolite is not known at present. This idea was often implicitly assumed as a ‘common sense hypothesis’ in chemodiversity research, without being explicitly named (Berenbaum & Zangerl, 1996). It was clearly formulated as a distinct hypothesis by Whitehead *et al*. (2021). The assumptions and predictions of the screening hypothesis and the interaction diversity hypothesis are discussed in more detail below and in Table 1.

**Table 1:** Table of assumptions and predicted quantitative effects of the screening and interaction diversity hypotheses, as well as how these assumptions are implemented and the predictions evaluated through measurements.

| Screening hypothesis | Interaction diversity hypothesis |  |
| --- | --- | --- |
| Assumptions |  | Implementation |
| Cost enzymes ≫ Cost metabolites | Cost enzymes ? Cost metabolites | Vary enzyme cost |
| Few metabolites with positive fitness effect | Number of metabolites with positive fitness effects depends on number of interactions | Vary number of beneficial metabolites |
| Predictions |  | Measurement |
| Promiscuous enzymes | Specialized enzymes | Promiscuity index: mean median $k_{cat}/K_M$ |
| Dominated by drift | Dominated by selection | Difference in number of gene copies and alleles through time |
| Present metabolites random | Present metabolites closely match beneficial ones | Ratio of beneficial metabolites |
| High metabolite richness | Metabolite richness dependent on number of beneficial metabolites | Metabolite richness |

Here, we build on previous work testing these hypotheses. Whitehead *et al*. (2021) performed bioassays to test them, rearing caterpillars and growing fungi on media containing different mixtures of commercially available compounds. All the compounds tested showed biological activity, in line with the interaction diversity, but not screening hypothesis. However, Whitehead *et al*. (2021) recognize that there is little market for compounds with no known biological activity. We agree that this limits their study’s ability to disprove the screening hypothesis outright. Modeling approaches do not face this issue. Wittmann & Bräutigam (2025) modeled the screening and interaction diversity hypothesis (along with others), providing valuable insights especially into the role of genetic dominance in bringing about chemodiversity. They found that plants produced a higher chemodiversity when interacting with more herbivores, in apparent support of the interaction diversity hypothesis. Additionally, more chemodiverse populations maintained higher fitnesses after an invasion by a new herbivore, having ‘stumbled upon’ beneficial metabolites while producing a high diversity of compounds, matching the screening hypothesis. However, their model did not include the enzymes and pathways that produce metabolites, and could not investigate the effects of the evolution thereof.

In this article, we present an individual-based model of chemodiversity with a particular focus on the evolution of enzymes and metabolic pathways. We aimed to create a general model that can incorporate the assumptions of multiple hypotheses, and evaluate their predictions. As a first application, we use it to perform proof-of-concept tests of the screening (Jones & Firn, 1991; Firn & Jones, 2003) and interaction diversity hypotheses (Whitehead *et al*., 2021), and propose avenues for future work.

## 2 Model description

We here give an overview of the individual-based simulation model. A more detailed model description following the ODD (Overview, Design concepts, Details) protocol (Grimm *et al*., 2006, 2020) can be found in supplement 1.

We model a population with a constant size of 1000 plants for 1000 generations. Each plant has its own heritable diploid genome. A plant may be homozygous at a locus with two copies of the same allele, heterozygous with two different ones, or have one or two empty alleles at that locus (which means that the population may have an absence-presence polymorphism, also known as indel polymorphism). Genes code for the characteristics of the plant’s enzymes, i.e. for a set of enzyme variables: the concentration in mMol/L; the function, that is, the modification the enzyme catalyzes; and two vectors of catalyzing rates (*k_cat_*, in *s^−^*^1^) and Michaelis constants (*K_M_*, in mMol/L) with entries for every substrate. All these variables can undergo mutation, as described in “Mutation” below. Depending on their variables, the enzymes then produce the different metabolites as described in the “Metabolism” submodel below.

SMs are modeled as follows: Unmodified metabolites can undergo a number of modifications (a carboxylation or desaturation for example). The metabolites are modeled as a binary string ‘code’ with as many bits as there are possible modifications. The bits of the unmodified metabolite are all 0. An enzyme with enzyme function *n* can irreversibly modify position *n* in this binary string from 0 to 1 in any metabolite that has not undergone this modification (Figure 1). In this article, we describe two sets of simulations. In the first, metabolites can undergo three modifications, resulting in genes with three possible enzyme functions and eight possible metabolites. Therefore, an enzyme with function ‘1’ catalyzes the reaction of a substrate 000 to a product 100, from 010 to 110, from 001 to 101, and from 011 to 111 (the light blue enzyme in Figure 1). In the other set of simulations, metabolites can undergo ten modifications, resulting in ten possible enzyme functions and 1024 possible metabolites.

**Figure 1:**
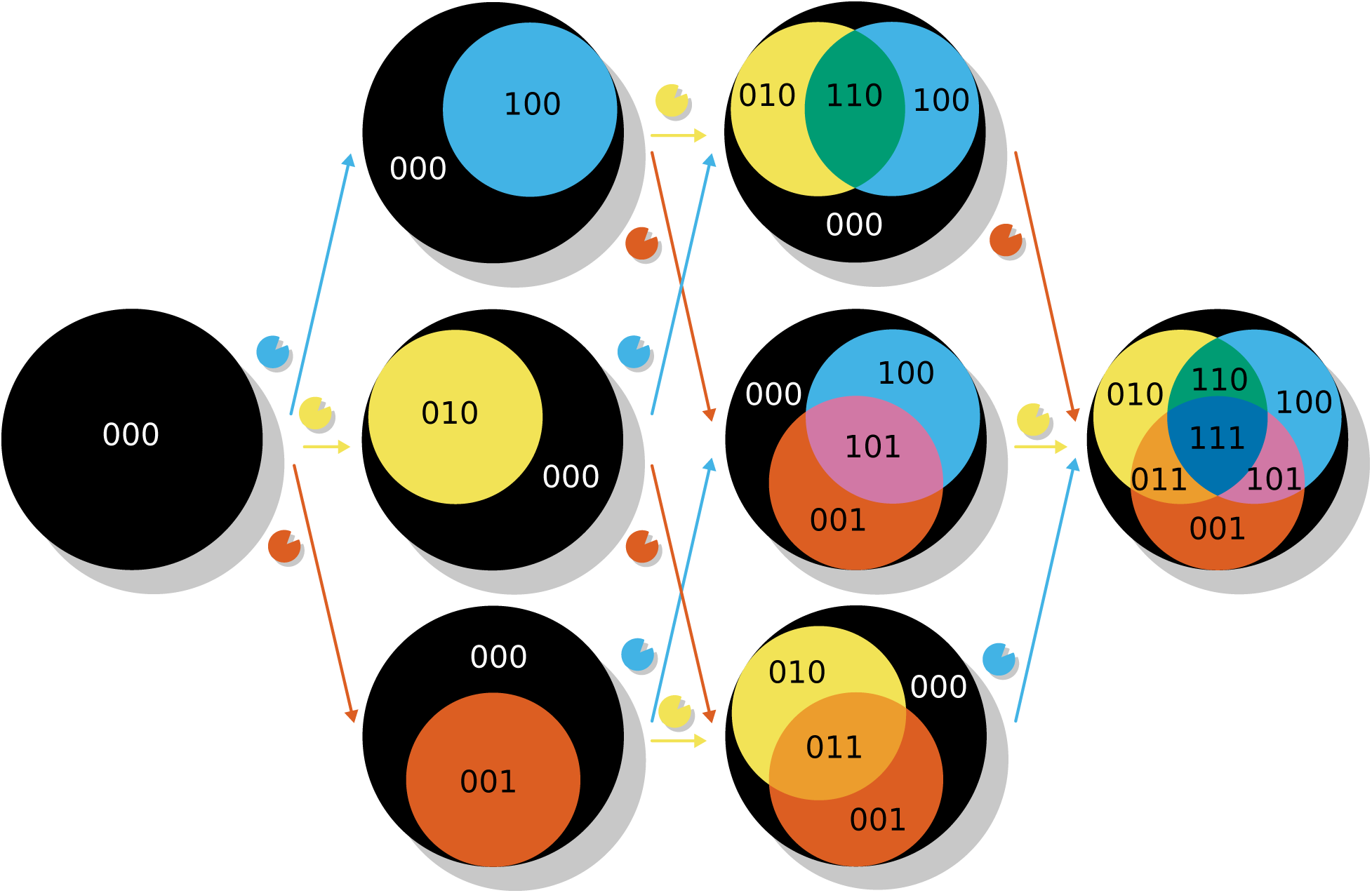
Illustration of the metabolic pathway, here with three possible modifications. Circles represent the metabolites present in a plant. The unmodified metabolite, in black, is always present. Each enzyme can catalyze one reaction, represented by arrows with the same colour as the corresponding enzyme. If all enzymes a plant contains catalyze the same reaction, a single derived metabolite is formed, shown by a single coloured circle in the first column of circles. Additional enzymes with different enzyme functions that can catalyze different reactions result in more metabolites, represented by coloured circles corresponding to the resulting metabolite with one modification. Additionally, the metabolites with multiple modifications are represented as the overlap in a Venn diagram. The number of metabolites produced increases exponentially with every additional enzyme with a new function (see also figure 3 in Firn & Jones, 2003).

The model’s 1000 generations are divided into ‘phases’ of 250 generations. Differences between phases reflect changes in the biotic environment, for example changes in the herbivore community due to invasion or local extinction. In each simulation run, a set number of derived metabolites convey a positive fitness effect. The specific metabolites that are beneficial are random within each phase, resulting in different ‘phase scenario’ sets (Supplement 3). Within each ‘phase scenario’ set, the scenario with one beneficial metabolite picks one random metabolite to be beneficial, and then adds another random metabolite for the scenarios with two beneficial metabolites, and so on. Thus, the scenarios with more beneficial metabolites of the same ‘phase scenario’ set always include the beneficial metabolites from scenarios of the same set with fewer beneficial metabolites (Supplement 3).

The model is initialized with identical plant individuals that each have as many homozygous loci as there are modifications possible, so either three or ten. The initial alleles either all produce enzymes that have the same metabolic function (that is, they modify their substrates in the same way; for example all loci have alleles that code for enzymes with function 1), or all have different functions (that is, there is one enzyme with each possible different enzyme function); produce an enzyme concentration of 0.01 mMol/L; have random kinetic variables (*k_cat_* and *K_M_*) drawn from a lognormal distribution (see process “Mutation” below); and a unique identifying ID. Initialization consists of two processes, “metabolism”, and “fitness calculation”. After initialization, the model life cycle consists of four processes in every generation (Figure 2): the aforementioned “metabolism” and “fitness calculation” processes, as well as “Reproduction” and “Mutation”. All these processes are described in more detail below. Every 10 generations, the metabolite concentrations, the genomes, and the fitness of each plant are recorded.

**Figure 2:**
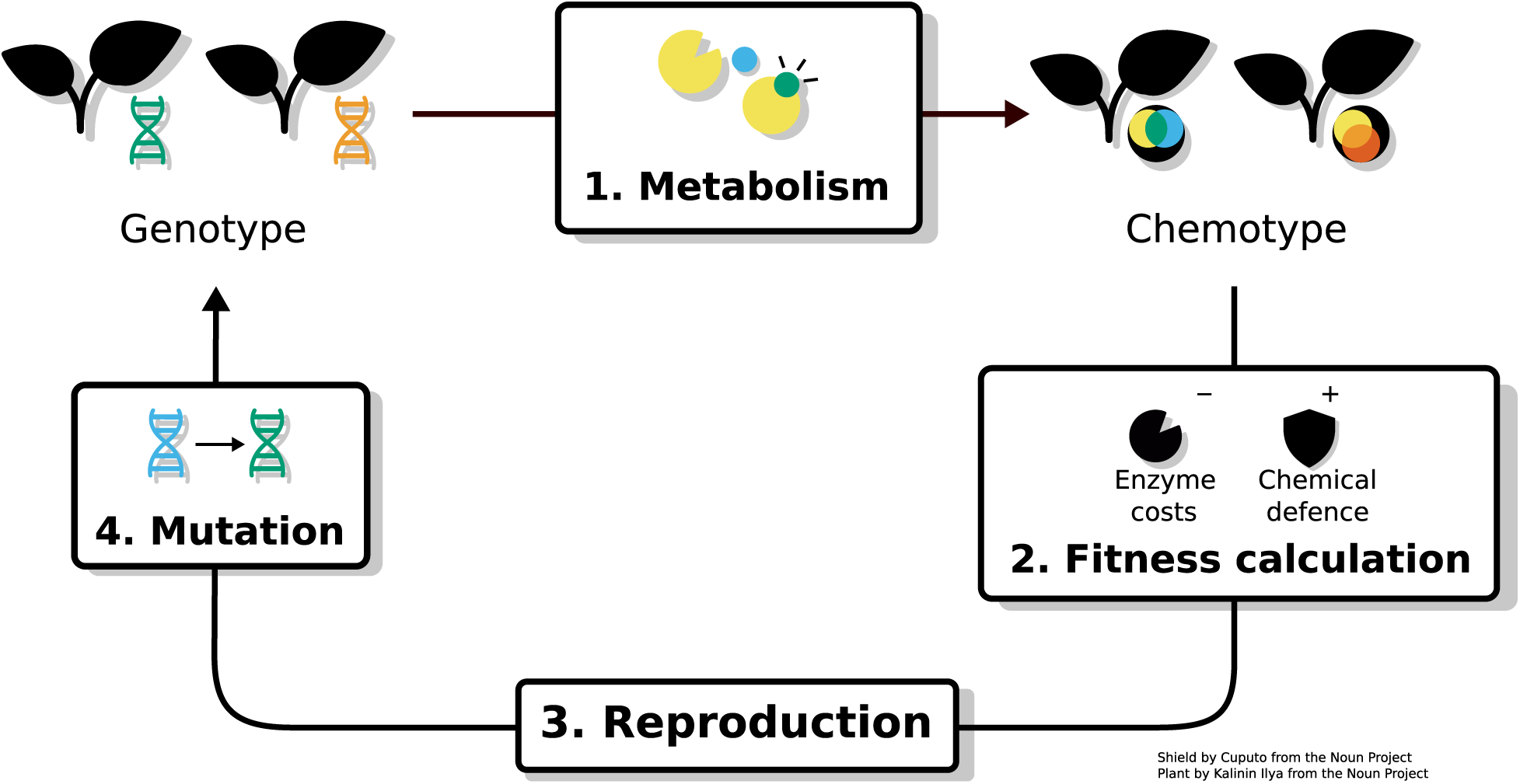
Overview of the model flow. A population consists of plants. The plants have genes, from which their enzymes are derived. These enzymes go into the metabolism, modeled as a system of ordinary differential equations based on Michaelis-Menten kinetics. The stable outcome of this system of ODEs are the concentrations of each metabolite, which form the chemotype. Then these metabolites have a fitness benefit based on the metabolite and its concentration, and the enzymes a fitness cost. Together, they determine the fitness of the plant. The fitnesses of the plants are then used to pick parent plants for the next generation during reproduction. The genes in the plants in the next generation may undergo mutation, and start the cycle anew.

### 1. Metabolism

Enzymes get their function, kinetic variables, and concentration from their corresponding genes. The creation of product metabolites from substrate metabolites is simulated as a set of irreversible enzyme-catalyzed reactions

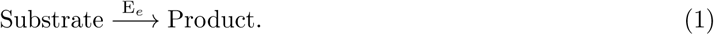

Here, every metabolite that has not yet undergone the modification an enzyme *E_e_* catalyzes is its substrate, and every metabolite that has is its product (Figure 1). The final concentrations are calculated using ordinary differential equations (ODEs) based on Michaelis-Menten kinetics. The change in concentration of the unmodified metabolite *M*_0_ is given by

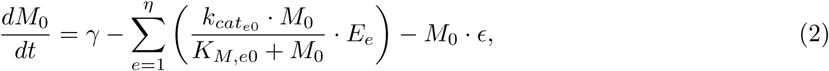

and the change in concentration of the other metabolites (*M_m_* for *m >* 0) is given by

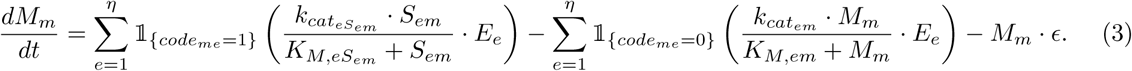

Here *E* is the vector of enzyme concentrations of length *η*, *M* is the vector of metabolite concentrations, *S* is the vector of vectors of substrate concentrations, so that *S_em_* is the concentration of the substrate metabolite that enzyme *E_e_* transforms into metabolite *M_m_*. For example, in a simulation with three modifications, the product metabolite *M*_6_ (011) of the enzyme *E_e_* with function 3, which modifies the third position in the metabolite, would be created from the substrate *S_e_*_6_, which is the metabolite *M*_2_ (010). *code_me_* is 1 if metabolite *m* is a possible product of enzyme *E_e_* and 0 if metabolite *M_m_* is a possible substrate of enzyme *E_e_* given the modification it carries out. *k_cat_* is the vector of catalyzing rates, and *K_M_* the vector of Michaelis constants. *γ* is the influx of *M*_0_ into the system, and *ɛ* the efflux of metabolites from the system.

### 2. Fitness calculation

The metabolite concentrations at the end of the metabolism submodel and their fitness effects determine the fitness of each plant according to the equation:

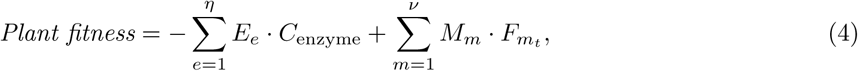

with *E* the vector of enzyme concentrations of length *η*, *C*_enzyme_ the fitness cost of enzyme production, *M* the vector of metabolite concentrations of length *ν*, and *F_m_t__* the fitness effect of metabolite *m* at the current time. In every generation, the fitness effects of beneficial metabolites are drawn from a uniform distribution between 0 and 500. This represents changes in the abiotic and/or biotic environment.

### 3. Reproduction

We randomly draw two parents from the population for each offspring plant, in a weighted lottery based on their fitness values. For each locus, a random gene copy at that locus (which can be an empty allele) is passed from each parent to the offspring plant. The offspring population replaces the parent population.

### 4. Mutation

Every gene copy in every plant may undergo any number of mutations of its variables (concentration, function, *k_cat_* and *K_M_*), be duplicated, or lost. Whenever a gene is duplicated or its variables are mutated, it is assigned a new unique ID. The probability that any variable of the allele is mutated or a duplication occurs is 10*^−^*^4^, a rapid mutation rate based on per-locus mutation rates in previous evolutionary models (Østman *et al*., 2012; Speed *et al*., 2015; Yeaman *et al*., 2018). This strikes a balance between allowing mutations to be selected away or becoming fixed over multiple generations, and simulation length. The probability a gene is lost is 10*^−^*^2^. Considering that a mutation that results in changes to the structure of an enzyme will change its affinities for all substrates in some way, for a mutation to a kinetic variable we draw new values of *k_cat_* and *K_M_* for all substrates independently from a lognormal distribution with *µ* = *−*3.45 and *σ* = 0.68. These values are chosen so that most of the resulting kinetic parameters are between 0.01 and 0.1 (mean 0.04, standard deviation 0.03, mode 0.02), the range of *K_M_* and *k_cat_* of enzymes found in the terpene pathway of *Salvia officinalis* (Croteau *et al*., 1994; Kampranis *et al*., 2007; Ignea *et al*., 2014). When an allele is duplicated, a new allele identical to the duplicated allele is put in a new locus of the genome of the mutated plant, with empty loci added to the genomes of all other plants. To prevent rare cases of uncharacteristically large genomes slowing down some simulations to a point that analysis becomes unfeasible, the maximum number of loci is set to 80. When the genomes in a simulation already have 80 loci and another duplication event occurs, this duplication is discarded (see Supplement 1 for details). A gene loss mutation leads to an empty allele.

### 2.1 Assumptions of the two hypotheses

The screening and interaction diversity hypotheses differ in two main assumptions, which are however not necessarily in conflict with each other (Table 1). First, the screening hypothesis assumes that while there is strong evolutionary pressure to produce fewer enzymes because of the fitness cost of producing them, the cost of producing numerous metabolites with few enzymes is low enough to allow the production of metabolites that are not beneficial, even over many generations (Jones & Firn, 1991; Firn & Jones, 2003). This implies that the production of enzymes is more expensive than the production of metabolites. The interaction diversity hypothesis, on the other hand, assumes nothing about the relative costs of enzymes and metabolites (Whitehead *et al*., 2021). To explore the role of enzyme costs, we simulate a ‘low’ (*C*_enzyme_ = 1) and ‘high’ (*C*_enzyme_ = 10) enzyme production cost scenario. Second, the screening hypothesis assumes that only few of all possible metabolites are biologically active, though which of these are beneficial may change with changing environmental factors and interactions (Jones & Firn, 1991; Firn & Jones, 2003; Firn, 2009) (In Firn (2009); Firn & Jones (2009) the authors leave space for ‘non-biomolecularly active’ compounds to have some function, but this ‘soft screening’ hypothesis is beyond the scope of this article). The interaction diversity hypothesis on the other hand makes no assumptions about the frequency of beneficial metabolites, but rather assumes that the number of metabolites that can convey positive fitness effects depends on the number of interactions the plant has with other organisms (Whitehead *et al*., 2021). We reflect this in two ways. First, in the three-modification simulations, we vary the number of beneficial derived metabolites in each phase between 1 and 7 (out of 7 derived metabolites), in 10 ‘phase scenario’ sets (different metabolites are beneficial in each ‘phase scenario’ set, Supplement 3). Sparse phase scenarios with few beneficial metabolites reflect the assumptions of the screening hypothesis. Both sparse and rich scenarios reflect the assumptions of the interaction diversity hypothesis, and the different ‘phase scenarios’ show us whether it matters what metabolites were beneficial in the past in a changed environment. However, seven derived metabolites is a small number. Even one beneficial metabolite in a system with only up to seven derived metabolites is quite a large proportion of beneficial metabolites for the screening hypothesis. After all, many groups of natural products, such as terpenes or alkaloids, comprise many thousands of different metabolites. Therefore, second, we run a set of ten-modification simulations, resulting in 1024 possible metabolites. Here, we vary the number of beneficial derived metabolites in each phase between 1, 10, and 100 beneficial metabolites out of 1023 in 10 ‘phase scenario’ sets. Neither the authors of the screening hypothesis nor any of the authors involved with the interaction diversity hypothesis make strong claims about where chemodiversity initially came from, though Firn & Jones (2003) claims that it must have existed before the evolution of land plants. Nevertheless, it must have come about at some point. To test whether chemodiversity can come about and be maintained under the assumptions of the two hypotheses, we therefore vary whether the initial alleles code for enzymes with identical or different functions, to test whether chemodiversity can come about or be maintained respectively.

### 2.2 Predictions of the two hypotheses

There are four ways in which the two hypotheses differ in how they predict the model to behave (Table 1).

First, the screening hypothesis predicts that plants will evolve promiscuous enzymes, giving rise to branched or grid-like pathways (branched pathways being essentially a special case of grid-like pathways) (Firn & Jones, 2003). These promiscuous enzymes are predicted to be prevented from becoming more specialized over time by changes in which metabolites are beneficial over time. The interaction diversity hypothesis on the other hand predicts plants to produce mostly currently beneficial metabolites (Whitehead *et al*., 2021), and therefore more specialized enzymes.

Enzyme promiscuity can be assessed using the specificity constant *k_cat_/K_M_*. The specificity constant is different for every substrate, with higher values indicating higher affinity. A specialized enzyme has a much higher specificity constant for one substrate than for the others, while a promiscuous enzyme has more evenly distributed specificity constants. We therefore propose a *promiscuity index*, which is the median normalized specificity constant of the enzyme over all substrates. Here, ‘normalized’ means that the specificity constants for all substrates are divided by the highest specificity constant of that enzyme. The median of these normalized specificity constants is our promiscuity index. High values mean the enzyme is promiscuous, low values that it is specific to the substrate with the highest specificity constant. The mean promiscuity index of the population indicates whether enzymes are more specific or more promiscuous in general. In our model, whenever the *k_cat_* or *K_M_* are mutated, new values of *k_cat_* or *K_M_* are randomly drawn for every substrate, making the mutated gene randomly more or less promiscuous than its parent gene. While mutation brings in new variation in promiscuity, selection can select for higher or lower promiscuity. For example, if enzyme promiscuity is beneficial, we would expect more promiscuous enzymes to become more common in the population. When there are no enzymes in the population, the promiscuity index is undefined. It should be noted that the absolute values of the promiscuity indices are not comparable between simulations with different numbers of modifications. Each enzyme has 7 values of *k_cat_/K_m_* in the three-modification model, and 1023 in the ten-modification model. When a higher number of values of *k_cat_/K_m_* is involved in calculating the promiscuity index, this value is expected to be lower.

Second, the screening hypothesis predicts selection to result in a metabolism where numerous metabolites are produced, not specific metabolites. Therefore, we would initially see selection for promiscuous enzymes, followed by genetic drift. The interaction diversity hypothesis on the other hand predicts selection to result in specific beneficial metabolites being produced. Therefore we would see selection after phase changes. We can track both how many alleles there are in sum across all loci and plants (sum of alleles across loci, SoA), and how many gene copies there are in total across all loci and plants (total number of gene copies across loci, NoG). For example, consider a population of 1000 plants with two loci. If, at locus 1, all plants are either homozygous for allele 1, homozygous for allele 2, or heterozygous with one copy of allele 1 and one copy of allele 2, and at locus 2, they have a presence-absence polymorphism with exactly one copy of an allele 3, this population would have a SoA of three, and a NoG of 3000. Together, SoA and NoG can inform us what kind of evolutionary dynamics are happening in the simulation. For genetic drift, the SoA and NoG should move to an equilibrium between mutation and random extinction and fixation of alleles. However, such an equilibrium could also be an equilibirium between directional selection and mutation. When there is an increase or decrease in the SoA and NoG, this indicates a change in selection pressures. For example, an increase in the SoA indicates that there is less selection against new alleles than previously. If the NoG also rises at the same time, there is less selection against all alleles. If the NoG stays the same or sinks instead, there is negative selection on previously common alleles.

Third, the screening hypothesis predicts that the metabolites produced should not change in response to a change in beneficial metabolites, as it assumes selection for a metabolism that produces numerous metabolites, not selection for the production of particular metabolites. The interaction diversity hypothesis on the other hand predicts that mostly beneficial metabolites are produced. This would have different effects on the ratio of beneficial metabolites in the simulation. This ratio is the sum of the concentration of beneficial metabolites, divided by the total amount of metabolite produced. When there are no metabolites, this beneficial ratio is undefined. Following the predictions of the screening hypothesis, we would expect no tendency towards a high ratio of beneficial metabolites. From the predictions of the interaction diversity hypothesis, we would expect both a high ratio of beneficial metabolites, and a quick selective response to phase changes. This selection could favor high copy numbers of alleles which produce enzymes with the right enzyme functions, alleles which produce enzymes with the right functions in high concentrations, or alleles with high enzyme efficiency.

Fourth, both hypotheses predict that their mechanisms can bring about and maintain chemodiversity. Therefore, we look at the mean richness of derived metabolites of the plants in the population to see how chemodiversity develops through time for the different experimental scenarios.

We used a three and a ten-modification model. As the number of metabolites increases exponentially with the number of modifications, higher-modification models both have longer runtimes, and the networks of beneficial metabolites are much more difficult to analyze. For this reason, we only covered the full range of high and low numbers of beneficial metabolites for the three-modification model.

The model was coded in C++, and ran from a wrapper in R 4.1.0 (R Core Team, 2021), parallelizing with the packages foreach (Microsoft & Weston, 2020) and doMC (Revolution Analytics & Weston, 2020). All analyses were performed in R 4.1.0 (R Core Team, 2021), using the packages reshape2 (Wickham, 2007), stringr (Wickham, 2023b) and forcats (Wickham, 2023a) for data handling, and the packages ggplot2 (Wickham, 2016), ggpubr (Kassambara, 2025), viridis (Garnier *et al*., 2024), and patchwork (Pedersen, 2025) for plotting. We performed a robustness analysis using the Morris method (Supplement 2, see Morris (1991); Campolongo *et al*. (2007); Imron *et al*. (2012)). The code to perform all simulations and analyses is available as a supplemental zip file (supplement 10).

## 3 Results

A typical run of the model is shown in Figure 3. Different chemotypes arise and are selected away in different phases (A). The amount of derived metabolites produced goes up over time, with different metabolites produced in different phases (B). The promiscuity index sinks steadily, indicating an increasing enzyme specialization (C). Differences in the sum of alleles across loci (SoA) indicate differences in selection pressures in different phases (D). The ratio of beneficial metabolites tends to be high (E), while the metabolite richness fluctuates depending on the phase (F). The model is highly stochastic. Even under the same parameter settings, different replicate populations may develop different chemotypes (see Figure S5.1, S5.2), so that we use summaries across replicates for our more general conclusions. In supplement 10, summary figures with the number of beneficial metabolites on the x axis can be found.

**Figure 3:**
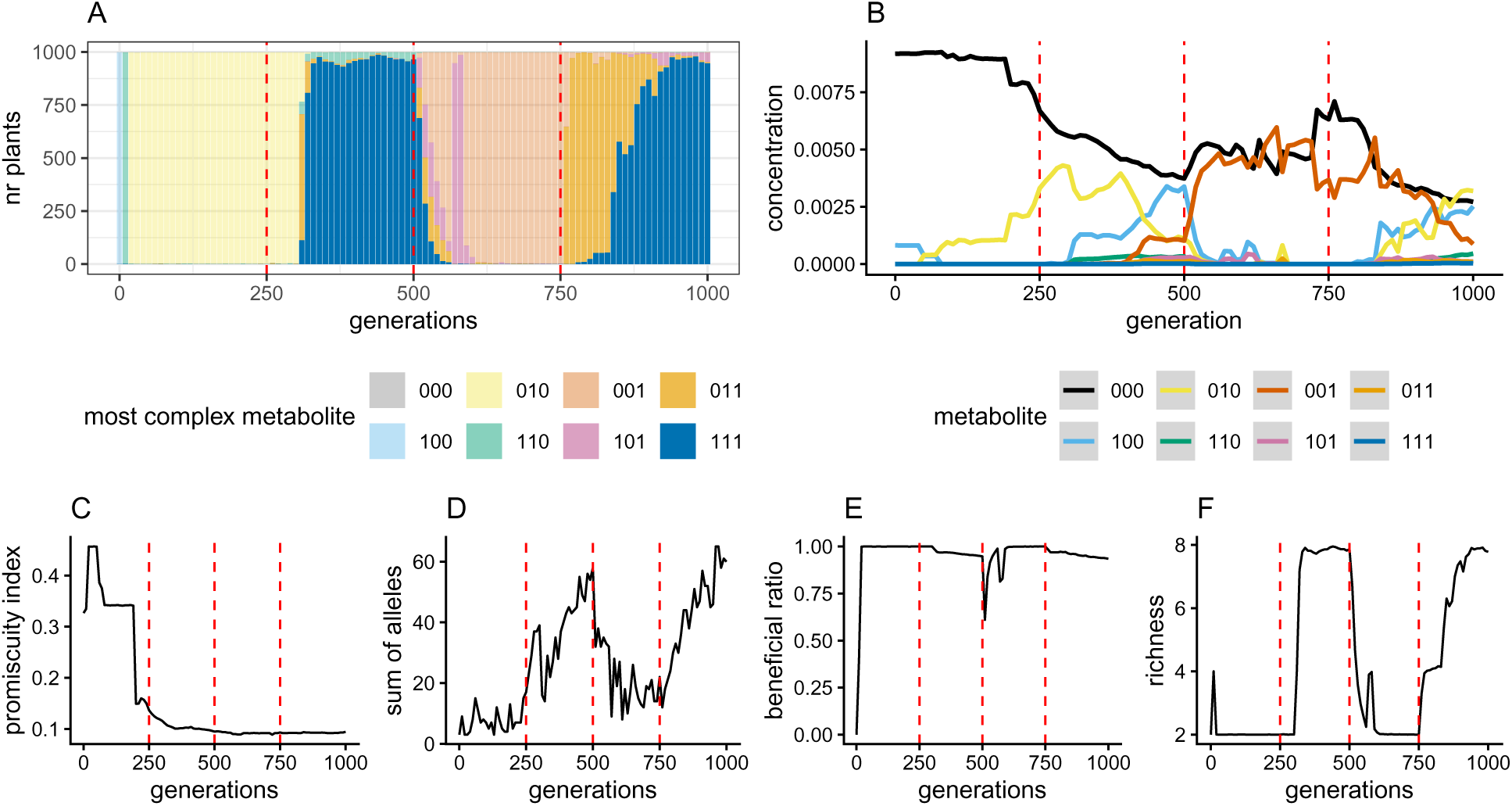
Output of one typical replicate of a simulation. In this run, enzyme costs are high (10), 5 metabolites are beneficial in phase effect scenario 1, and the alleles in the first generation code for the same enzyme function. A) the metabolite composition (chemotypes) of the population through time. B) the mean metabolic concentrations through time. C) the mean promiscuity index through time. D) the sum of alleles across loci in the population (SoA) through time. E) the mean ratio of beneficial metabolites to total metabolites through time. F) the mean richness of derived metabolites through time. The red lines indicate phase changes.

The robustness analysis (Morris, 1991; Campolongo *et al*., 2007; Imron *et al*., 2012) found that the most influential parameters were the number of beneficial metabolites, followed by the mutation rate and cost of enzyme production (Supplemental table S2.1). The number of beneficial metabolites and the cost of enzyme production are central to the hypotheses we want to test, and higher mutation rates make evolution take place faster. We conclude from this that the simulations we present here are a good representation of general model behavior.

As explained above, we run a set of simulations with three possible modifications, resulting in up to seven derived metabolites, to test the assumptions of both the screening and interaction diversity hypothesis. We also run a set of simulations with ten possible modifications, resulting in up to 1023 derived metabolites, to test the assumption of the screening hypothesis that very few metabolites of the high number of possible metabolites are beneficial. In the following sections, we will discuss the results for each of the four measurements we took to evaluate the predictions of the hypotheses.

Before that, we would like to point out a striking result we were not looking for: some simulations have no genes remaining after 1000 timesteps (Figures S4.1, S4.2). This is more common in simulations with fewer beneficial metabolites, higher enzyme costs, and simulations that start with genes that all code for enzymes with identical functions, than in those with more beneficial metabolites, lower enzyme costs, and those simulations that start with genes with different functions. All genes being selected away is an outcome that matches neither of the two hypotheses presented here. However, it is an outcome that is more common for simulation scenarios that match the assumptions of the screening hypothesis.

### 3.1 Promiscuity of enzymes

In the three-modification simulations, enzymes get less promiscuous and therefore more specific with time (Figure 4 A). This matches the predictions of the interaction diversity hypothesis (Table 1). In some scenarios with low enzyme cost, in later phases where only metabolites with at least two modifications are beneficial, the downward trend may halt for as long as the phase lasts (For example in scenario 1 phase 3, supplemental figures **??**S3.1fig:S4.1), S3.1n few metabolites are beneficial, the development of promiscuity more resembles a random walk in some simulations (Supplemental figures S6.1, S6.2). This matches the screening hypothesis (Table 1). However, under these circumstances, the more common result is for the population to evolve to not have enzymes and not produce any derived metabolites at all (Supplemental figure S4.1, which is why the lines end before 1000 generations are reached in most cases in supplemental figures S6.1, S6.2). This is similar in the ten-modification simulations. Here, promiscuity follows what amounts to a random walk in all simulations, and most simulations evolve to not produce any derived metabolites (Figures 5, S6.3, S6.4).

**Figure 4:**
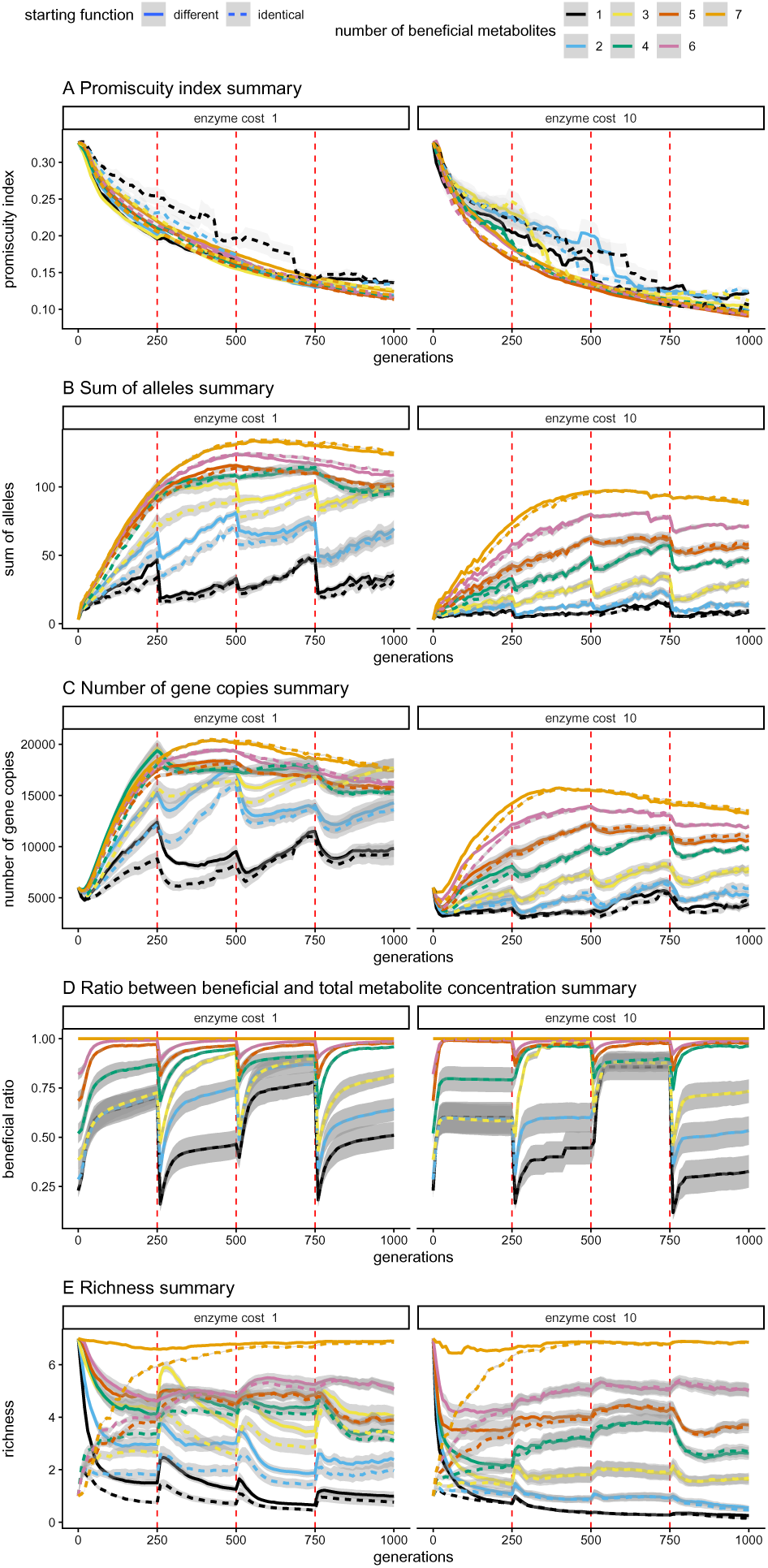
Summary figures of different diversity measures of the simulations with three modifications. Vertical dashed red lines in all graphs indicate phase changes. The other lines represent the mean value of a metric for all simulations split out by starting function, number of beneficial metabolites, and enzyme cost. The bands represent standard errors. Undefined values were left out of the analysis. A) the mean promiscuity index of the enzymes. High values indicate promiscuity, low values specialization. B) the mean SoA (sum of alleles across loci). C) the mean NoG (total number of gene copies across loci). D) the ratio of beneficial metabolites. E) the richness of derived metabolites.

**Figure 5:**
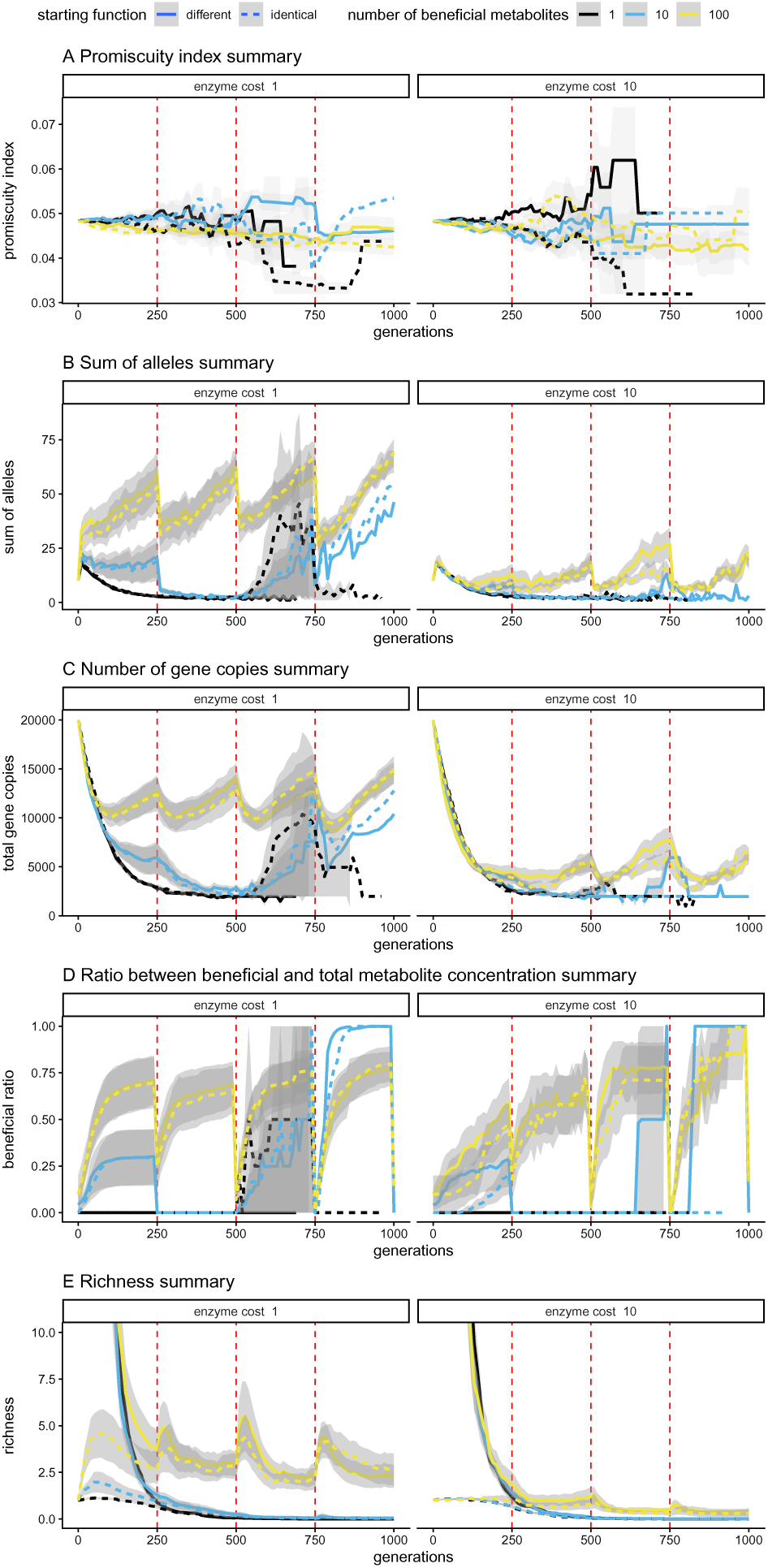
Summary figures of different diversity measures of the simulations with ten modifications. Vertical dashed red lines in all graphs indicate phase changes. The other lines represent the mean value of a metric for all simulations split out by starting function, number of beneficial metabolites, and enzyme cost. The bands represent standard errors. Undefined values were left out of the analysis. A) the mean promiscuity index of the enzymes. High values indicate promiscuity, low values specialization. B) the mean SoA (sum of alleles across loci). C) the mean NoG (total number of gene copies across loci). D) the ratio of beneficial metabolites. E) the richness of derived metabolites.

### 3.2 Selection/drift

The development of the SoA (sum of alleles across loci) and NoG (total number of gene copies across loci) is quite different between scenarios (Figure 4 B, C, 5 B, C). There are, in general, both fewer gene copies and alleles present in scenarios with higher enzyme costs, as well as in scenarios where fewer metabolites are beneficial, both when there are 3 and 10 modifications possible. At times, this leads to all genes being selected away (Supplement 4). This matches the prediction of selective effects from the interaction diversity hypothesis (Table 1).

In the shapes of the graph of the SoA and NoG (Figure 4 B, C), we see two kinds of developments in scenarios with three possible modifications, depending mostly on the number of beneficial metabolites. First, when there are many beneficial metabolites, the SoA gradually increases, then decreases slightly, without a clear effect of phase changes. The NoG shows a similar pattern, albeit with a small initial decrease. The increase in the SoA also continues longer than the increase in NoG. This indicates a simultaneous selection against the initial alleles and a selection for new alleles: adaptive radiation to fill the fitness space.

Second, in scenarios with fewer beneficial metabolites, what happens depends on the phase scenario. In these scenarios, the SoA and NoG follow the same pattern in the first phase if at least one metabolite with only one modification is beneficial. If enzyme costs are low, and all three metabolites with two modifications are beneficial, the SoA rises even higher than when all are beneficial, and the NoG does so even when not all two-modification metabolites are beneficial. This shows an adaptive radiation, and selection for the spread of genes in these circumstances. At high enzyme costs however, both SoA and NoG are low in these circumstances, showing strong selection to reduce enzyme costs (Figure S7.1, S7.2, S7.3, S7.4, S3.1).

This peaking above the SoA and/or NoG of the simulations where all derived metabolites are beneficial also sometimes occurs in later phases. For the SoA, this reliably happens when enzyme costs are low, and only metabolites with more than one modification are beneficial (Supplemental figures S7.1, S3.1). In these situations, both allelic variation and the sheer amount of enzyme is increasing rapidly. This results in high metabolite richness (Supplement 9, Figure S9.1).

After phase changes, the SoA abruptly drops when the modifications required to produce beneficial metabolites change (Figure S7.1, S7.2, S3.1). This gives the summary SoA of these scenarios a jagged shape. The NoG decreases more gradually after phase switches (Supplemental figures 4 C, S7.3, S7.4). This pattern, where the SoA starts increasing again while the NoG is still decreasing, points towards selection against previously beneficial alleles, with a few common ones needing longer to disappear than numerous rare ones, while new alleles with beneficial properties spread. Any resulting chemodiversity therefore is a transient reflection of past adaptations, rather than a current adaptation. Both the SoA and NoG may stay low for the rest of the phase if the beneficial metabolites are complex and/or enzyme costs are high (Supplemental figures S7.1, S7.2, S7.3, S7.4, S3.1)

In some simulations, the SoA may rise above that of the simulations with 7 beneficial metabolites, despite that a metabolite with only one modification is beneficial, such as in scenario 2 phase 3 with 2 beneficial metabolites. When we look at the NoG, we see that in this simulation, the NoG already rose above the simulations with 7 beneficial metabolites in the second phase, where only metabolites with at least 2 modifications are beneficial (Supplemental figures S7.1, S7.3, S3.1). We see a similar high NoG without a high SoA in phase 1 of scenarios 2 and 6 with 3 beneficial metabolites, phase 2 of scenario 3 with 2 beneficial metabolites, and phase 4 of scenario 9 with 2 beneficial metabolites (Supplemental figures S7.3, S3.1). A higher NoG without a higher SoG means that a small number of alleles is spreading rapidly, indicating selection for particular alleles.

In the ten-modification simulations with 1 or 10 beneficial metabolites, the SoA initially increases, then the SoA and NoG both decrease and stay low in almost all phase scenarios, which is reflected in the summary figure (Figure 5, Supplemental figures S7.5, S7.6, S7.7, S7.8). A partial explanation for this is that the plants start with a higher NoG than in the three-modification simulations. The simulations that still have enzymes by the end of the simulations show an increase to a similar range of NoG as in the three-modification simulations after the initial decrease (Supplemental figures S7.3, S7.4, S7.7, S7.8). In simulations with 10 or 1 beneficial metabolites this high SoA and NoG is achieved only in phases where there is at least one beneficial metabolite that only requires a single modification, and the simulation still produces enzymes (Supplemental figures S7.5, S7.6, S7.7, S7.8). For the simulations with 1 beneficial metabolite, this is only in phase 3 of scenario 3, at low enzyme cost (Supplemental figures S3.2, S7.5, S7.7). The fact that most simulations with 1 or 10 beneficial metabolites do not have genes by the end of the simulation, makes these single simulations stand out in the summary figure (Figure 5B, C). In the simulations with 100 beneficial metabolites, the SoA frequently rises higher than at the start of the simulation (Supplemental figures S7.5, S7.6). The NoG only does so at low enzyme cost (Supplemental figures S7.7, S7.8). In these simulations, most phases have at least one beneficial metabolite that only has one modification (Supplemental figure S3.2). In most phases then, there is an initial drop in SoA and NoG, after which there is a radiation, leading to a similar jagged pattern as in the three-modification simulations with few beneficial metabolites (Figures 4B, C, 5B, C, Supplemental figures S7.5, S7.6, S7.7, S7.8). However, when there is no beneficial metabolite with only one modification, the SoA and NoG frequently stay low after selection, particularly when enzyme costs are high. All these patterns indicate selection taking place, and therefore match the interaction diversity hypothesis (Table 1).

### 3.3 Match with beneficial metabolites

In summary over the different scenarios, the ratio between positive and neutral metabolite concentrations tends to increase over time within a phase (Figures 4 D, 5 D). There is a profound effect of phase changes. The ratio dips after phase changes and then rises, usually plateauing towards the end of a phase. In the summary (Figures 4 D, 5 D), this leads to high ratios for simulations with higher numbers of beneficial metabolites, but when we look at the individual scenarios (Figures S8.1, S8.2, S8.3, S8.4), we see this is mostly the effect of averaging.

In most scenarios, in most phases, the ratio is either close to 1 or 0 at the end of a phase (Supplemental figures S8.1, S8.2, S8.3, S8.4). Ratios are generally close to 1 if there is at least one beneficial metabolite with only one modification, and close to 0 or intermediate when there is no such metabolite. When it is intermediate, the ratio does usually increase with time, including in the phases and simulations with unusually high SoA and NoG. In simulations with higher enzyme costs, selection happens faster, but the ratio is also closer to 0 when in their equivalent low-enzyme cost simulations the ratio is more intermediate. This movement towards a high ratio of beneficial metabolites matches the predictions of the interaction diversity hypothesis (Table 1). In the simulations with 10 modifications, the phases with ratios close to 1 are those which have a beneficial metabolite that requires only 1 modification, and the phases with ratios at 0 are those which do not have such a beneficial metabolite (Figures S8.3, S8.4, S3.2). When 100 metabolites are beneficial, the ratio reaches a more intermediate level when only more complex metabolites are beneficial and enzyme costs are low. The result is that the ratio is highly scenario-dependent, making the summary appear more intermediate than that values actually are (Figure 5 D).

### 3.4 Metabolite richness

Summarized over multiple phase scenarios, richness is lower when enzymes are costly, and higher when there are more beneficial metabolites (Figure 4 E, 5 E). This matches the prediction of the interaction diversity hypothesis that richness depends on the number of useful metabolites (Table 1). Richness is the only metric for which the initial functions of the enzymes have an obvious effect, though this effect becomes smaller over time.

However, for specific scenarios, richness depends not only on the number of beneficial metabolites, but also on their complexity. In the three-modification simulations, for high enzyme cost, richness is driven by the simple metabolites. The richness almost always corresponds to the richness expected when all those metabolites that are beneficial and have one modification are made. As a consequence, the metabolites with multiple of these modifications are also made, whether they are beneficial or not (Supplemental figures S9.2, S3.1). For low enzyme cost, we see the same general pattern. However, different from the scenarios with high enzyme cost, when only more complex metabolites are beneficial, there can be a much higher richness than when some single-modification metabolites are also beneficial (see for example the scenarios with 4 vs 5 beneficial metabolites in phase 3 of scenario 1 in Supplemental figures S3.1, S9.1). Simulations with lower numbers of beneficial metabolites, and the 10-modification simulations with 100 beneficial metabolites show richness spikes at the start of the phase (Figure S9.1, S9.2, S9.3, S9.4). When enzyme cost is higher, these spikes are lower, disappearing in the summary (Figure 4 E, 5 E). The spikes reflect a temporarily increased diversity resulting from a switch from the previous set of beneficial metabolites to the current one. In the ten-modification model, richness is generally low. This selection matches the interaction diversity hypothesis (Table 1).

## 4 Discussion

We will now discuss the implications of the above for the screening and interaction diversity hypotheses. Additionally, we will discuss further insights from these simulations, the limitations of the model, how it can be expanded, and its implications for broader chemodiversity research.

In the results of the three-modification simulations (Figure 4), we see that usually the enzymes become more specific very quickly, usually without any effect of phase changes, matching the interaction diversity hypothesis. The only exceptions are found when there are sudden increases in SoA and NoG in later phases of simulations where, in that phase, only metabolites with more than one mutation are beneficial. In those simulations, the richness is also higher than the number of beneficial metabolites, and the ratio of beneficial metabolites is lower, all matching the expectations of the screening hypothesis. However, a central prediction of the screening hypothesis is that a large chemodiversity comes about through a low number of enzymes. The much higher SoA and NoG at these moments in these simulations is directly opposite to this prediction. Additionally, for some simulations with few beneficial metabolites, as well as in the ten-modification simulations, there is neither selection for specialization nor promiscuity. This could be interpreted to support the screening hypothesis. However, the simulations with no decrease in promiscuity almost always are also simulations that lose all enzymes, and therefore, metabolites, as can be seen from these lines ending before 1000 generations are reached. If there is selection against producing any enzyme, the promiscuity or specialization of those enzymes does not matter. Additionally, in the 10-modification simulations where 1000 generations are reached, the metabolite richness is a fraction of all the metabolites that could exist. This means that the values of *k_cat_/K_M_* for most substrates do not affect fitness at all, so that selection cannot act on them. Under these circumstances, there is an absence of selection on promiscuity, but not for the reasons why it is expected by the screening hypothesis.

We do see a strong selective response to phase changes in the sum of alleles across loci in the population (SoA), total number of gene copies across loci (NoG), and richness when there are few beneficial metabolites, particularly when enzyme costs are low. This lines up well with the predictions of the interaction diversity hypothesis. The ratio of beneficial metabolites to neutral metabolites is generally high regardless of the number of beneficial metabolites, which also matches the predictions of the interaction diversity hypothesis. Our findings complement those of Wittmann & Bräutigam (2025) and Whitehead *et al*. (2021). Both Wittmann & Bräutigam (2025) and us find a higher average richness per plant when more metabolites are beneficial, and Whitehead *et al*. (2021) finds that mixtures of higher richness affect more herbivores. This all supports the interaction diversity hypothesis. Both Wittmann & Bräutigam (2025) and our study also find that plants are more resilient to changes in which metabolites are beneficial when they produce numerous metabolites before the change takes place. However, like Wittmann & Bräutigam (2025), we note that this chemodiversity before the phase change comes about only when this chemodiversity is beneficial at that time. Therefore, it is a side-effect of interaction diversity rather than the selection for a flexible metabolism predicted by the screening hypothesis. Finally, Wittmann & Bräutigam (2025) were limited in their ability to test the screening hypothesis for lack of modeling of the metabolic pathway, and Whitehead *et al*. (2021) by being limited to commercially available metabolites. Without these limitations, we clearly show that, even when few metabolites are beneficial, and there are numerous potential metabolites, there is selection to produce a high ratio of beneficial metabolites, not to create a metabolic system that makes numerous compounds.

Both their and our results support the interaction diversity hypothesis to a far greater extent than the screening hypothesis (Table 1). The evidence for the screening hypothesis is rather weak for an evolutionary mechanism that is supposed to apply in a wide set of scenarios. Firn and Jones’ observation that promis-cuous enzymes can create a great chemodiversity when enzymes are expensive (Jones & Firn, 1991) is still useful, but the end result will still be focused on the production of useful metabolites, not a great diversity of metabolites in general. We do hold space for the possibility that modeling the metabolism in some other way, or introducing a different fitness function (for example by calculating fitness cost per catalyzed reaction, introducing a small fitness benefit of ‘non-beneficial’ metabolites, or diminishing returns of increased concentrations of beneficial metabolites) could lead to a different outcome.

Although we focused on comparing the screening and interaction diversity hypotheses, we also found some other interesting patterns. Different runs of the same parameter settings can lead to different chemotypes (Supplement 5). When multiple different pathways produce beneficial metabolites, investing in multiple of these pathways may convey more additional fitness costs than benefits. This, through stochasticity and positive feedback loops, can result in alternative stable chemotypes. There is no spatial element to the model, but alternative stable chemotypes could explain why spatial disconnected populations may develop different chemodiversity patterns even without local differences in evolutionary pressures. Rahimova *et al*. (2024) for example found a geographic effect on the distribution of sets of biosynthetically linked monoterpenoids in Germany, with more distant populations being more dissimilar along both a north-south and east-west gradient, even when accounting for the differences in chemotype stemming from soil type differences. They suggest that this may be caused by differences in the soil type more subtle than they assessed in their study. Our model results suggest the additional possibility of alternate stable strategies as an explanation for this geographic difference in chemotypes.

Accidental findings like this highlight the two ways the model could be further developed. On the one hand, it could be made more specific to particular species and populations, to test hypotheses about their chemodiversity evolution. For this, good information on the metabolism and interactions of the focal plants are paramount. Alternatively, the model could be kept general, but be modified to test additional hypotheses. For example, while we model the benefit of a metabolite linearly by multiplying its fitness value with its concentration, we would need a fitness function that allows metabolites to increase each others effectiveness to test the synergy hypothesis (Richards *et al*., 2016). Other nonlinear fitness functions could convey diminishing returns with increased metabolite concentrations. One could also include a spatial component to test how associational effects affect chemodiversity evolution (Hambäck *et al*., 2014). Additionally, while the fitness benefits of metabolites implicitly modeled herbivore pressures, herbivores could be modeled explicitly. These herbivores would cause damage, and be attracted or repelled by certain metabolites, similar to previous models (Speed *et al*., 2015; Wittmann & Bräutigam, 2025). All of these options introduce a lot more added complexity, but they are worthwhile endeavors nonetheless.

With this model, we aim to add to the discussion about the use of different chemodiversity hypotheses. In our study, the patterns predicted by the screening hypothesis were not found in most scenarios in our model. Those of the interaction diversity hypothesis were. We suggest the field to look towards interaction diversity rather than screening as an explanation of patterns of chemodiversity.

## Supporting information

Simulation code and files

## 5 Acknowledgments

This project was funded by the German Research Foundation (DFG), projects WI 4544/2 as part of the Research Unit (RU) FOR 3000 Ecology and Evolution of Intraspecific Chemodiversity of Plants. We thank the members of the theoretical biology group in Bielefeld and the members of FOR 3000 for their invaluable contributions in numerous discussions. This work was supported by the de.NBI Cloud within the German Network for Bioinformatics Infrastructure (de.NBI) and ELIXIR-DE (Forschungszentrum Jülich and Wde.NBI-001, W-de.NBI-004, W-de.NBI-008, W-de.NBI-010, W-de.NBI-013, W-de.NBI-014, W-de.NBI-016, W-de.NBI-022).

## 7 Author contributions

Both authors contributed substantially to the model design. Frans Thon programmed the model and analyzed output data. Frans Thon wrote the first draft of the manuscript. Both authors contributed substantially to rewrites and revisions.

## 8 Data availability

Model code and parameter files to recreate model output, as well as code to recreate the analysis can be found in the supplemental .zip file

## Brief supplement captions

**Supplement 1:** ODD model description; detailed description of the model according to the ODD protocol

**Supplement 2:** Robustness analysis

**Supplement 3:** Phase legend; graphic legend of the phase scenarios

**Supplement 4:** Allele presence and maximum; figures describing the relationship between parameters and all genes dying out/the maximum number of loci being hit

**Supplement 5:** Chemotype diversity through time; examples of the chemotype development through time

**Supplement 6:** Promiscuity: Figure 4a and 5a split out by phase scenario and enzyme cost

**Supplement 7:** Selection/drift: Figures 4b, 4c, 5b, and 5c split out by phase scenario and enzyme cost

**Supplement 8:** Match with beneficial metabolites: Figure 4d and 5d split out by phase scenario and enzyme cost

**Supplement 9:** Richness: Figure 4e and 5e split out by phase scenario and enzyme cost

