## Supplement 1 for "A metabolic model to investigate the evolution of chemodiversity"

### S1 Metabolic individual-based chemodiversity model

This model description follows the ODD (Overview, Design concepts, Details) protocol for describing individual- and agent-based models (Grimm *et al.*, 2006), as updated by Grimm *et al.* (2020).

#### S1.1 Purpose and patterns

The purpose of this model is to describe the evolution of the chemodiversity of a population of plants over several generations. We model a simplified metabolic pathway that determines what the chemotype of a plant is from a gene level up to the metabolites.

We want to see whether more complex chemodiversity can develop from a population with no or little initial chemodiversity, and whether chemodiversity can be maintained. We also want to see if we find the patterns of selection and enzyme kinetics expected from the screening (Jones & Firn, 1991) and interaction diversity hypothesis (Whitehead *et al.*, 2021). For the screening hypothesis, this would mean a large diversity of metabolites that changes through time, but not in response to changes in the fitness situation. For the interaction diversity hypothesis, this would mean a diversity of metabolites specifically adapted to the current fitness situation (Table 1)

The hypotheses tested here make assumptions about how the enzyme pathway constrains what patterns of chemodiversity are possible. Therefore, modeling metabolites directly, the way other models do it, would be insufficient. Instead, we model the genes that code for enzymes and their evolution, and how these enzymes bring about the metabolites.

#### S1.2 Entities, state variables, and scales

##### S1.2.1 Entities and state variables

We describe the variables of the different model entities (the environment, plants, and genes) here. These variables are described in what they represent (their meaning), their units, and their range if applicable.

The **environment** consists of variables which contain important information about the simulation that can be called on by any other entity, such as the population size.

The vector of **metabolite codes** is a vector of vectors saved in the environment. Every entry in the vector corresponds to a possible metabolite in the system, and the vector is as long as the number of possible metabolites.

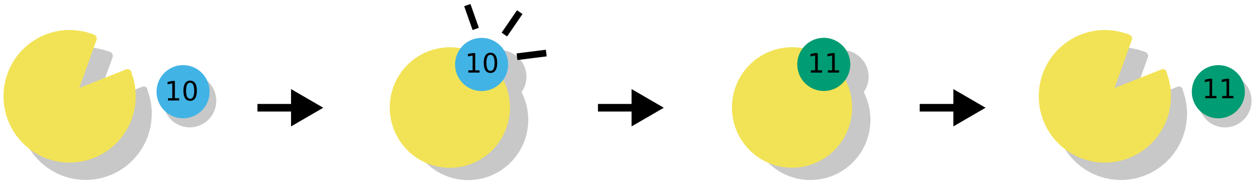


Figure S1.1: Graphical explanation of the metabolite codes. The enzyme (in yellow) modifying a metabolite (in blue) is modeled as an enzyme changing a particular position in the metabolite code from 0 to 1. Here, an enzyme modifies a metabolite that already had the modification in position 1 to also have the modification in position 2.

A single metabolite code consists of a vector of ones and zeroes as long as the number of possible modifications. An enzyme can modify a metabolite by, for example, adding or removing a group. This is implemented as an enzyme irreversibly changing a single position in the metabolite code from 0 to 1. So a '0' in a metabolite code indicates that in this metabolite, the modification in that position has not taken place. a '1' indicates that is has. In this way, to use the example of a system with two possible modifications, the vector of metabolite codes has four entries: '00' for the unmodified metabolite 0, '10' for the metabolite 1 where an enzyme has made the first modification, '01' for the metabolite 2 with the second modification, and '11' for the metabolite 3 with both modifications (Figure S1.1).

The **population** is a collective that represents a population of **plants**. Its state variable is a dynamic vector of **plant** entities.

An individual **plant** is an entity in the **population**. It contains as its state variable a vector of gene pairs containing **gene** entities (the alleles in this 'genome' that are not empty correspond to the enzymes $E$) with variables from which the metabolism (described in the next subsection) is derived, and which is used in reproduction. This metabolism determines the plant's metabolite concentrations $M$ and therefore its fitness value. The amount of enzyme produced by an allele is only determined by the concentration coded in the allele, independent of whether it is present as a homozygote or heterozygote.

The **gene** entity represents an enzyme-coding gene of a plant.

##### S1.2.2 Scales

The model consists of a population of plants, with each plant having its own genome and metabolism.

The spatial interactions between the **plants** in the **population** are not included in any way in the model.

The population size is determined by an input value and is not changed during the course of reproduction. The plants reproduce in discrete generations, that is, all plants die and are replaced by a new population of the same size.

The ordinary differential equations (ODE) of the **metabolism** (see subsection S1.3.2) represent the development of metabolites in mM/s, for 100 seconds (time steps), with the final concentration in mM

Table S1.1: State variables of a gene entity

| Variable name | Variable type and units | Meaning |
|---|---|---|
| does exist | boolean | value that determines whether a gene exists |
| enzyme concentration | double $E_e$ | Concentration: The enzyme concentration produced by the gene. |
| enzyme function | unitless integer | position in the vector of metabolite codes that the enzyme modifies |
| vector $k_{cat}$ | vector of doubles in $s^{-1}$ | Catalyzing rates $k_{cat}$ for each metabolite |
| vector $K_M$ | vector of doubles in $mM$ | Michaelis constants $K_M$ for each metabolite |

reported to the output.

#### S1.3 Process overview and scheduling

##### S1.3.1 Summary

The model covers the evolution of a population of plants over many generations. It is structured in the following way. First, the **initialization** process is run, setting up the simulation by creating the initial genome and running the **metabolism** and **fitness calculation**. Then, four processes take place in every generation, in order: **reproduction**, in which the genomes of the last generation are used to produce genomes for the new generation, which replaces it, **mutation**, in which the genes of these new plants have a chance to mutate in various ways, **metabolism**, in which the genes in each individual are used to produce enzymes and run the metabolism submodel to calculate the metabolite concentrations of each plant, and **fitness calculation**, in which the fitness of each individual is calculated from their metabolites. Finally, the **recording** process is run in some but not all generations, as described in **observation**. In this process, the state of the model is recorded.

At the end of the run, **recording** takes place one last time.

##### S1.3.2 Metabolism

Metabolism follows after either initialization or mutation.

Every plant metabolism is ran in order.

The metabolism is modeled as a system of ordinary differential equations (ODE) that are automatically generated for each plant. It is part of the **updating** step of the model.

These ODEs take the form of Michaelis-Menten equations:

$$\text{Substrate} \xrightarrow{\text{E}_e} \text{Product} \tag{S1.1}$$

$$\frac{dM_0}{dt} = \gamma - \sum_{e=1}^{\eta} \left( \frac{k_{cat_{e0}} \cdot M_0}{K_{M,e0} + M_0} \cdot E_e \right) - M_0 \cdot \epsilon \tag{S1.2}$$

$$\frac{dM_m}{dt} = \sum_{e=1}^{\eta} \mathbb{1}_{\{code_{me}=1\}} \left( \frac{k_{cat_{eS_{em}}} \cdot S_{em}}{K_{M,eS_{em}} + S_{em}} \cdot E_e \right) - \sum_{e=1}^{\eta} \mathbb{1}_{\{code_{me}=0\}} \left( \frac{k_{cat_{em}} \cdot M_m}{K_{M,em} + M_m} \cdot E_e \right) - M_m \cdot \epsilon \tag{S1.3}$$

with an equation $\frac{dM_0}{dt}$ for the change in concentration of the unmodified metabolite, and an equation $\frac{dM_m}{dt}$ for the change in every other metabolite concentration. Here, $E_e$ is the enzyme concentration of enzyme $e$ in the vector $E$, $M_m$ is the concentration of metabolite $m$, and $S_{em}$ is the concentration of the precursor substrate of $M_m$ before modification by enzyme $e$. $code_{em}$ is the value of the bit in the metabolite code of the metabolite $m$ in the position that the enzyme $e$ modifies (the function of the enzyme), which indicates whether the metabolite is a substrate (if 0) or product (if 1) of the enzyme in question. The first sum of the equation $\frac{dM_m}{dt}$ therefore describes how much of the metabolite is formed, and the second sum describes how much of the metabolite is metabolized into a different metabolite. $\gamma$ is the influx (production) of new unmodified metabolite by the metabolism, $\epsilon$ the constant efflux (through evaporation, breakdown etc.) of all metabolites. The values of kinetic parameters $K_M$ (Michaelis constant) and $k_{cat}$ (catalyzing rate) are specific to an enzyme-metabolite pair. They are taken from the **gene** entities of the plant. The initial value of $M_0$ is taken from the **input**, and the other initial values of $M$ are set to 0 at the start of the ODE.

Solving the ODE is resource-intensive. For this reason, whenver a plant does not have enzymes with every possible function, the parts of the ODE which reflect those functions (which would resolve to 0) are removed to conserve resources.

This system of ODEs is then solved using a Runge-Kutta-4 stepper from the boost ODEINT library run for a time length of 100 units. The step size for the ODE is 1.0, and the results are checked for whether they produce NaNs or negative numbers. If there are any, the ODE is re-run at a step size of step size/10, until a minimum step size of 0.01. If there are still any NaNs or negative values at that point, the values of the ODE are set to 0 and the simulation puts out an error before continuing (This should not happen with the parameter values used in the study).

A value in the **input** can be set to TRUE to output the time series of the ODE. In this case, the metabolite concentrations in every time step are written to a comma separated file while the ODE is run. The final state of the ODE is returned to the plant as the final concentration of the metabolites.

Metabolism is followed by fitness calculation.

##### S1.3.3 Fitness calculation

The fitness of a plant is calculated as follows. For every gene, the fitness cost of the enzyme it produces is calculated by multiplying the enzyme concentration $E_e$ from the vector $E$ by the enzyme cost $C_{\text{enzyme}}$ from the **input** object. Then the fitness effect of each metabolite at the timepoint $F_{m_t}$ is taken from the **input** object or the randomly generated values, and multiplied by the concentration of that metabolite $M_m$ in the metabolite concentration vector $\nu$. Together, these determine the fitness of the plant:

$$Plantfitness = \sum_{e=1}^{\eta} E_e \cdot C_{\text{enzyme}} + \sum_{m=1}^{\nu} M_m \cdot F_{m_t}. \tag{S1.4}$$

These plant fitnesses are stored as a vector on the population level where the position of the fitness value in the fitness vector corresponds to the position of the corresponding plant in the population vector.

##### S1.3.4 Reproduction

Reproduction uses a discrete weighed lottery with replacement to determine the parent plants of each plant in the new population. The fitness value (see equation S1.4) of each plant in the current population is used as its weight, so that plants with a higher fitness are more likely to reproduce. For each plant in the new population, two parents are sampled from the population. Then, as we assume free recombination between loci, one allele from each parent is randomly selected at each locus. This can be an empty allele. Once alleles have been assigned to all loci, the genes that exist in the plant are copied into a vector $E$ for easy referencing by the metabolism. This process is followed for every plant in the new population, so that the new plants have genomes, but nothing else is initialized. The new vector of plants replaces the old vector of plants in the population. Reproduction is followed by mutation.

##### S1.3.5 Mutation

The state variables of every extant gene copy in every plant may be mutated. Mutations happen randomly with a certain probability as set by values in the **input** object. There are separate probabilities of mutation for duplication and gene loss, and all other mutations happen at an equal rate. Depending on the mutation, there might be a second random factor to determine the scope of the mutation. In these cases, this is explained where the mutation is described below. After every mutation, the details of the mutation and new gene are written to output files. In all cases except for gene loss, the mutated gene is also assigned a new unique ID.

We set a maximum number of loci in the **input** file. The vector of gene pairs is not allowed to grow longer than this value, to prevent excessively long simulations due to very large numbers of enzymes. If a gene is to be duplicated, it is first checked whether the genome is smaller than the maximum number of loci. If it is, a new gene pair is created for that plant, and the gene is copied into one of its alleles. The other allele on this new locus is marked non-existent. Gene pairs of alleles marked as non-existent are added to the genomes of all other plants. If however the maximum number of loci has been reached, a warning is produced instead and written to an output file before the simulation continues without the gene being duplicated.

If a gene is lost, it is marked as non-existent.

Gene duplication and loss happen before the other mutations, as a duplicated gene should not inherit mutations that happen after duplication from its ancestor, and mutating a non-existent gene is pointless.

If the enzyme concentration produced by the gene is mutated, a random value is picked from a continuous uniform distribution centered around 0, with a minimum and maximum determined by a variable defined in the input **input**. This value is then added to the existing enzyme concentration variable of the gene, resulting in a new gene producing a new enzyme concentration.

If the enzyme function of the enzyme the gene codes for is mutated, a random value is picked from an integer uniform distribution with a minimum value of 0 and a maximum value determined by the variable that sets the number of possible enzyme modifications from the **input**. This value is the new function of the enzyme the gene codes for.

If the values of $K_M$ are to be changed in a coding gene, random values are picked for every position in the vector (representing $K_M$ for each metabolite) from a lognormal distribution with a normal mean ($\mu$) and normal standard deviation ($\sigma$) determined by the input file. The values of $K_M$ are then replaced by these values.

The same procedure is followed when $k_{cat}$ is mutated. In both cases, $K_M/k_{cat}$ is evaluated to make sure it remains within the thermodynamically realistic frame of $10^{-9}$.

Once all genes have completed mutation, the metabolism follows.

##### S1.3.6 Cleanup

Whenever the length of the gene pair vector is more than two-thirds the maximum number of alleles at the end of **fitness calculation**, loci which are empty in every single plant in the population are removed from all genomes.

#### S1.4 Design concepts

##### S1.4.1 Basic concepts

The basic principle of this model is that it is meant to elucidate the circumstances under which the chemodiversity we see in plant populations can come about and be maintained. To this end, we model a population of plants and their evolution over multiple generations. In this way, we can test different hypotheses about the origin and maintenance of chemodiversity, such as the screening hypothesis (Jones & Firn, 1991) or interaction diversity hypothesis (Whitehead *et al.*, 2021). We can test these hypotheses by seeing whether the patterns show up that we would expect to see if the hypotheses were accurate.

We model a simplified metabolic pathway that determines what the chemotype of a plant is from a gene level up to the metabolites, which determine the fitness of the plant. In this way, we can see the connection between chemodiversity and evolution over many generations.

##### S1.4.2 Emergence

The emergent pattern of interest of the model is the genotypic and chemotypic diversity of the population over time. Starting from a single, simple genotype and chemotype, the system can evolve into one which has complex genetic and chemical diversity.

The results come about through random mutation of existing genetic diversity, followed by genetic drift, recombination, and selection during reproduction.

##### S1.4.3 Adaptation

During reproduction, plants with a higher fitness are likely to produce more offspring for the next generation than plants with a lower fitness. This selection is expected to cause adaptation to the scenario modeled in the model.

##### S1.4.4 Objectives

The objective measure in reproduction is the fitness value of each plant.

##### S1.4.5 Stochasticity

Stochasticity exists in the model in several different places.

1. Fitness effects of metabolites $F_{m_{\text{phase}}}$ may be set randomly at initialization. These fitness effects are drawn from a uniform distribution with a maximum and minimum value provided by the input and using a random number generator (RNG) seed provided by the input. Alternatively, fitness effects of metabolites can be set manually through the input file.

2. Random **mutations** occur at **reproduction**, introducing mutated genes into the population. These mutations are drawn from uniform or lognormal distributions, with their boundaries set by the input and using a RNG seed provided by the input. The initial kinetic values $k_{cat}$ and $K_M$ can be randomly drawn from the same distribution as during mutation, or set manually through the input file.

3. **Reproduction** happens through a weighed lottery with replacement with the fitness values of the parent plants as the weights. This lottery uses a RNG seed provided by the input.

##### S1.4.6 Observation

The model has several outputs.

1. In the input an option to output every metabolism time series can be set to 'true'. If it is, the metabolism time series is written to a comma-separated file as the time series is created. All such time series created in the model are written to separate files. These time series consist of the time step within the time series and the metabolite concentrations $M$.

2. Whenever a non-loss mutation takes place, the state variables of the new allele are noted down in a comma-separated file, including all of its ancestors.

The following outputs are created after initialization of the population, and then at intervals of as many generations as specified by the input. This happens after reproduction and updating are complete in the previous generation, but before reproduction happens to produce the new generation.

4. The fitness value (see equation S1.4) of each plant.

5. If the option "outputMetabolites" is set to TRUE: The concentrations of each metabolite at the end of the ODE time series of each plant.

6. The extant gene copies of each plant.

7. The ratio of beneficial metabolites in each plant.

8. The metabolite richness of each plant.

#### S1.5 Initialization

Table S1.2: Input parameters

| Variable name | Variable type and units | Meaning | Standard value |
|---|---|---|---|
| seed | unitless integer | random seed used to draw all random numbers in the simulation that are not drawn using their own seed | different for every simulation |

| | | | |
|---|---|---|---|
| population size | unitless integer | number of plants in the population | 1000 |
| number of generations | unitless integer | number of generations the simulation runs for | 1000 |
| number of gene pairs | unitless integer | number of initial gene pairs initialized | 3/10 |
| identical function | boolean | value that determines whether the initial alleles all produce enzymes with the same enzyme function or not | variable |
| base enzyme concentration | double in $mM$ | starting concentration of enzymes all starting alleles produce | 0.001 |
| gene loss odds | unitless double | chance in every timestep of every gene copy of being randomly lost | 0.01 |
| gene duplication odds | unitless double | chance in every timestep of every gene copy of being randomly duplicated | 0.0001 |
| maximum nr. of loci | unitless integer | cutoff of genome size after which gene duplications are discarded | 80 |
| mutation odds | unitless double | chance in every timestep of ever gene copy of being randomly mutated in its enzymatic variables | 0.0001 |
| maximum concentration | double in $mM$ | maximum concentration of enzymes | 0.1 |
| maximum concentration change | double in $mM$ | maximum change of the enzyme concentration from a mutation | 0.001 |
| $\mu$ | unitless double | normal mean of the lognormal distribution that values of $k_{cat}$ and $K_M$ are drawn from during mutation | -3.45 |
| $\sigma$ | unitless double | normal standard deviation of the lognormal distribution that values of $k_{cat}$ and $K_M$ are drawn from during mutation | 0.68 |

| | | | |
|---|---|---|---|
| number of reactions | unitless integer | number of changes to the metabolite that enzymes can catalyze | 3/10 |
| set initial kinetic variables | boolean | whether the initial kinetic variables are provided by the user or drawn from the lognormal distribution | false |
| initial K seed | unitless integer | random seed used to draw the initial values of the kinetic variables | 1 |
| initial $k_{cat}$ | vector of vectors of doubles in $s^{-1}$ | values of $k_{cat}$ for every substrate for every initial enzyme | N/A |
| initial $K_M$ | vector of vectors of doubles in $mM$ | values of $K_M$ for every substrate for every initial enzyme | N/A |
| initial metabolite concentration | double in $mM$ | concentration of the unmodified metabolite at the start of the ODE | 0.01 |
| influx | double in $mM$ | amount of unmodified metabolite added in each timestep in the ODE | 0.001 |
| efflux | unitless double | proportion of all metabolites removed every timestep of the ODE | 0.1 |
| ODE length | unitless double | number of timesteps in the ODE | 100 |
| ODE stepper | string | stepper used to solve the ODE | runge-kutta |
| smallest ODE step size | unitless double | smallest stepsize the ODE will try before giving up | 0.01 |
| fitness has effect | boolean | value that determines whether fitness affects reproduction | true |
| number of phases | unitless integer | number of phases | 4 |
| phase transition times | unitless integer vector | times at which phase switches take place | 250, 500, 750 |
| fitness cost per enzyme | unitless double | fitness cost per unit enzyme | 1, 10 |

| | | | |
|---|---|---|---|
| fitness is random | boolean | value that determines whether fitness is set by the user or randomly generated | true |
| fitness values | unitless double vector of vectors | fitness values for each metabolite in each phase | N/A |
| fitness seed | unitless integer | random seed that fitness values are drawn from for each metabolite in each generation | 0-3/0 |
| minimum fitness effect | unitless double | minimum value of randomly drawn fitness effects | 0 |
| maximum fitness effect | unitless double | maximum value of randomly drawn fitness effects | 500 |
| phase scenario seed | unitless integer | random seed that fitness values set randomly to 0 in the different phase scenarios are drawn from | 1-10 |
| number non-beneficial metabolites | unitless integer | number of metabolites randomly set to 0 | 0-6/1022,1013, 923 |
| output ODE | boolean | value that determines whether the ODE timeseries is written to a file | false |
| output Metabolites | boolean | value that determines whether the metabolite concentrations are written to a file | true/false |
| output intervals | unitless integer | number of generations between writing to output | 10 |

At initialization, a population of plants is created using parameters from the **input** (Table S1.2). A single plant is initialized with only the vector of gene pairs and the gene vector initialized.

A number of gene pairs as defined by the **input** is added to the genome, with each locus comprising two identical gene copies. Every allele is assigned a unique ID. It is initialized with the following state variables:

1. A boolean value that indicates it does exist.

2. The resulting enzyme concentration of the enzyme it produces.

3. An enzyme function for the enzyme it codes for (determined in the following way: The first allele will have the function of modifying the metabolite in the first position, with each next enzyme initialized modifying the next position, starting over at position 1 if the number of gene pairs is greater than the number of possible modifications)

4. Vectors of the kinetic values of the allele, $k_{cat}$ and $K_M$. Depending on an option in the **input** object, these values may be taken from the **input** object or randomly determined for each pair in the same way as during **mutation**.

   The fitness values of the metabolites are randomly generated based on the fitness seed (with as many set to 0 in each phase based on the number non-beneficial metabolites parameter), or set by the user.

After the creation of the genotype, the initial plant is copied into the population as many times as the population is big, so that the whole population consists of plants with this initial genotype. After that, **metabolism** and **fitness calculation** are run to create the phenotypes. Finally, the output files are created, after which initialization is finished.

#### S1.6 Data

When the kinetic variables are mutated, values are drawn from a lognormal distribution described by $\mu$ and $\sigma$ taken from a parameter file. In the paper, these values are $\mu = -3.45$ and $\sigma = 0.68$, and were chosen so that most of the values drawn from the distribution would be between 0.01 and 0.1 (mean 0.04, sd 0.03, mode 0.02). These values are informed by information on the kinetics ($K_M$ and $k_{cat}$) of the terpene pathway of *Salvia officinalis* (Croteau *et al.*, 1994; Ignea *et al.*, 2014; Kampranis *et al.*, 2007). The initial values of these kinetic variables can either be randomly drawn from this same distribution, or are set at a particular value taken from a parameter file at the start of the simulation.

The odds that any parameter of the gene is mutated is $10^{-4}$, based on per-locus mutation rates in previous evolutionary models (Østman *et al.*, 2012; Speed *et al.*, 2015; Yeaman *et al.*, 2018).
