## Supplement 2-10 for "A metabolic model to investigate the evolution of chemodiversity"

### S2 Robustness analysis

We performed a robustness analysis using the Morris method (Morris, 1991; Campolongo *et al.*, 2007; Imron *et al.*, 2012).

We sampled 50 random trajectories through parameter space over four levels (that is, the parameter range is divided into four bins for each parameter), and calculated the elementary effects of fourteen parameters on richness, promiscuity, SoA, and beneficial metabolite ratio in the last timestep of the simulation. As the parameter "identical initial allele function" is boolean, we let two levels be identical initial functions, and two levels different initial functions. This resulted in 15 trajectories where no change in this boolean value happened. Those trajectories were removed from the analysis for the elementary effects of identical initial allele function parameter.

We calculated the savage scores (Iman & Conover, 1987) of the estimated means of the absolute values of the elementary effects ($\mu*$) of all parameters for the four outputs, and summed the savage scores for each of the outputs for each parameter to make a ranking of the effects (Campolongo *et al.*, 2007) (Table S2.1). Savage scores are calculated as follows:

$$S_i = \sum_{j=i}^{n} 1/j, \tag{S2.5}$$

with $i$ the rank of the parameter for a particular output when parameters are ordered from highest to lowest $\mu*$, and $n$ the number of parameters.

We also plotted the $\mu*$ against the standard deviation $\sigma$ of the elementary effects of the parameters for each output, to show not only the simple rank of each parameter's elementary effects, but also the whether that difference in rank reflects a large or small difference between different parameters (Figure S2.1).

The most influential parameter is the number of non-beneficial metabolites (which equals the total number of derived metabolites minus the number of beneficial metabolites), followed by the mutation rate and enzyme cost (Table S2.1). The number of non-beneficial metabolites also has the highest $\mu*$ of all parameters for two out of four outputs (Figure S2.1). The effect strength of the parameters ranked next stands out less. The elementary effects of the mutation rate are still notably higher than those of enzyme cost, but after that, the effects of the middling parameters are more similar. It must be noted that the mode of the distribution the kinetic values are drawn from and the initial enzyme concentration are particularly influential when looking at the promiscuity output. This makes sense, as the mode very directly influences the likelihood of an enzyme being particularly promiscuous or specialized, and the initial enzyme concentration has a great influence on how much metabolite an enzyme produces in a different way from enzyme promiscuity. Together, they play

Table S2.1: Ranked table of parameters based on the sum of their savage scores

| Parameter | Score | Rank |
|---|---|---|
| Nr. of non-beneficial metabolites | 9.9 | 1 |
| Mutation rate | 7.2 | 2 |
| Enzyme cost | 5.4 | 3 |
| Population size | 4.8 | 4 |
| Initial enzyme concentration | 4.2 | 5 |
| Gene loss rate | 4.0 | 6 |
| Mode kinetic distribution | 3.8 | 7 |
| Maximum fitness effect | 3.7 | 8 |
| Duplication rate | 3.6 | 9 |
| Maximum number of loci | 2.8 | 10 |
| Mean kinetic distribution | 2.7 | 11 |
| Metabolite influx | 2.2 | 12 |
| Identical initial allele function | 0.9 | 13 |
| Maximum enzyme concentration change | 0.7 | 14 |

a large role in generating the diversity selection can act on, and the strength of this selection, respectively.

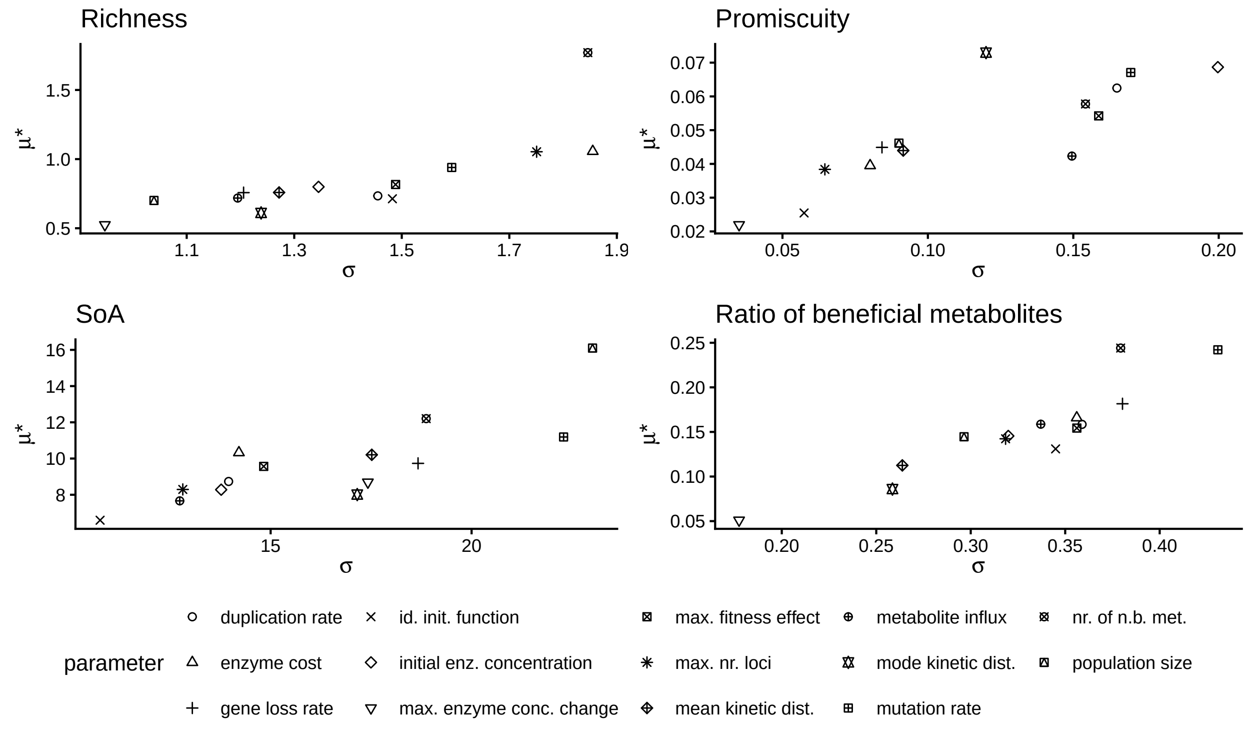


Figure S2.1: Plot of mean variance $\mu*$ and standard deviation $\sigma$ of the elementary effects of the parameters.

The relatively large effect of the number of beneficial metabolites, and the middling effect of the enzyme cost, gives us confidence that the effects we find of these parameters in this work are a robust representation of the model behaviour, rather than a very specific case dependent on very specific settings of the other parameters. The large effect of mutation rate, and the still substantial effect of the 'medium effect' cohort of parameters means that we select the values of these parameters with particular care.

### S3 Phase legend

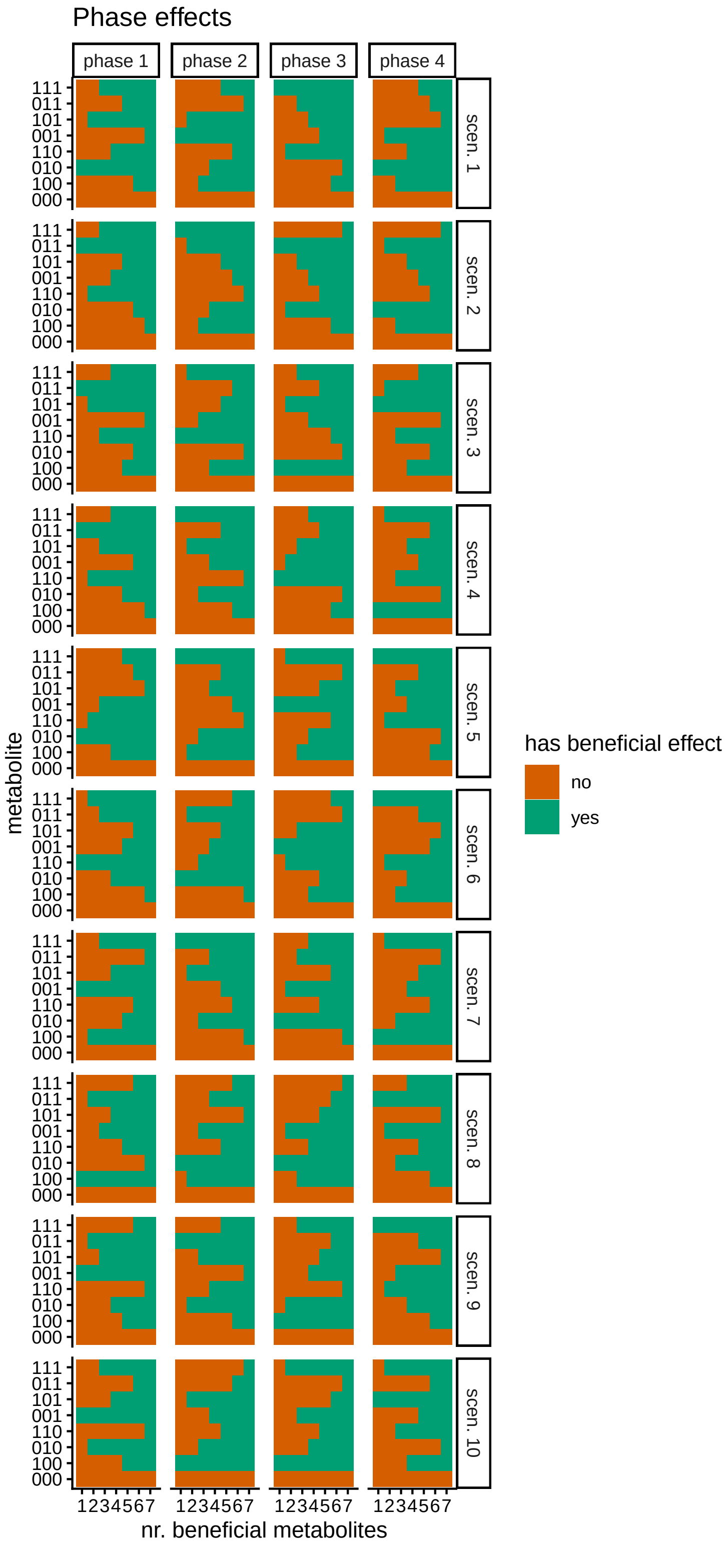


Figure S3.1: Map of the beneficial metabolites in the four different phases in the ten different experimental scenarios with three modifications. On the x-axis, the number of beneficial metabolites is indicated from 1 to 7, with the metabolite codes on the y-axis. 1 indicates that that position in the metabolite is modified, 0 indicates no modification. Red indicates that for that number of beneficial metabolites, a metabolite does not have a positive effect on fitness. Green indicates a metabolite does have a positive effect.

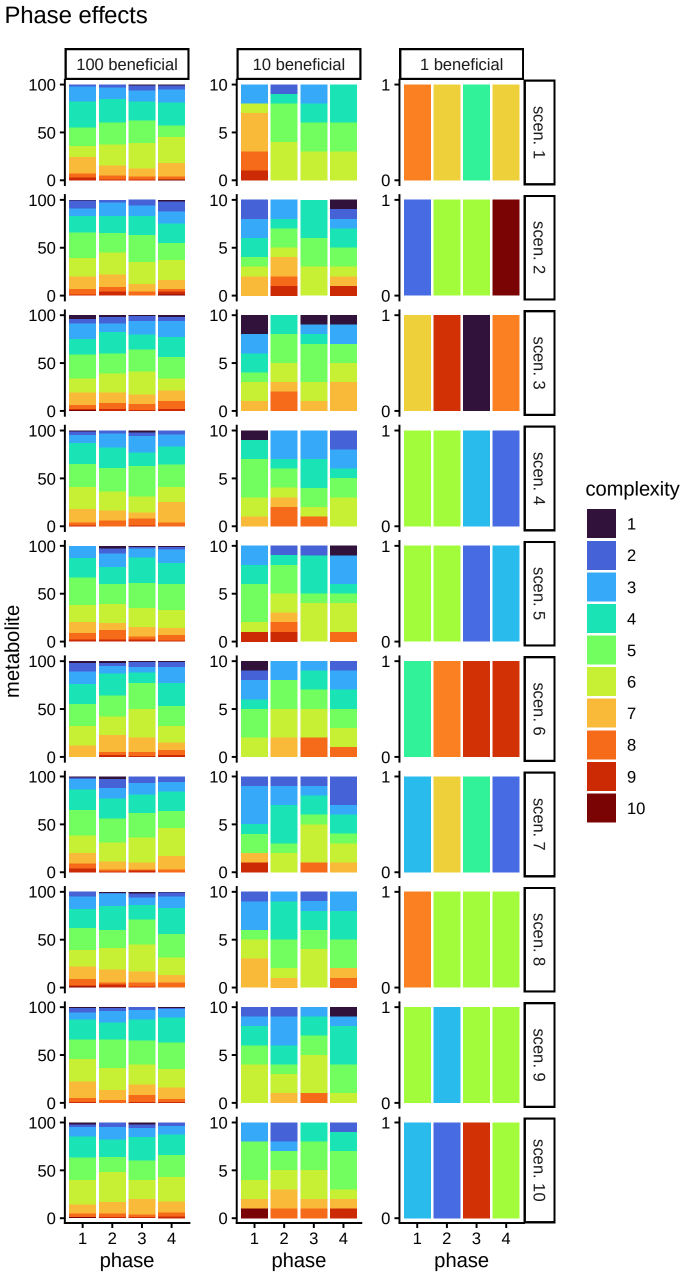


Figure S3.2: Map of the beneficial metabolites in the four different phases in the ten different experimental scenarios with ten modifications. On the x-axis, phases are indicated from 1 to 4, with the metabolites on the y-axis. Colours indicate the complexity of the metabolites, from 1 modification to 10. The three columns show the versions of the same scenario with 100, 10 and 1 beneficial metabolites. Within the same phase scenario, a version with more beneficial metabolites always includes the metabolites that are beneficial in the versions with fewer beneficial metabolites

#### S4 Allele presence and maximum

In a proportion of the simulations, at some point, all alleles are lost from the population by either drift or selection. As alleles cannot come about spontaneously, this meant no alleles would exist for the rest of the simulation. As can be seen in figures S7.3, S7.4, S7.7, and S7.8, the number of gene copies in the simulation can be much lower for simulations with fewer beneficial metabolites, and also for particular phase scenarios. In figures S4.1 and S4.2, we split out gene presence by enzyme cost, number of beneficial metabolites, and identical or different initial allele functions. We see that when enzyme cost is low, fewer simulations lose all alleles than when enzyme cost is high. Similarly, the more metabolites are beneficial, the fewer simulations lose all alleles. Additionally, in the three-modification model, when the initial alleles are different and enzyme cost is low, there are fewer simulations which lose all alleles than when initial alleles are identical.

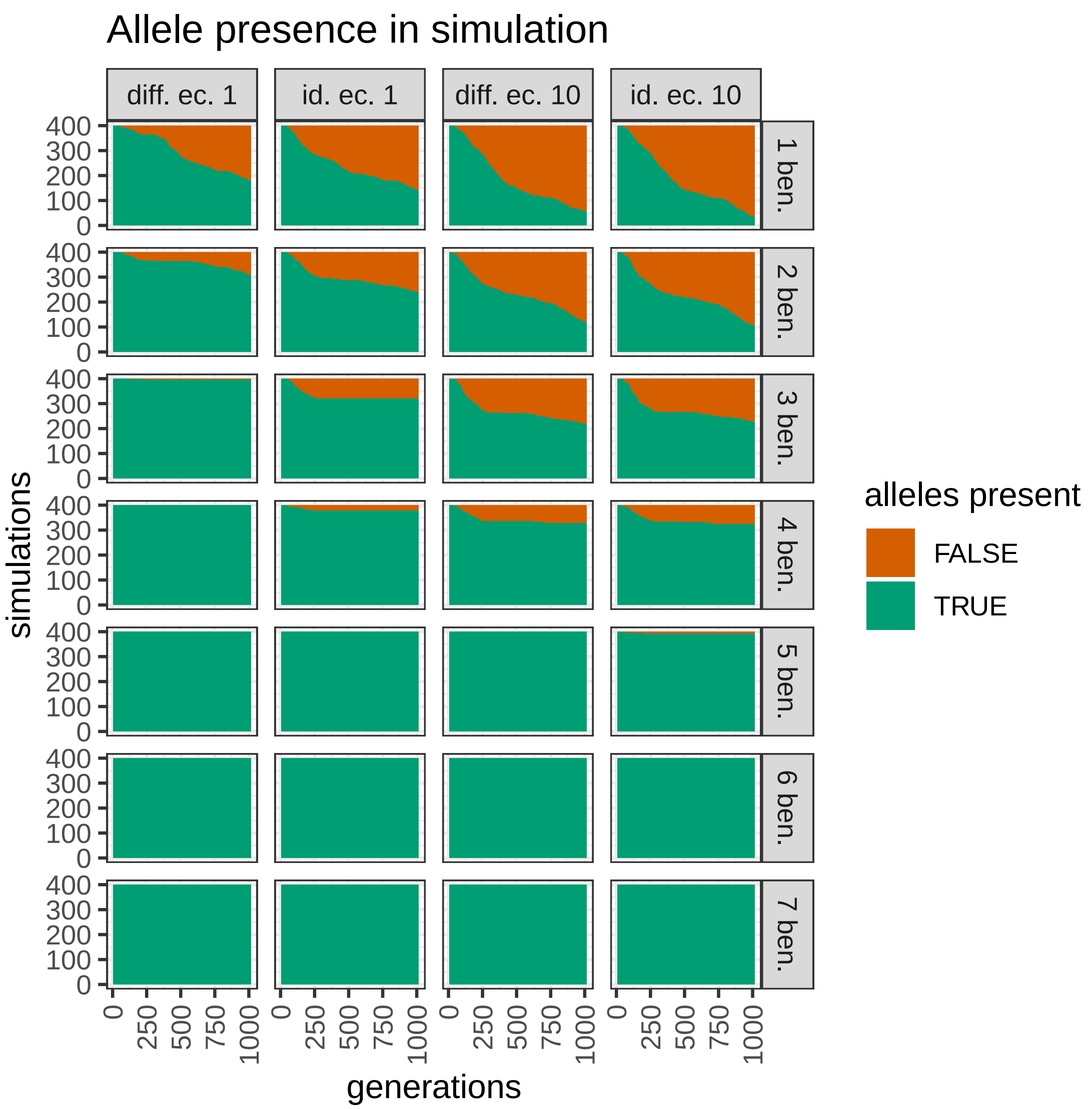


Figure S4.1: Allele presence in all simulations with three modifications through time split out by identical (id.) or different (diff.) initial allele function; low or high enzyme costs (ec.); and number of beneficial metabolites (ben.).

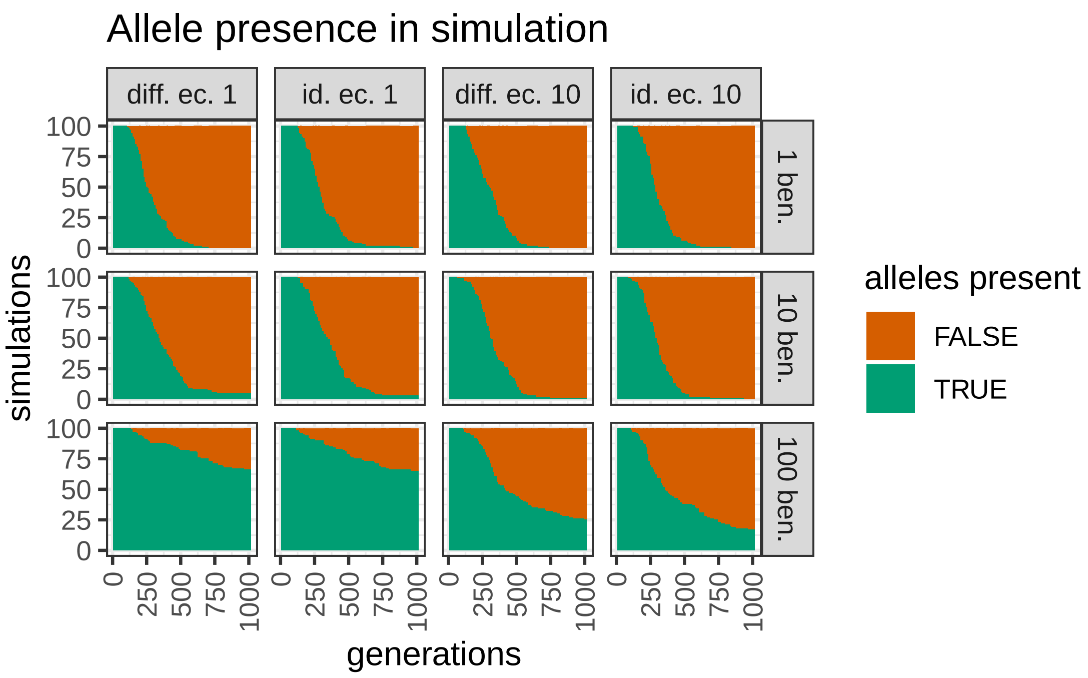


Figure S4.2: Allele presence in all simulations with ten modifications through time split out by identical (id.) or different (diff.) initial allele function; low or high enzyme costs (ec.); and number of beneficial metabolites (ben.).

We set a maximum number of 80 loci for the simulations, because the ODE that calculates the metabolite concentrations quickly gets more complex with every extra enzyme, and an overly high number of enzymes would make a simulation infeasibly slow. However, this hard limit on the number of loci carries the risk of creating artifacts in the results. Fortunately, the robustness analysis (Supplement 2) ranks the influence of the maximum number of loci low. In figure S4.3, we can see that in the simulations with three modifications, the maximum number of loci is hit only when enzyme costs are low, and even then usually not very often. This gives us confidence that our results are not strongly influenced by the maximum number of loci. For the simulations with ten modifications, none of the simulations hit the maximum number of loci at all.

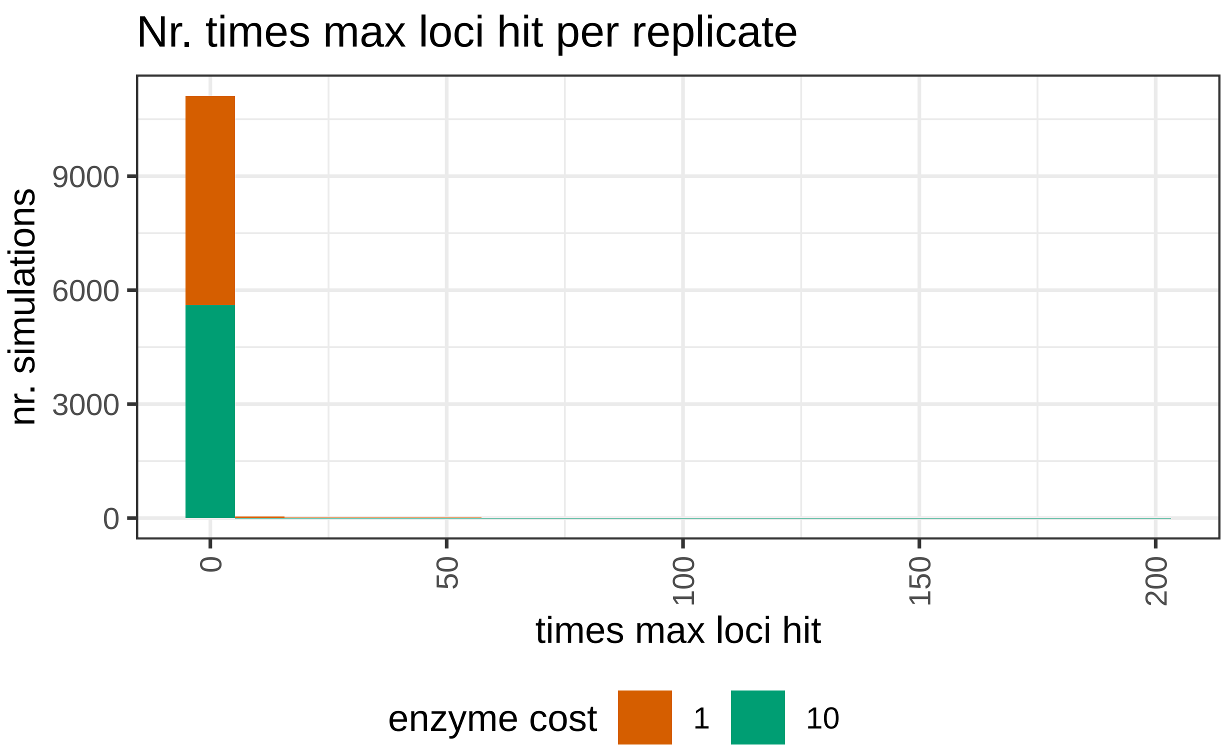


Figure S4.3: Number of simulations per number of times a simulation skipped a gene duplication during mutation because the maximum number of loci was reached in the simulations with three modifications (11200 simulations total).

### S5 Chemotype diversity through time

The specific distribution of chemotypes found in the simulation at a specific point in time is highly unpredictable due to the stochastic nature of mutations and reproduction. However, for most parameter combinations, a similar pattern of chemotypes emerges for different runs. We discuss an example here.

Figure S5.1 shows the ten replicates of a simulation where all metabolites being produced are beneficial. The ten replicates are identical in all their parameters, and differ only in their random number seed used for mutation and reproduction. At the start of the simulation, all plants have three alleles which produce enzymes that each modify a different position in the metabolite, so all possible metabolites are produced. This chemotype with all possible metabolites produced is mostly maintained throughout the simulations. In all simulations, some portion of the population has a different chemotype without any particular pattern to which chemotype at which times this is. To summarize, there appears to be selection favoring the maintenance of the production of all metabolites.

Figure S5.2 shows another set of simulations in which there is a clear selection on chemotype. It is the same scenario as the previous, except that only 4 metabolites are beneficial in each phase. The phase scenario is scenario 1 (Figure S3.1). In all simulations, in the first phase, there is quick selection for a chemotype where only metabolite 010 is made, followed by selection towards the production of all metabolites in the second phase. This is the chemotype of more than three-quarters of the plants by the end of that phase in all but two simulations. In the third phase however, the simulations diverge. Seven remain dominated by a chemotype which produces all, of which one reaches this chemotype after initially being dominated by one where the most complex chemotype is 101; one is dominated by a chemotype where the most complex metabolite is 100; and one by 010. In the last phase then, six become dominated by a chemotype 110, three 111, and one is an even mix of 110 and 111.

When looking at the beneficial metabolites within the phases (Figure S3.1), it becomes clear that the divergence happens in phase 3, where only metabolites with at least two modifications are beneficial. Through stochasticity, the system can end up with selection favoring different chemotypes, which then also has effects in the following phase, despite there being single-modification metabolites that are beneficial again.

These simulations demonstrate how, also in nature, separate populations in similar environments could develop different patterns of chemodiversity through pure chance.

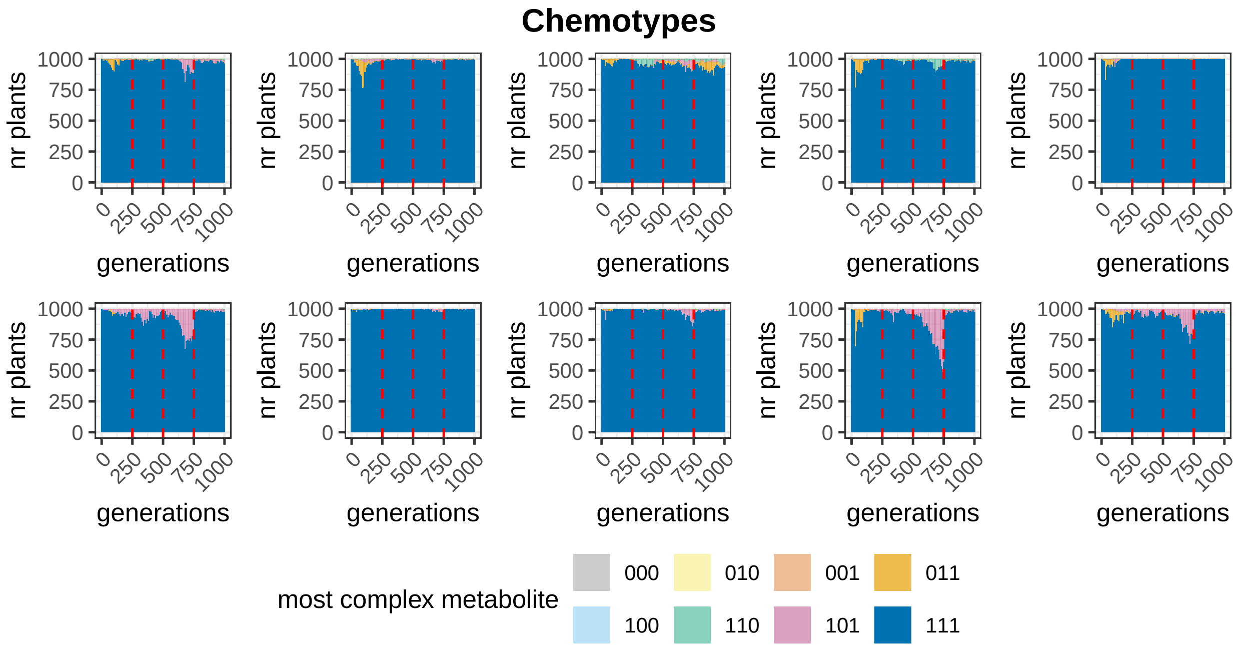


Figure S5.1: 10 replicates of a simulation with three modifications, enzyme cost 10, starting enzymes which differ in their enzyme function, and beneficial effects of all seven metabolites. Vertical red lines indicate phase switches.

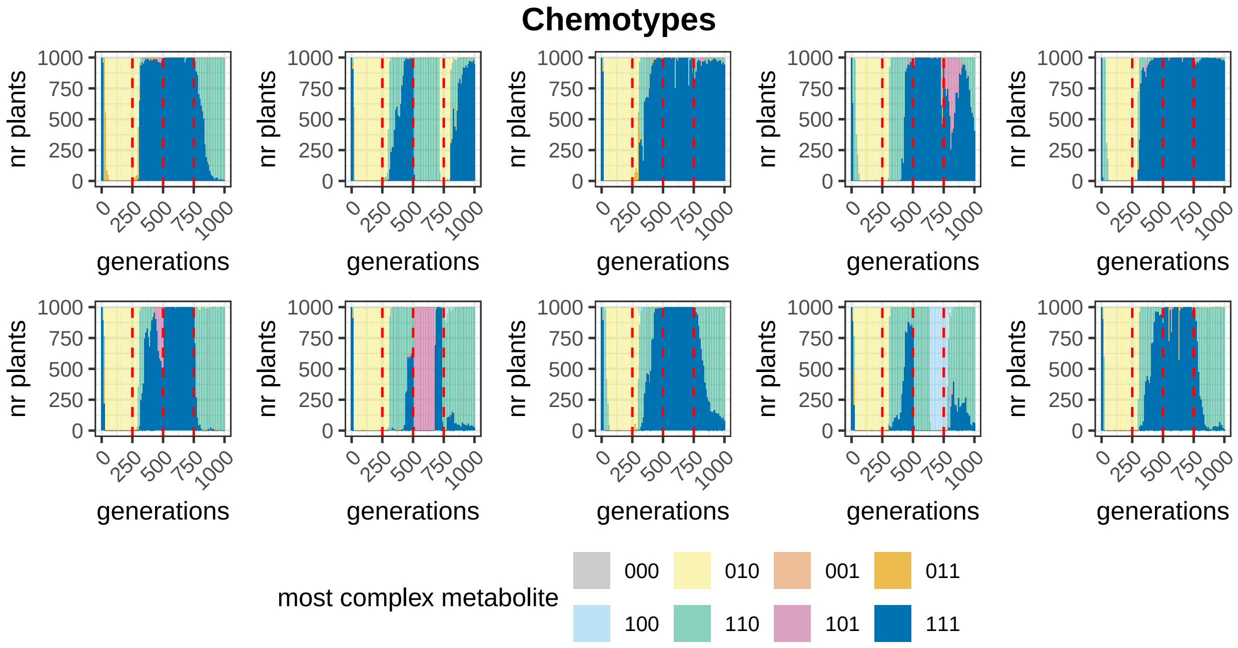


Figure S5.2: 10 replicates of a simulation with three modifications, enzyme cost 10, starting enzymes which differ in their enzyme function, and 4 metabolites in each phase conveying beneficial effects, using phase scenario 1. Vertical red lines indicate phase switches.

### S6 Promiscuity

In all scenarios with three modifications, promiscuity generally goes down (Figures S6.1, S6.2). However, promiscuity sometimes stays higher, resembling a random walk, particularly when all or most beneficial metabolites require fewer than two modifications. This pattern is seen more often in scenarios with higher enzyme costs.

In all scenarios with ten modifications, the development of promiscuity resembles a random walk (Figures S6.3, S6.4).

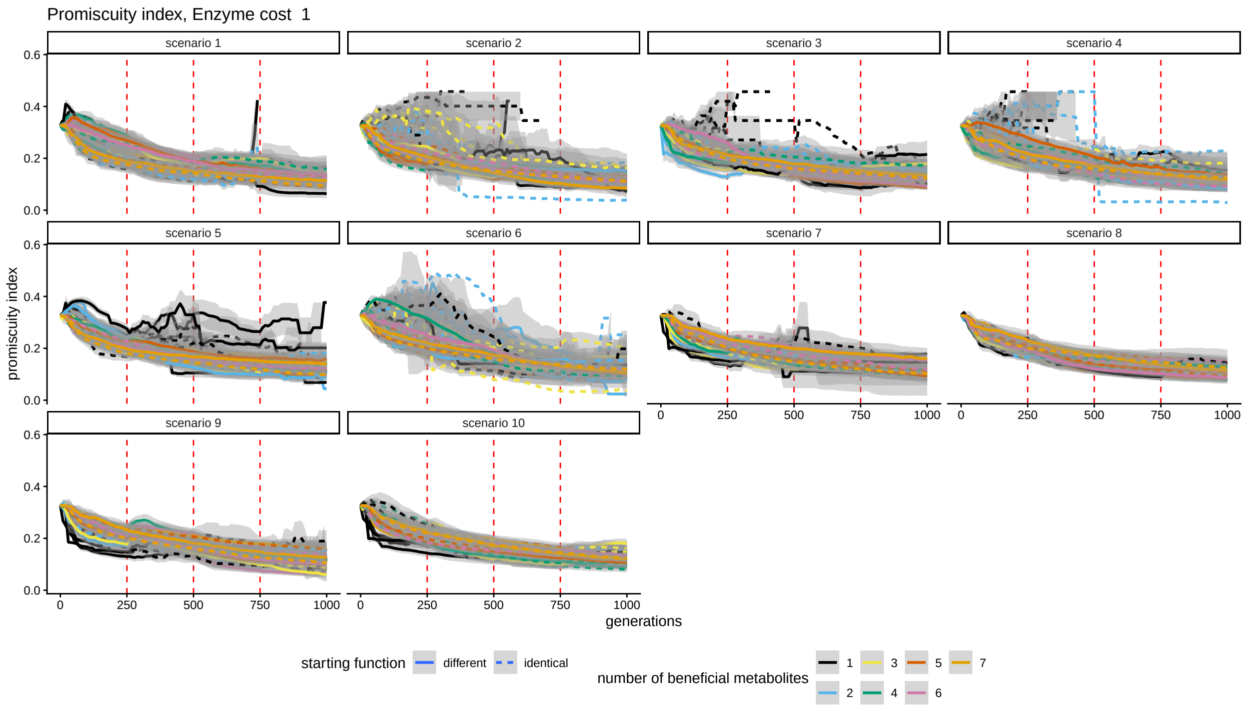


Figure S6.1: Promiscuity index split out by scenario with an enzyme cost of 1 and three modifications. Vertical red lines indicate phase switches. Every line is an average over 10 simulations with the same fitness effects of beneficial metabolites at each timestep. The bands represent standard errors.

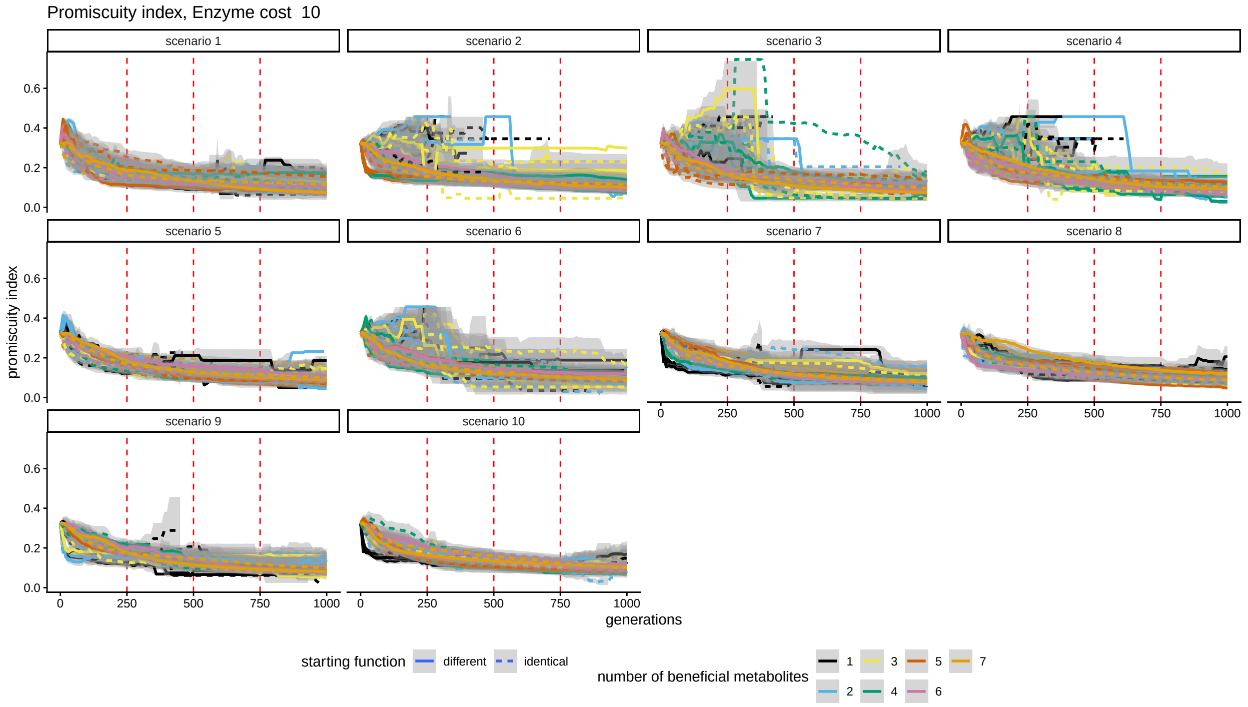


Figure S6.2: Promiscuity index split out by scenario with an enzyme cost of 10 and three modifications. Vertical red lines indicate phase switches. Every line is an average over 10 simulations with the same fitness effects of beneficial metabolites at each timestep. The bands represent standard errors.

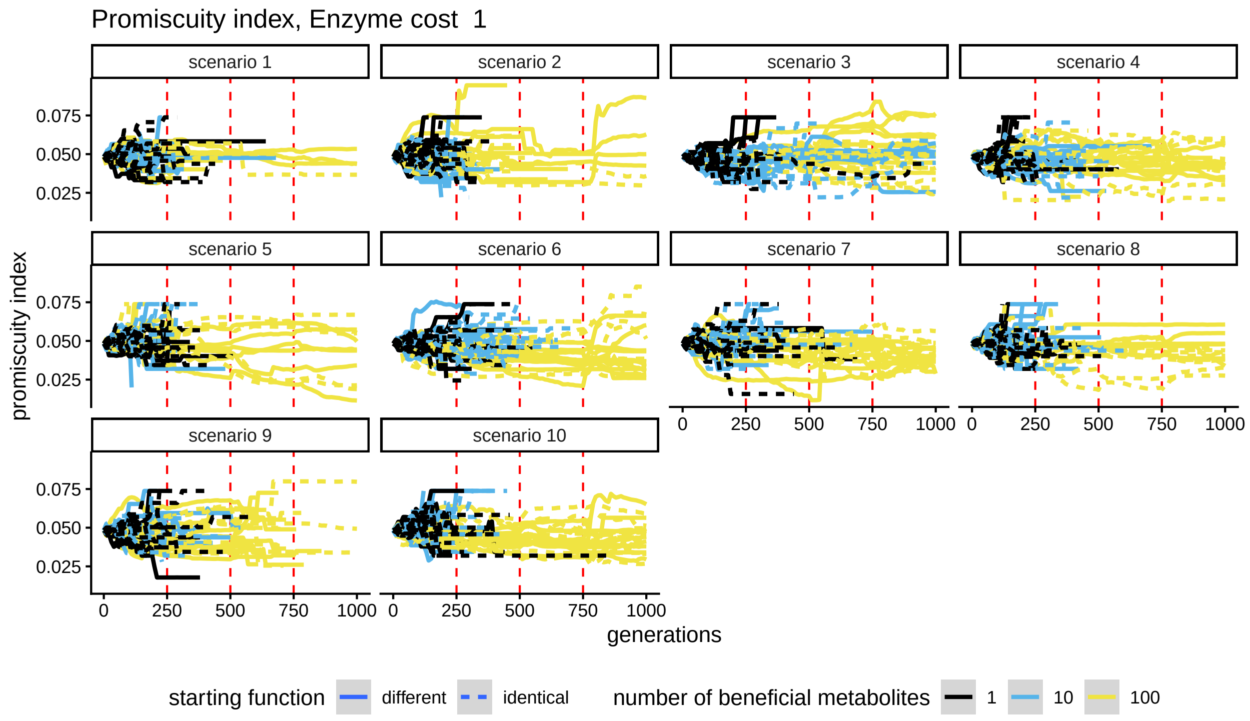


Figure S6.3: Promiscuity index split out by scenario with an enzyme cost of 1 and ten modifications. Vertical red lines indicate phase switches. Every line is one simulation. The bands represent standard errors.

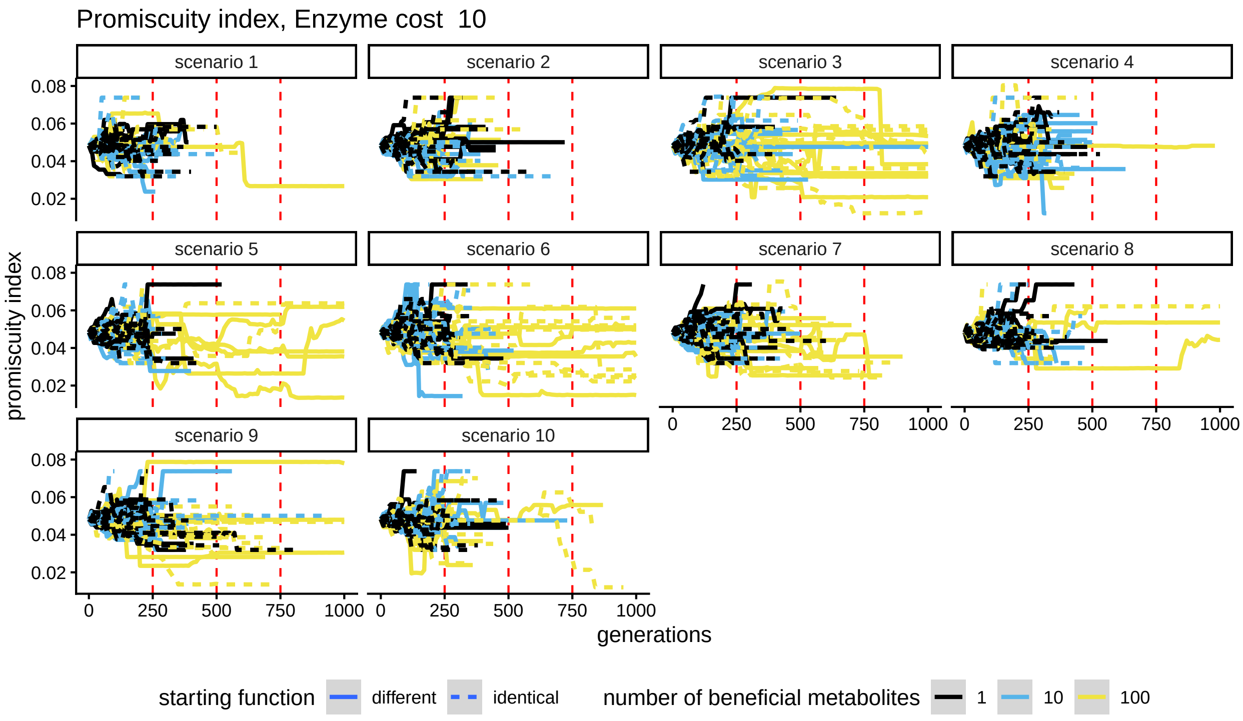


Figure S6.4: Promiscuity index split out by scenario with an enzyme cost of 10 and ten modifications. Vertical red lines indicate phase switches. Every line is one simulation. The bands represent standard errors.

### S7 Selection/drift

The SoA and NoG are roughly correlated with metabolite richness (Supplement 9, Figures S9.1, S9.2). The more metabolites are being made, the more alleles and gene copies there are. At the same time, there are fewer alleles and gene copies when enzyme costs are higher (Figures S7.1, S7.2).

When we split out the three-modification simulations by phase scenario, it becomes clear that the drop and subsequent spike in SoA after phase changes is not universal for all scenarios, but rather is found when there is a mismatch between the beneficial metabolites. In scenario 6 with four beneficial metabolites for example, there are always two beneficial metabolites with one modification and one beneficial metabolite with both those modifications in the last three phases, but in every phase one of these modifications is different from the previous ones (Figure S3.1). In both the scenario with high and low enzyme costs, this results in a sawtooth pattern, though for low enzyme costs, the SoA drops relatively lower and does not recover as well as for high enzyme costs (Figures S7.1, S7.2).

When there are more or fewer metabolites with only one modification in a phase than in the previous one, the SoA can rise higher or stay lower than in the previous phase, respectively. For example, in phase scenario 9 with three beneficial metabolites, the third phase has two beneficial metabolites with one modification, the fourth one only one (Figure S3.1). Both with high and low enzyme cost, the SoA drops and stays low in the fourth phase (Figures S7.1, S7.2). Interestingly, in this scenario, the first two phases also have only one beneficial metabolite with one modification, but at low enzyme cost, the SoA is just as high in the first two phases as in the third phase. When enzyme costs are high, these phases have a low SoA (and an average richness of 1, see Figure S9.2), just like the fourth phase. When enzyme costs are low, they have a similar SoA to the third phase (and a similar richness at the end of the phase, see Figure S9.1). The difference is that in the first two phases, the beneficial metabolite with one modification is a precursor to at least one of the two beneficial metabolites with two modifications, while in the fourth phase, the beneficial metabolite with one modification is not a precursor to the beneficial metabolite with two modifications, and the metabolite it is a precursor to has three modifications. Despite that the beneficial metabolites with one modification from the third phase are the precursors of the beneficial metabolite with two modification in the fourth phase, the genes which produce enzymes that make these modifications are selected away. This is evidence of a strong selection against making non-beneficial precursors when there is a less-complex beneficial metabolite available. When enzyme costs are high, this selection is stronger, so that a more complex metabolite is not made if its precursors are not beneficial. When more metabolites with more modifications are made, the

SoA and NoG are higher than when fewer metabolites are made (Figures S7.1, S7.2, S7.3, S7.4, S9.1, S9.2).

However, sometimes, a sharp increase in the SoA can be seen instead when enzyme costs are low and only more complex metabolites are beneficial in the new phase (Figure S7.1). In phase scenario 1, with three or four beneficial metabolites, the effect of the phase change from phase 2 to 3 is opposite for low or high enzyme costs (Figures S7.1, S7.2). The second phase has two (for three beneficial metabolites) or three (for four beneficial metabolites) beneficial metabolites with only one modification and one with two modifications of which both precursors are beneficial. The third phase in contrast has only beneficial metabolites with two or three modifications, with none of the precursors being beneficial, for both scenarios (Figure S3.1). Due to this, the plants produce metabolites with up to two modifications by the end of the second phase (Figures S9.1, S9.2). Then, after the phase switch, the metabolites with one modification are not beneficial anymore, and the metabolite with two modificaitons only in the scenario with four beneficial metabolites. When enzyme costs are high, and there are not so many alleles or gene copies to select and mutate to fit the new situation, the SoA and NoG go down rapidly, as there is strong selection to reduce costs by reducing the number of gene copies. They start to rise again towards the end of the phase for the scenario with four beneficial metabolites only (Figure S7.2). However, when enzyme costs are low, there is a veritable radiation of alleles, including a slight increase in promiscuity (Figures S7.1, S6.1). This raises both the SoA and the total number of gene copies even over those of the scenario where all metabolites are beneficial. The version with 5 beneficial metabolites of this phase scenario, however, includes a single beneficial metabolite with a single modification in the third phase, and does not display this radiation. A similar pattern can be seen for the three and/or two -beneficial metabolite versions of phase scenarios 3, 5, and 10 between phases 3 and 4; and in the first phases of the three- and four-beneficial metabolite versions of phase scenario 3 and 4 (Figures S7.1, S7.3).

Simulations with low numbers of beneficial metabolites are prone to lose all alleles, particularly when enzyme costs are high (Figures S7.2, S7.4). This is also true for the ten-modification simulations (Figures S7.6, S7.8). With ten modifications and 100 beneficial metabolites, there usually is a dip in the SoA and NoG after phase changes, followed by an increase when there is at least one beneficial metabolite with only one modification. When there are no beneficial metabolites with only 1 modification, and enzyme cost is high, the SoA and NoG stay low. However, when enzyme cost is low, the NoG and SoA recover in many cases also when there are no beneficial metabolites with only one modification. For scenarios with 10 and 1 beneficial metabolites we find a high SoA or NoG only when at least one beneficial metabolite has only one modification. For 10 beneficial metabolites, this is in phase 1, 3, and 4 of scenario 3, and phase 1 of scenario

4 and 6 at low enzyme cost, and phase 1 of scenario 3, 4, and 6 at high enzyme cost (Figures S3.2, S7.5, S7.6).

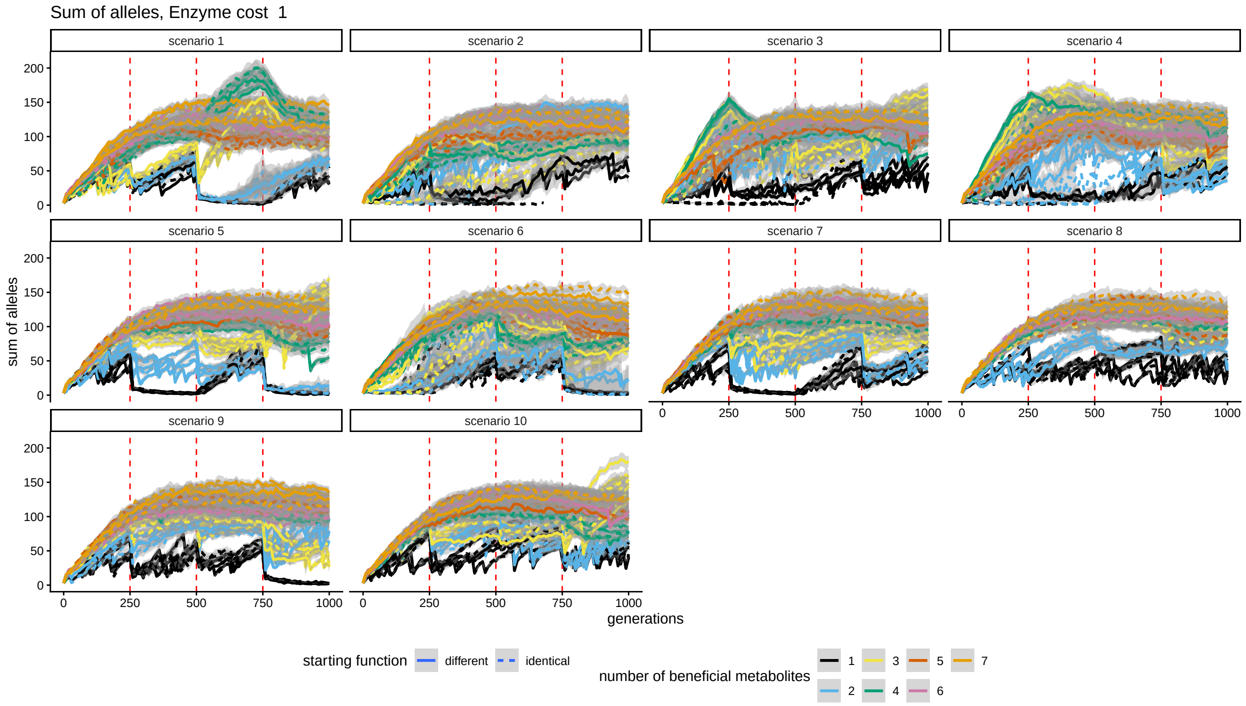


Figure S7.1: Sum of alleles across loci split out by scenario: enzyme cost of 1 and three modifications. Vertical red lines indicate phase switches. Every line is an average over 10 simulations with the same fitness effects of beneficial metabolites at each timestep. The bands represent standard errors.

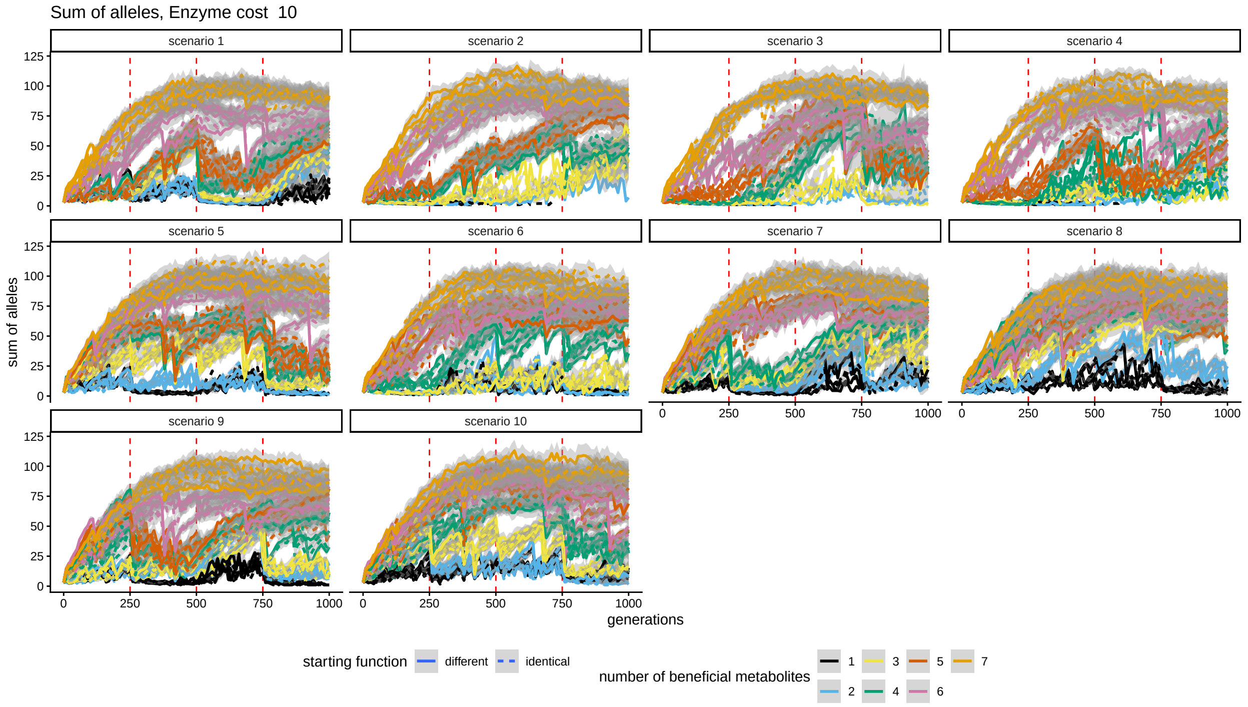


Figure S7.2: Sum of alleles across loci split out by scenario: enzyme cost of 10 and three modifications. Vertical red lines indicate phase switches. Every line is an average over 10 simulations with the same fitness effects of beneficial metabolites at each timestep. The bands represent standard errors.

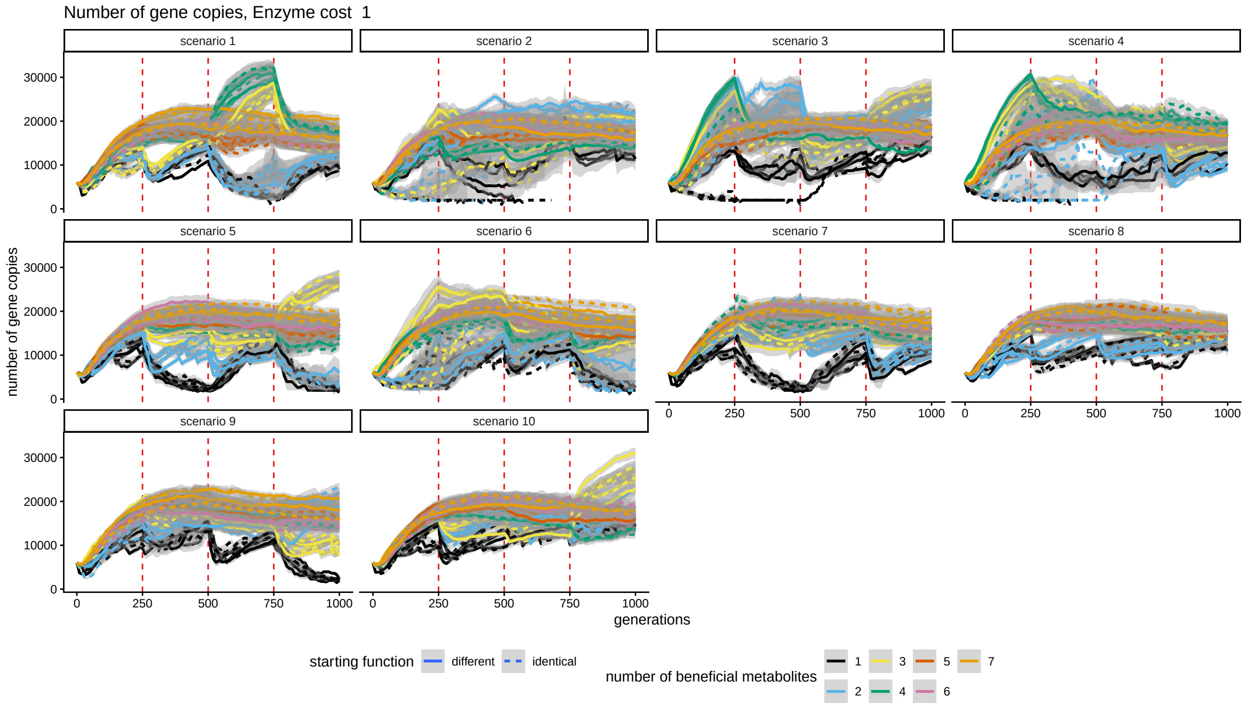


Figure S7.3: Total number of gene copies split out by scenario: enzyme cost of 1 and three modifications. Vertical red lines indicate phase switches. Every line is an average over 10 simulations with the same fitness effects of beneficial metabolites at each timestep. The bands represent standard errors.

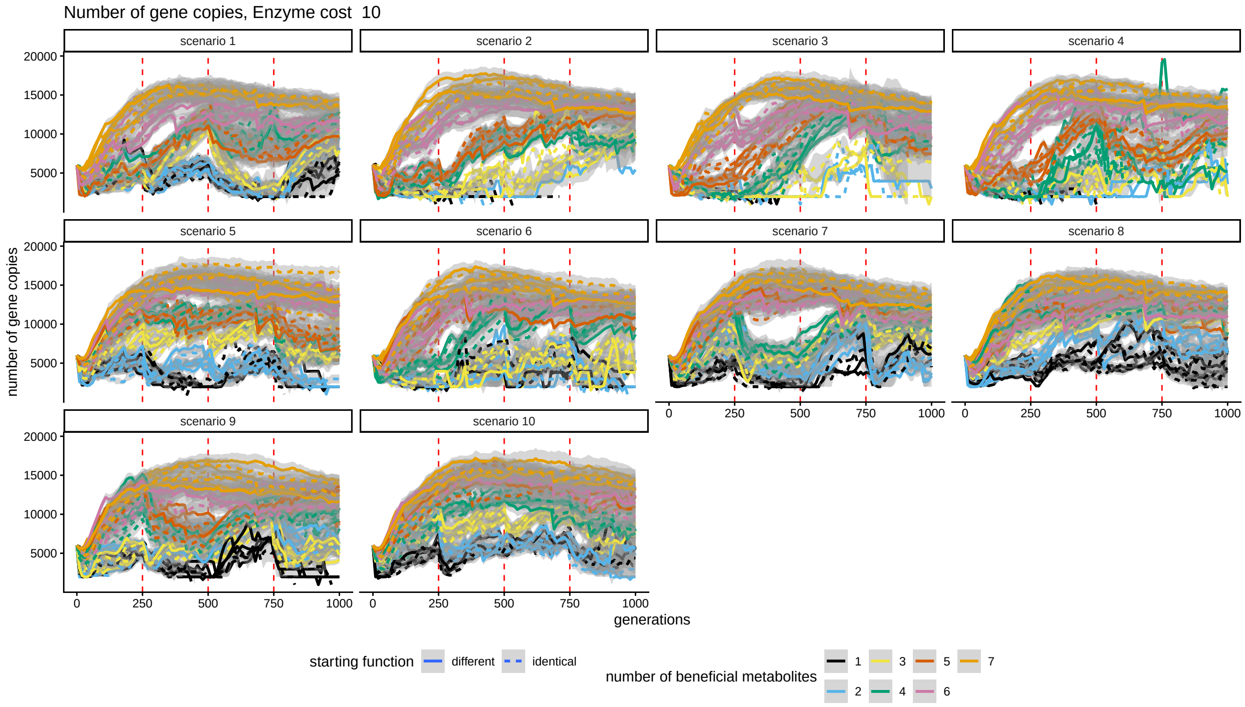


Figure S7.4: Total number of gene copies split out by scenario: enzyme cost of 10 and three modifications. Vertical red lines indicate phase switches. Every line is an average over 10 simulations with the same fitness effects of beneficial metabolites at each timestep. The bands represent standard errors.

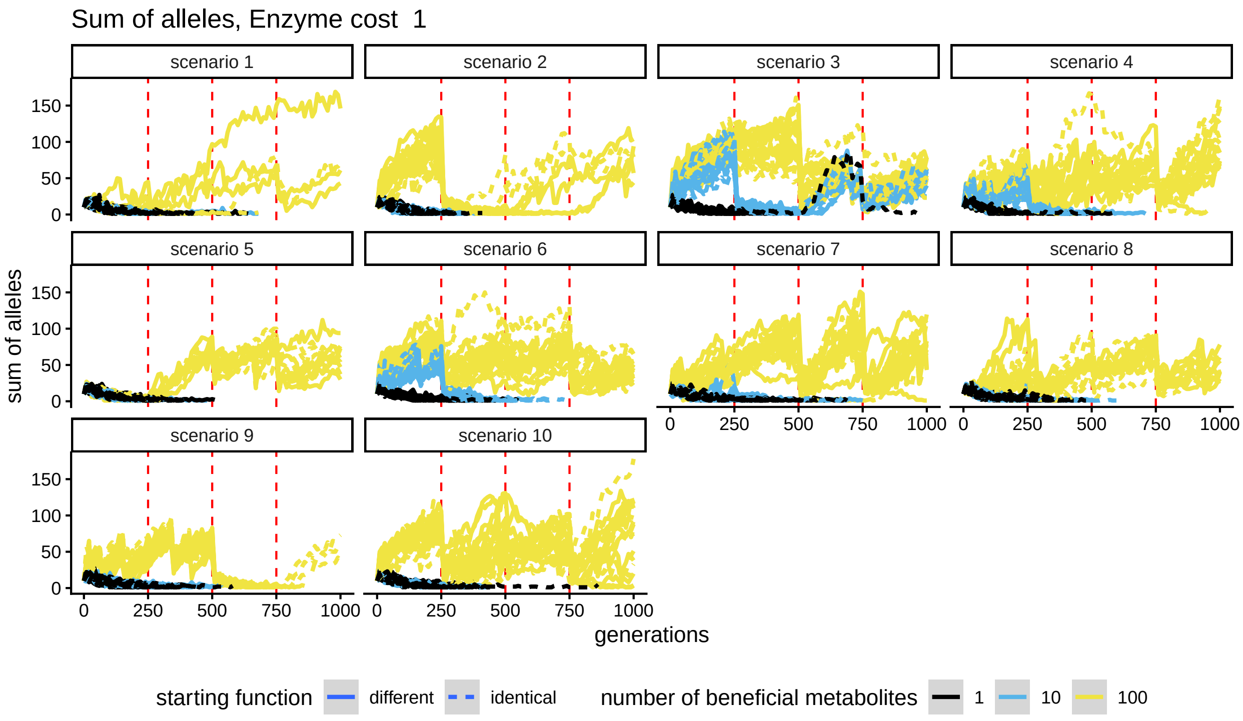


Figure S7.5: Sum of alleles across loci split out by scenario: enzyme cost of 1 and ten modifications. Vertical red lines indicate phase switches. Every line is one simulation. The bands represent standard errors.

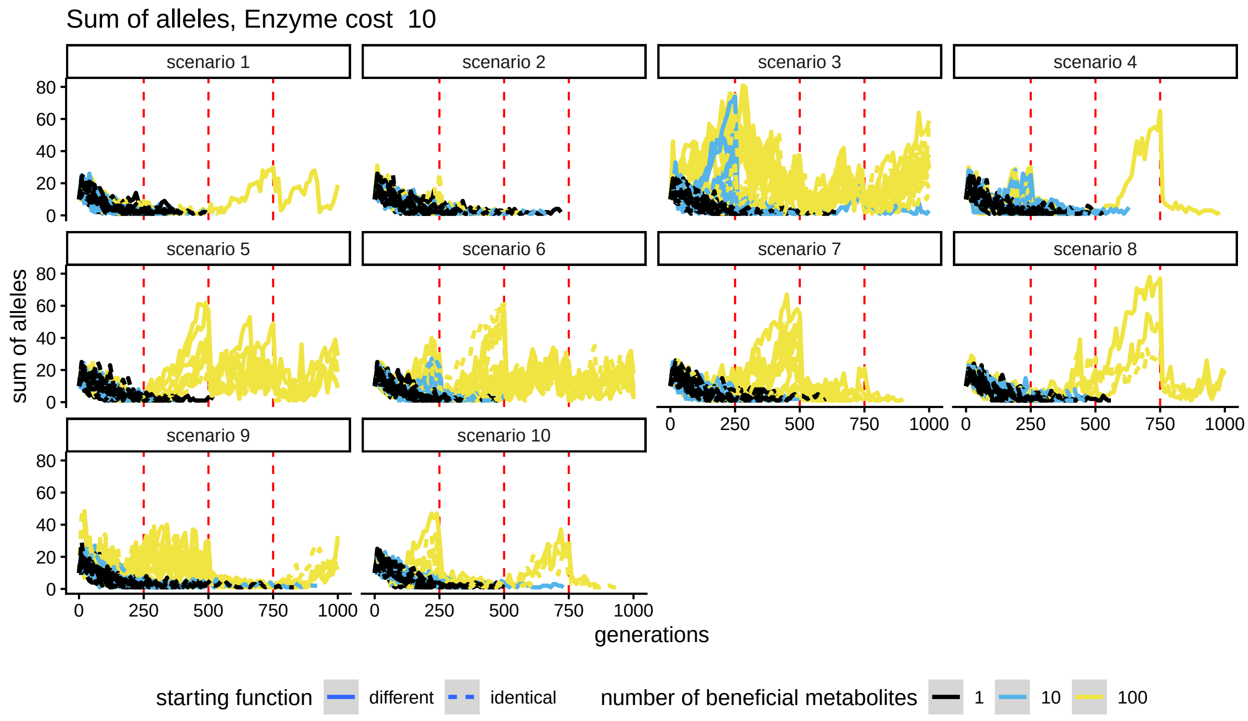


Figure S7.6: Sum of alleles across loci split out by scenario: enzyme cost of 10 and ten modifications. Vertical red lines indicate phase switches. Every line is one simulation. The bands represent standard errors.

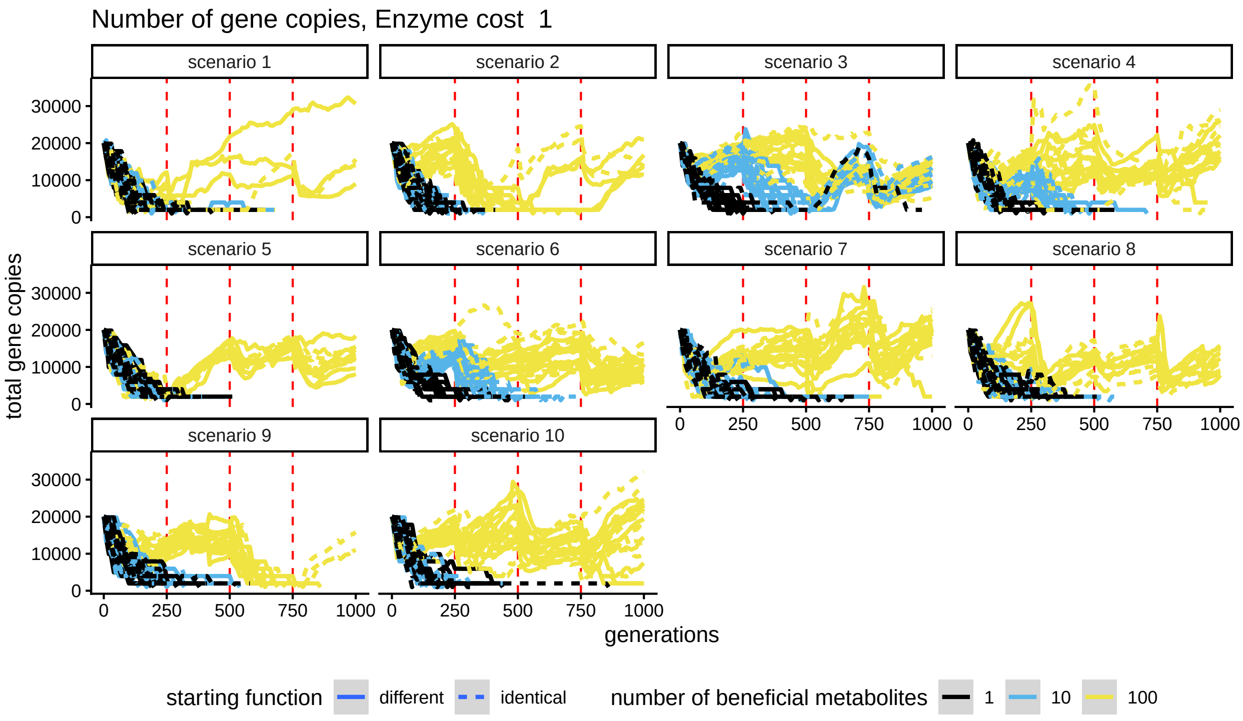


Figure S7.7: Total number of gene copies split out by scenario: enzyme cost of 1 and ten modifications. Vertical red lines indicate phase switches. Every line is one simulation. The bands represent standard errors.

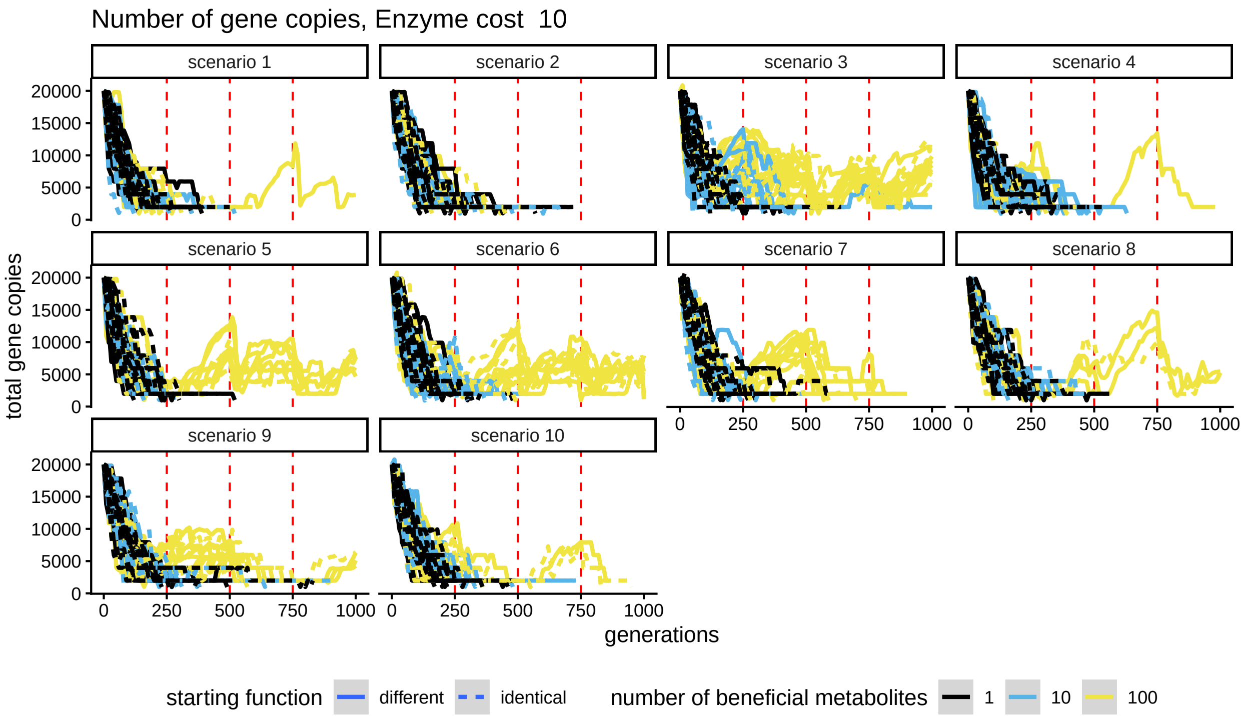


Figure S7.8: Total number of gene copies split out by scenario: enzyme cost of 10 and ten modifications. Vertical red lines indicate phase switches. Every line is one simulation. The bands represent standard errors.

### S8 Match with beneficial metabolites

Generally, the ratio of beneficial metabolites to total metabolites is either high, or zero, in all simulations (Figures S8.1, S8.2, S8.3, S8.4). The ratio is lower in phases where there are few beneficial metabolites and these metabolites are more complex, usually near or at zero, particularly when enzyme costs are high (Compare for example phase 3 in scenario 1 in the simulations with three modifications, Figures S8.1, S8.2, S3.1). Within phases, metabolites reach the maximum ratio faster when enzyme costs are high (Figures S8.2, S8.4).

For the three-modification scenarios, at low enzyme costs, some of the scenarios where the ratio is near or at zero for high enzyme costs, show an intermediate ratio instead. This happens when there are few beneficial metabolites which all have at least two modifications (such as in scenario 1, phase 3, figures S8.1, S8.2). The plants produce non-beneficial metabolites with fewer modifications as precursors for beneficial metabolites. The ratio of beneficial metabolite made increases over time, indicating an evolution to a more efficient production of beneficial metabolites. For the ten-modification scenarios, the phases with ratios close to 1 line up perfectly with the phases that have at least one beneficial metabolite with only one modification. This high ratio is again reached faster at higher enzyme cost. Many of those scenarios with 100 beneficial metabolites and lower enzyme cost show an intermediate and growing ratio in phases where such a metabolite is not present, while those same phases show a ratio at or near zero for their equivalent with high enzyme cost (Figures S8.3, S8.4).

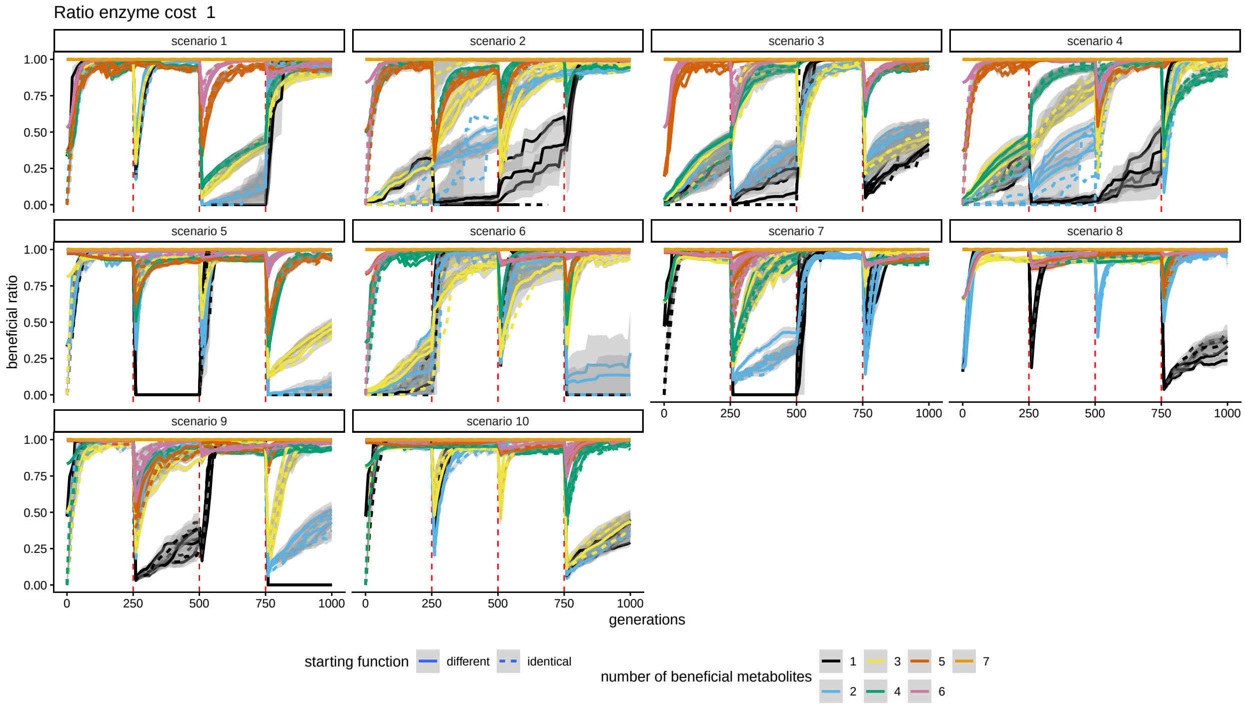


Figure S8.1: The ratio of beneficial metabolites divided by all metabolites, split out by scenario: enzyme cost of 1 and three modifications. Vertical red lines indicate phase switches. Every line is an average over 10 simulations with the same fitness effects of beneficial metabolites at each timestep. The bands represent standard errors.

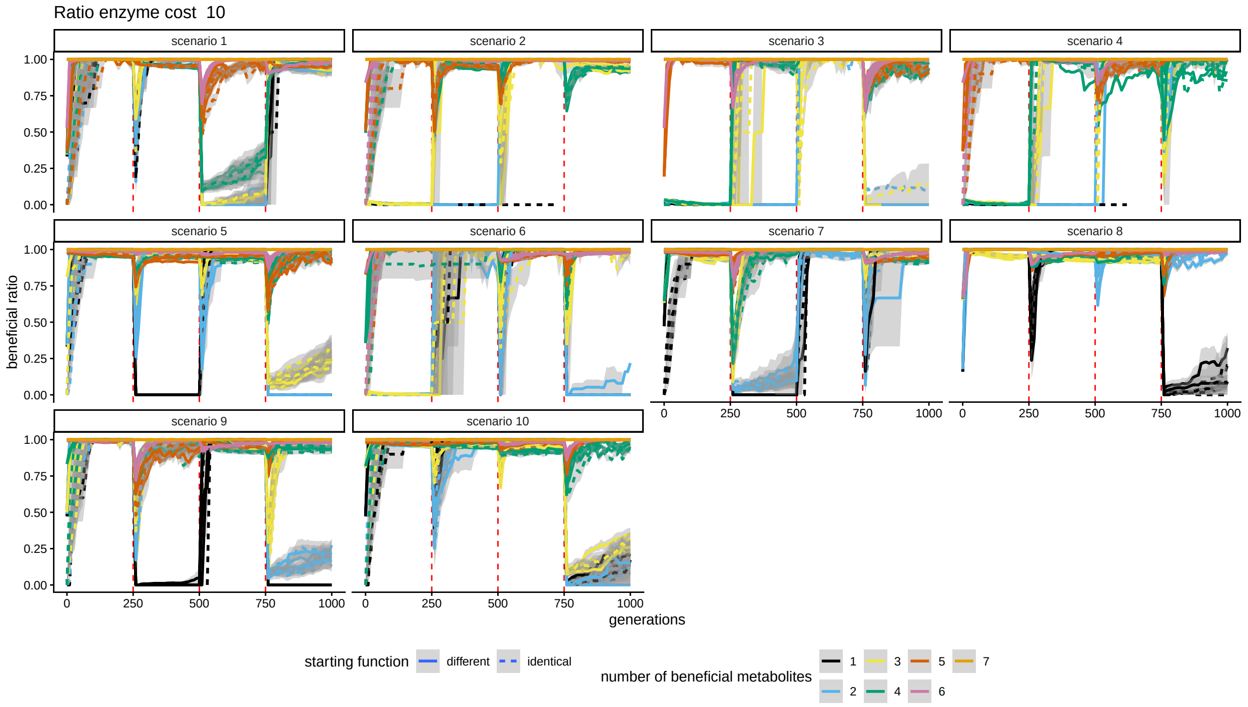


Figure S8.2: The ratio of beneficial metabolites divided by all metabolites, split out by scenario: enzyme cost of 10 and three modifications. Vertical red lines indicate phase switches. Every line is an average over 10 simulations with the same fitness effects of beneficial metabolites at each timestep. The bands represent standard errors.

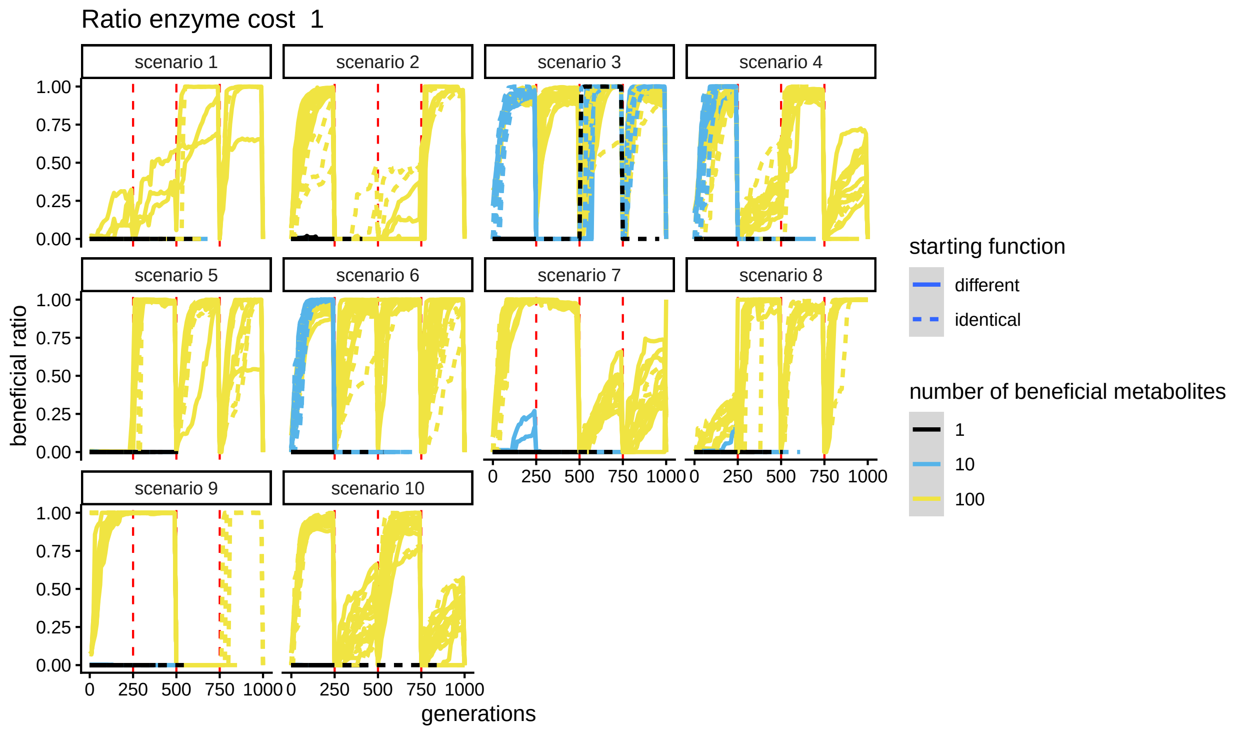


Figure S8.3: The ratio of beneficial metabolites divided by all metabolites, split out by scenario: enzyme cost of 1 and ten modifications. Vertical red lines indicate phase switches. Every line is one simulation. The bands represent standard errors.

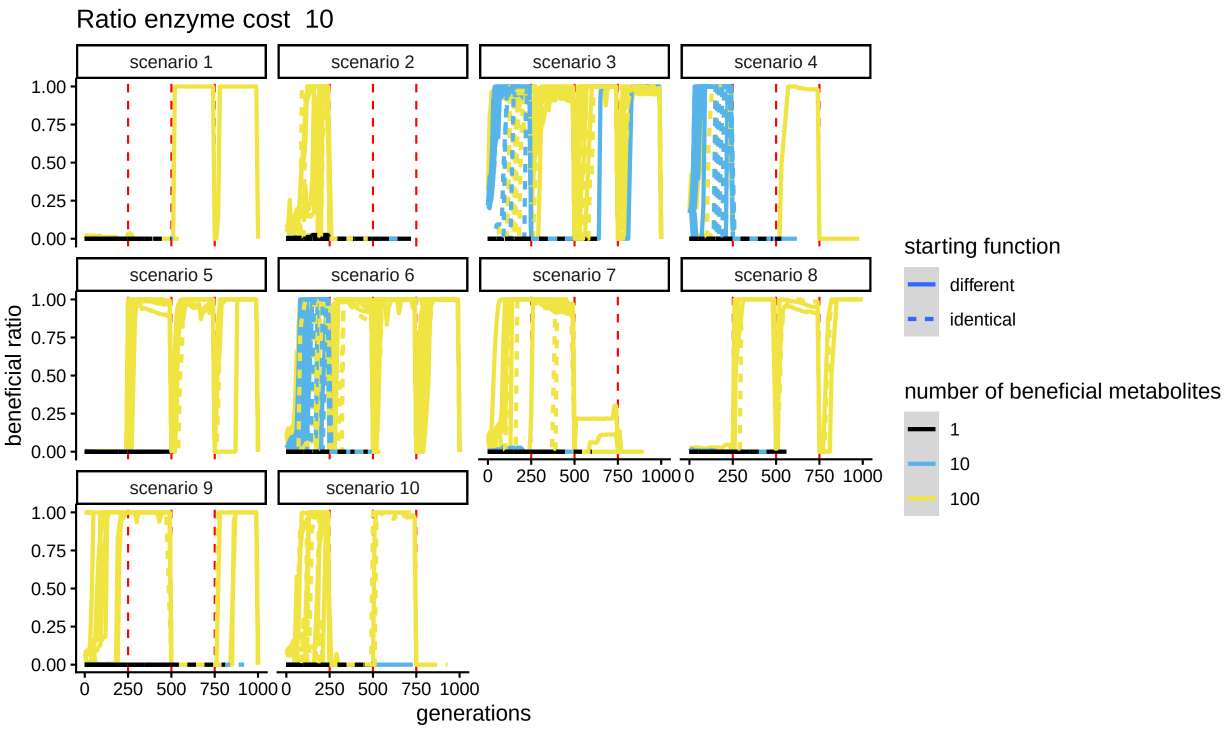


Figure S8.4: The ratio of beneficial metabolites divided by all metabolites, split out by scenario: enzyme cost of 10 and ten modifications. Vertical red lines indicate phase switches. Every line is one simulation. The bands represent standard errors.

### S9 Richness

In the three-modification scenarios, metabolite richness depends highly on which metabolites are beneficial. This means that higher numbers of beneficial metabolites sometimes result in higher richness, but not always. In simulations with fewer beneficial metabolites, a major change in which modifications are required to make beneficial metabolites from one phase to the next causes spikes at the start of the new phase. This is because selection favors plants that produce the newly beneficial metabolites and the enzymes that produce them, while the enzymes that produce the newly non-beneficial metabolites have not yet been selected away (Figures S9.1, S9.2).

Complex metabolites being beneficial does not necessarily result in higher richness. In low enzyme cost scenarios, there is high richness if complex metabolites can be made with existing enzymes, and more simple metabolites are not beneficial, see for example phase 3 in scenario 1 or phase 1 for scenario 4 with different starting functions (Figure S9.1). When simpler metabolites become beneficial however, the richness drops to the richness that is expected when all of the least complex beneficial metabolites are made (compare for example phase 1 and 2 for scenario 3 with 2 beneficial metabolites with phase 3 of the same, or phase 1 and 2 for the same scenario with 3 beneficial metabolites (Figures S9.1, S3.1)). In high enzyme cost scenarios, richness is driven by the simple metabolites. The richness consistently corresponds to the richness expected when all those metabolites which are both beneficial and only have beneficial precursors are made. That is to say, a more complex metabolite may be made, despite not being beneficial, if its precursors are beneficial, but a beneficial metabolite is not made if its precursors are not also beneficial (Figure S9.2, S3.1).

As ratios of beneficial to non-beneficial metabolites are high in almost all scenarios (Figure S8.1, S8.2, S8.3, S8.4), it is clear that a high richness does not correlate with a large amount of non-beneficial metabolites being made.

In the ten-modification scenarios, richness is generally low compared to how high it could theoretically be. In the 100 beneficial metabolite scenarios, it spikes at the start of phases as the specifics of which metabolites are made changes, higher in scenarios with low enzyme cost. There are differences between phase scenarios, but it is unclear what the cause of those patterns is (Figures S9.3, S9.4).

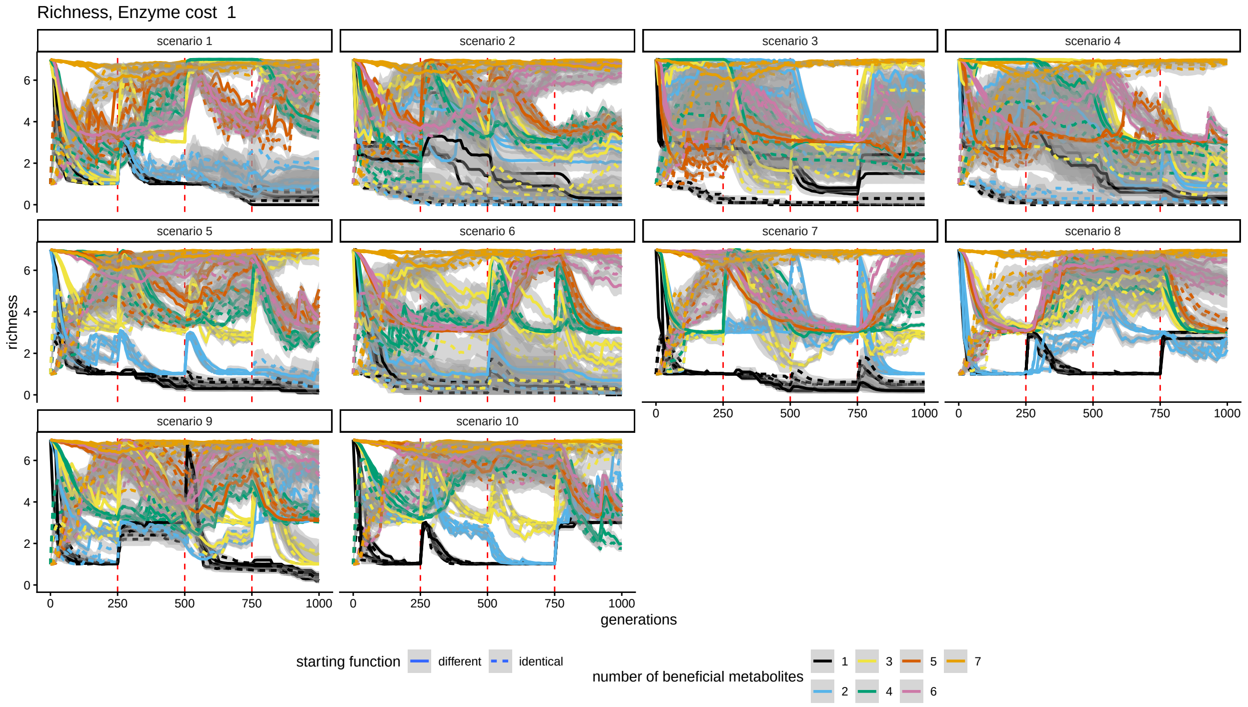


Figure S9.1: Metabolite richness split out by scenario: enzyme cost of 1 and three modifications. Vertical red lines indicate phase switches. The bands represent standard errors.

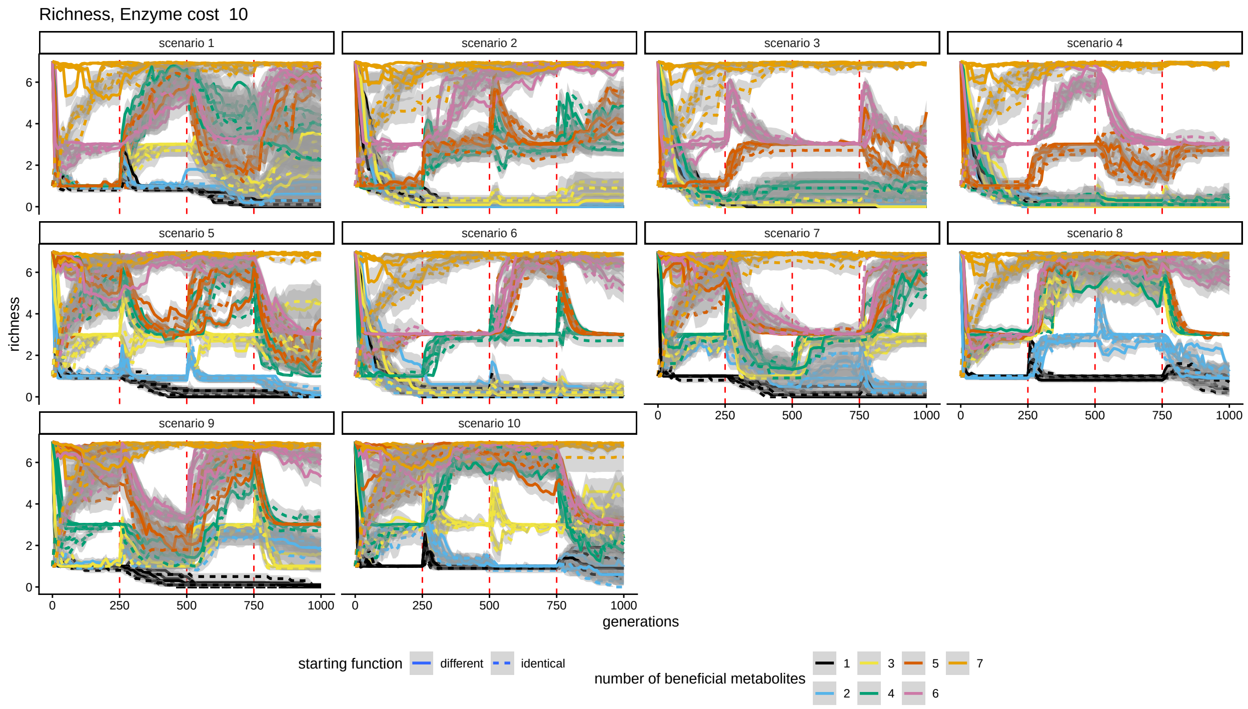


Figure S9.2: Metabolite richness split out by scenario: enzyme cost of 10 and three modifications. Vertical red lines indicate phase switches. The bands represent standard errors.

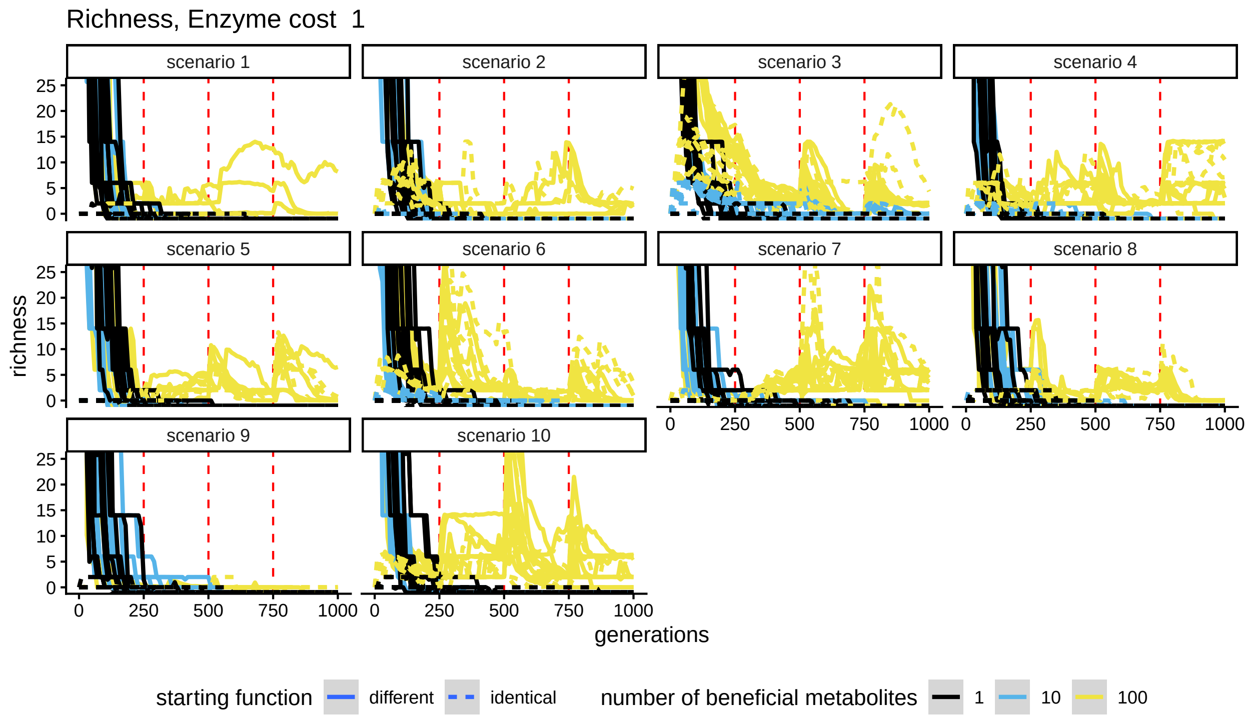


Figure S9.3: Metabolite richness split out by scenario: enzyme cost of 1 and ten modifications. Vertical red lines indicate phase switches. The bands represent standard errors.

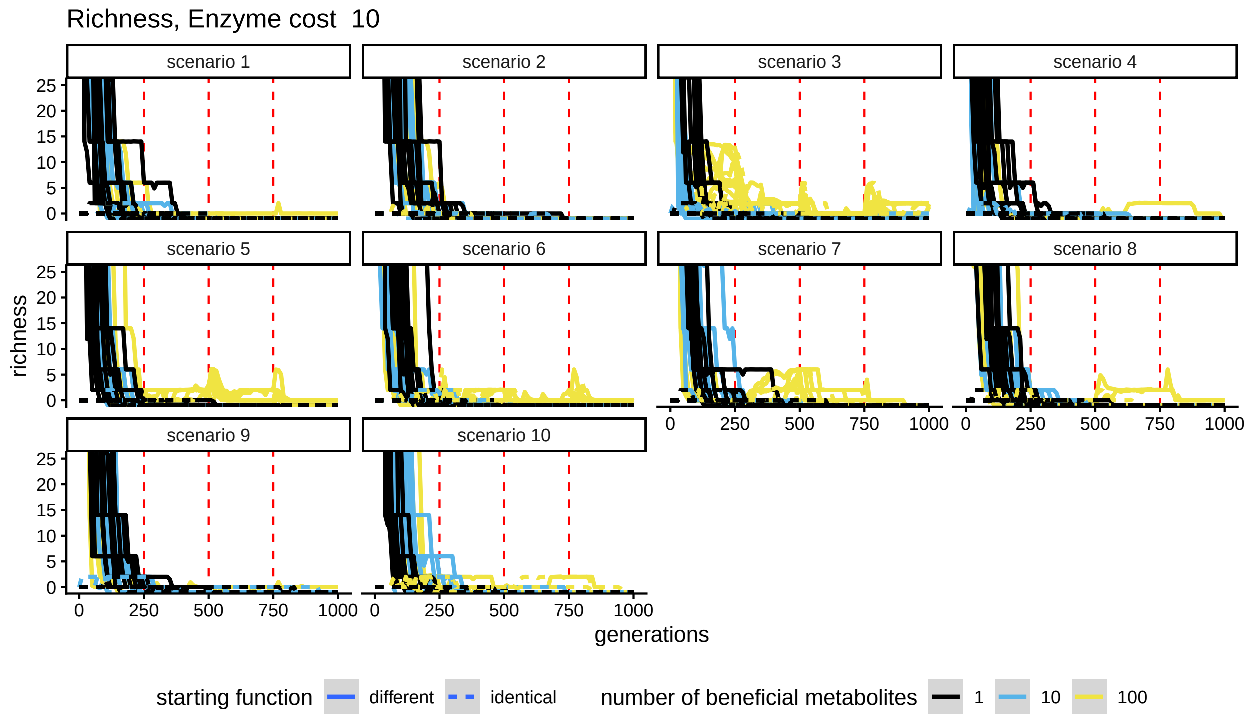


Figure S9.4: Metabolite richness split out by scenario: enzyme cost of 10 and ten modifications. Vertical red lines indicate phase switches. The bands represent standard errors.

### S10 Summary by number of beneficial metabolites

When we plot the metrics against number of beneficial metabolites instead of against time, we see the same patterns in a different way.

In the scenarios with 3 modifications, we see that promiscuity is lower every phase. However, promiscuity also is lower the more beneficial metabolites there are in the first two phases, if enzyme costs are high or the initial metabolites have the same function (S10.1 A). In the scenarios with 10 modifications, there is not much of a pattern in the promiscuity index. This may be because the specificity constant for a reaction is not selected on if that reaction cannot take place, and that is the case for most reactions during the vast majority of timesteps in all simulations, rendering the promiscuity index less informative (S10.2 A).

The SoA and NoG increase with a higher number of beneficial metabolites, though there is a kind of maximum when enzyme costs are low, in the simulations with three possible modifications (S10.1 B, C, S10.2 B, C). The ratio of beneficial metabolites generally goes up the more metabolites are beneficial, but there is a lot of variation, particularly in the simulations with 10 possible modifications (S10.1 D, S10.2 D). Metabolite richness goes up the more metabolites are beneficial (S10.1 E, S10.2 E).

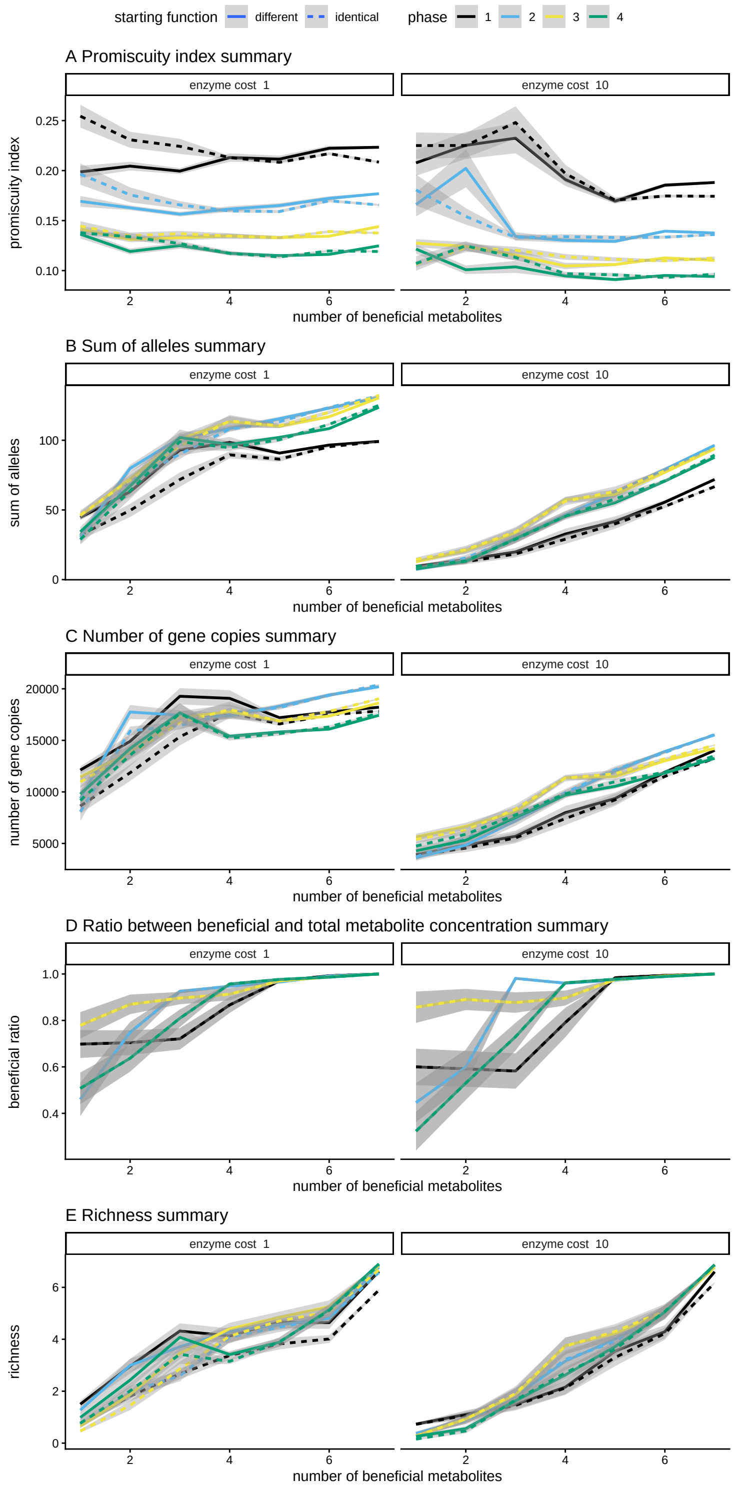


Figure S10.1: Summary figures of different diversity measures of the simulations with three modifications. The lines represent the mean value of a metric at the end of the phase for all simulations split out by starting function, enzyme costs, and phase, with number of beneficial metabolites on the x axis. The bands represent standard errors. Undefined values were left out of the analysis. A) the promiscuity index of the enzymes. High values indicate promiscuity, low values specialization. B) the SoA (sum of alleles across loci). C) the NoG (total number of gene copies across loci). D) the ratio of beneficial metabolites. E) the richness of derived metabolites.

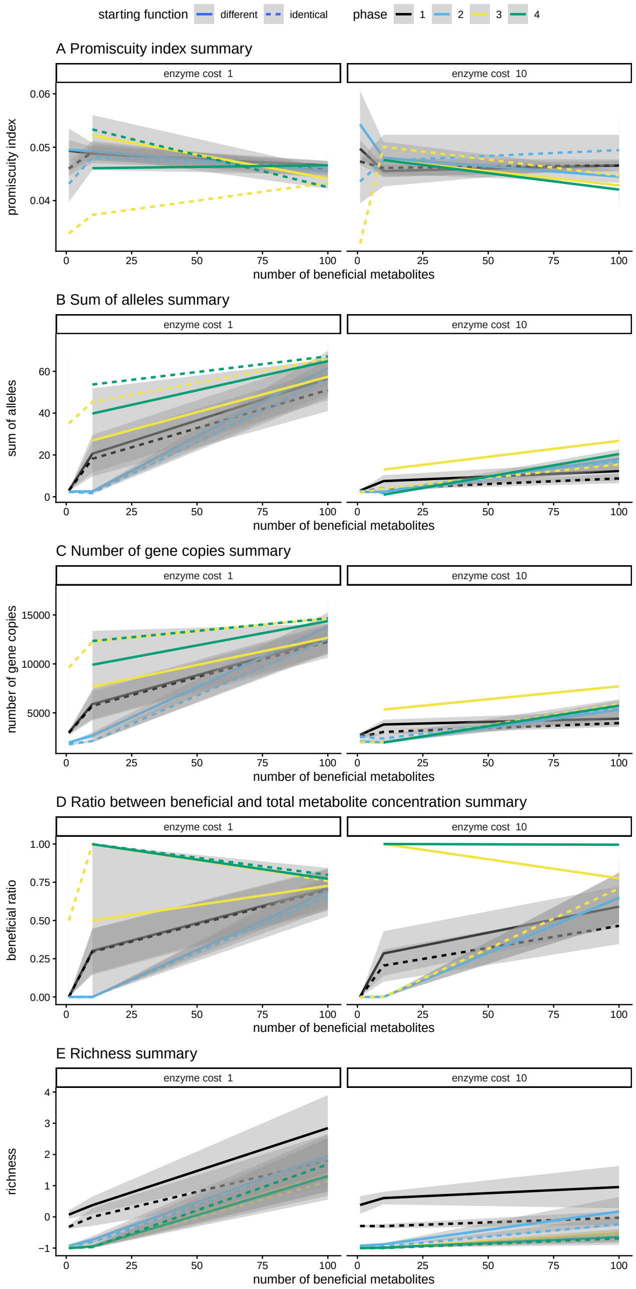


Figure S10.2: Summary figures of different diversity measures of the simulations with ten modifications. The lines represent the mean value of a metric at the end of the phase for all simulations split out by starting function, enzyme costs, and phase, with number of beneficial metabolites on the x axis. The bands represent standard errors. Undefined values were left out of the analysis. A) the promiscuity index of the enzymes. High values indicate promiscuity, low values specialization. B) the SoA (sum of alleles across loci). C) the NoG (total number of gene copies across loci). D) the ratio of beneficial metabolites. E) the richness of derived metabolites.
