## Supplementary material for "A metabolic model to investigate the evolution of chemodiversity": Simulation code and files: makefile

```
################################################################################
# Automatically-generated file. Do not edit!
################################################################################

-include ../makefile.init

RM := rm -rf

# All of the sources participating in the build are defined here
-include sources.mk
-include src/subdir.mk
-include subdir.mk
-include objects.mk

ifneq ($(MAKECMDGOALS),clean)
ifneq ($(strip $(CC_DEPS)),)
-include $(CC_DEPS)
endif
ifneq ($(strip $(C++_DEPS)),)
-include $(C++_DEPS)
endif
ifneq ($(strip $(C_UPPER_DEPS)),)
-include $(C_UPPER_DEPS)
endif
ifneq ($(strip $(CXX_DEPS)),)
-include $(CXX_DEPS)
endif
ifneq ($(strip $(CPP_DEPS)),)
-include $(CPP_DEPS)
endif
ifneq ($(strip $(C_DEPS)),)
-include $(C_DEPS)
endif
endif

-include ../makefile.defs

OPTIONAL_TOOL_DEPS := \
$(wildcard ../makefile.defs) \
$(wildcard ../makefile.init) \
$(wildcard ../makefile.targets) \


BUILD_ARTIFACT_NAME := program
BUILD_ARTIFACT_EXTENSION :=
BUILD_ARTIFACT_PREFIX :=
BUILD_ARTIFACT := $(BUILD_ARTIFACT_PREFIX)$(BUILD_ARTIFACT_NAME)$(if $(BUILD_ARTIFACT_EXTENSION),.$(BUILD_ARTIFACT_EXTENSION),)

# Add inputs and outputs from these tool invocations to the build variables 

# All Target
all: main-build

# Main-build Target
main-build: program

# Tool invocations
program: $(OBJS) $(USER_OBJS) makefile objects.mk $(OPTIONAL_TOOL_DEPS)
	@echo 'Building target: $@'
	@echo 'Invoking: GCC C++ Linker'
	g++  -o "program" $(OBJS) $(USER_OBJS) $(LIBS)
	@echo 'Finished building target: $@'
	@echo ' '

# Other Targets
clean:
	-$(RM) $(CC_DEPS)$(C++_DEPS)$(EXECUTABLES)$(C_UPPER_DEPS)$(CXX_DEPS)$(OBJS)$(CPP_DEPS)$(C_DEPS) program
	-@echo ' '

.PHONY: all clean dependents main-build

-include ../makefile.targets
```
