## Supplementary material for "A metabolic model to investigate the evolution of chemodiversity": Simulation code and files: Gene.cpp

```
/*
 * Gene.cpp
 *      Author: Frans
 */

#include "Gene.h"
#include "Input.h"
#include <iostream>
#include <fstream>

//extern std::ofstream outputLog;

void Gene::initializeGene(int index, const Input input)
{
    // Function to initialize the genes in the very fist step of the simulation. Genes created later are either copies of existing genes, or empty alleles.
	// Takes:
	// - index:						integer number used to create the unique gene ID. (for debugging purposes)
	// - dBaseTranscriptionRate:	initializes 'promotor': the transcription rate when not influenced by regulatory elements.
	// - iNumReactions:			 	total number of possible modifications to the base metabolite
	// - iMetabolicFunction:		location in the metabolite (represented by a binary vector) that the enzyme modifies
	// - vecCat:					vector of catalizing rates for every possible metabolite
	// - vecMichaelis:				as vecCat but michaelis constants
	// Puts out:
	// - Nothing, it initializes the member variables of a gene object at the start of the simulation.

	dConcentration = input.dBaseConcentration;
	//For coding genes
	cGeneID         = "C" + std::to_string(index);
	if (input.identicalFunction == true){
		iFunction = 0;
	} else {iFunction = index % input.iNumReactions;}
	vecKC.resize(std::pow(input.iNumReactions, 2));
	vecKM.resize(std::pow(input.iNumReactions, 2));
	vecKC = input.vecKCat[index];
	vecKM = input.vecKm[index];
}

Genome::Genome(bool initializeGenes, const Input input)
{
	// Function to initialize the genome of a plant.
	// Makes genomePairs, but no network.
	// This function should only be used during the initialization of the system.
	// Afterwards, genomes should be created by copying and modifying existing genomes.
	// Takes:
	// - boolean initializeGenes: 	Whether genes should be initialized (true) or whether the genome should be empty (false)
	// - input:						Input object provided by user.

	if (initializeGenes == true)
	{
		genomePairs.resize(input.iNumGenePairs); 					// Resize the genomePairs version of the genome

		for(int i = 0; i < input.iNumGenePairs; ++i)  // For each gene in the initial genome
		{
			genomePairs[i].resize(2);
			genomePairs[i][0].initializeGene(i, input);
			genomePairs[i][1].initializeGene(i, input);
		}
		listGenes();
	}
}

void Genome::listGenes()
{
	//function to add enzymes to the proteome, wrapper for initializeEnzyme
	//Takes:
	// - gene: A Gene to make an enzyme from
	//Puts out:
	// - Nothing, but a gene is added to the proteome affected by this function
	geneList.clear();
	for (int i = 0; (unsigned)i < genomePairs.size(); ++i)
	{
		for (int j = 0; j < 2; ++j)
		{
			if (genomePairs[i][j].proveExistence() == true)
			{
				geneList.push_back(genomePairs[i][j]);
			}
		}
	}

}

void Genome::duplicateGene(int iPair, int iAllele, int iNumGenes)
{
	// Function to duplicate a gene
	// Takes:
	// - iPair: 	position of the genepair the target gene belongs to in the genomePairs vector
	// - iAllele: 	position in the genepair the target gene belongs to.
	// Puts out:
	//  - Nothing, but it copies the target gene to the end of the genome and adds an empty gene with it.
	//    On a population level, whenever this function is called, all other plants must gain an empty gene pair to not mess up reproduction.

	Gene geneEmpty;
	geneEmpty.loseGene();
	std::vector<Gene> pairDuplicate = {genomePairs[iPair][iAllele], geneEmpty}; // vector of two genes, one of which is the duplicate, the other an empty gene.
	pairDuplicate[0].vecHistory.push_back(pairDuplicate[0].cGeneID);
	pairDuplicate[0].cGeneID = "C" + std::to_string(iNumGenes);
	genomePairs.push_back(pairDuplicate);
}
```
