## Supplementary material for "A metabolic model to investigate the evolution of chemodiversity": Simulation code and files: Gene.h

```
/*
 * Gene.h
 *      Author: Frans
 */

#ifndef GENE_H_
#define GENE_H_

#include <boost/graph/adj_list_serialize.hpp>

using namespace boost;

#include "Input.h"

class Gene
{
	//Every gene has a base transcription rate (governed by the promotor), can
	//mutate, be duplicated or lost.
public:
    //Function to initialize a gene.
	void initializeGene(int index, Input input);

    //Functions that return private values
    bool   proveExistence() const          			{return doesExist;}
    double outputConcentration() const 				{return dConcentration;}
    int    outputFunction() const          			{return iFunction;}
    std::vector<double> outputKCat() const          {return vecKC;}
    std::vector<double> outputMichaelis() const     {return vecKM;}

    //Functions to set gene variables to particular values
    void setFunction(int iNewFunction)   			{iFunction = iNewFunction;}
    void setConcentration(double dNewConc)			{dConcentration = dNewConc;}
    void setKCatVal(int iRate, double dValue)   	{vecKC[iRate] = dValue;}
    void setKMichaelisVal(int iRate, double dValue) {vecKM[iRate] = dValue;}
    void loseGene()									{doesExist = false; cGeneID = "empty";}

    std::string cGeneID;                       //Unique ID of the gene
	std::vector<std::string> vecHistory;       // Vector recording the ancestral genes of the gene

 private:
    //Both types of genes
	bool   doesExist = true;                   // Whether the aleles exist (becomes 'false' at gene loss)

	//Coding genes
	double dConcentration;                     // Enzyme concentration of the enzyme produced by the gene
	int    iFunction;                          // Metabolic function of the gene product (enzyme)
    std::vector<double> vecKC;                 // Vector of catalizing rates for each metabolite
    std::vector<double> vecKM;			       // As above, but for the michaelis constant

};


//Define the 'typeGeneNetwork' datatype which stores a genome.
typedef boost::adjacency_list<boost::vecS, boost::vecS,                    // Two vectors, one of vertices, one of edges
	    		                      boost::directedS,                    // Gene regulatory network as directed graph
									  Gene,                                // Vertices of type gene
									  boost::no_property> TypeGeneNetwork; // Edges for now without properties - idea to make edge weights depend on effect size.

class Genome
    // Every plant has a genome, which consists of a network of genes. These
    // genes can have enzyme-coding or regulatory functions.
{
	// Graph of genes and their interactions
public:
	// Function to initialize the genome.
	// Note: This should only be used for the first genes, later ones are copies.
	Genome(bool initializeGenes, const Input input);

	// Functions to output the genome
	std::vector<std::vector<Gene>>     outputGenePairs()  {return genomePairs;}                 // Return the genome as a vector gene pairs, both existant and empty
	std::vector<Gene>                  outputGeneList()   {return geneList;}                    // Return the genome as a vector of only the extant genes
	Gene outputGene(const int iPair, const int iAllele)   {return genomePairs[iPair][iAllele];} // Return a specific gene from the genome
	std::vector<std::string> IDlist;

	// Functions to operate on the genome
	void removeLocus(int iLocus)       	{genomePairs.erase(genomePairs.begin()+iLocus);}

	// Functions to change the genome after initialization
	void setGene(const Gene geneNew, int iPair, int iAllele)    		{genomePairs[iPair][iAllele] = geneNew;} // Replace a gene by another gene
	void addEmptyGenepair()								  				{Gene geneEmpty; geneEmpty.loseGene(); std::vector<Gene> pairEmpty = {geneEmpty, geneEmpty}; genomePairs.push_back(pairEmpty);}
	void duplicateGene(int iPair, int iAllele, int iNumGenes);                                                   // Duplicate a gene from the genome
	void setGenomePairs(const std::vector<std::vector<Gene>> genomeNew) {genomePairs = genomeNew; listGenes();}  //Replace the vector of gene pairs by another vector of gene pairs
	void listGenes();																							 //put gene pairs into a list without empty genes

private:
	std::vector<std::vector<Gene>>     genomePairs;   // Vector of all alleles of the genome, including ones that aren't there
	std::vector<Gene> geneList;   //List of enzyme-producing genes
};

#endif /* GENE_H_ */
```
