## Supplementary material for "A metabolic model to investigate the evolution of chemodiversity": Simulation code and files: Input.cpp

```
/*
 * Input.cpp
 *      Author: frans
 */

#include <iostream>
#include <fstream>
#include <random>

#include <boost/random.hpp>

#include "Input.h"

extern std::ofstream outputLog;

void Input::readInput(const std::string &inputFileName)
{
	// Function to read input from an input file
	// Should only be used once during the running of the program
	// Takes:
	// - inputFileName: The name of the input file to be read
	// Puts out:
	// - Nothing, but it initializes the input object that is used in various outher functions.

	// open the input file
	std::ifstream ifs(inputFileName);

	// trackers
	int 	iEnzyme				= 0;		//enzyme iterator
	int 	iMetabolite 		= 0;		//metabolite iterator
    int		iPhase				= 0;		//phase iterator
    int 	iPhaseTransition 	= 1;		//phase transition iterator
	bool 	knowNumInitialGenepairs = false;	//bool that checks that iNumInitialCodingGenes is known before reading particular parameters
	bool 	knowNumReactions 	= false;	//bool that checks that iNumReactions is known before reading particular parameters
	bool	knowNumPhases		= false;	//bool that checks that iNumPhases is known before reading particular parameters
	bool	knowNumGenerations	= false;	//bool that checks that iNumPhases is known before reading particular parameters
	extern 	std::mt19937_64 rng;			//random number generator
	std::string parId = "null";				//current parameter ID

	if(!ifs.is_open()) //check if data file can be opened
	{
		std::cerr << "error: unable to open data file\n";
		exit(EXIT_FAILURE);
	}

	for(;;)
	{
		// read a parameter name
		ifs >> parId;
		if(ifs.good()) //if parameter name read correctly
		{
			// Read in parameter values
			if(parId == "logFile")
			{
				ifs >> logFile;
				outputLog.open(logFile, std::ios::app);
				outputLog << "file opened" << std::endl;
			}
			//population parameters
			else if(parId == "iSeed")
			{
				ifs >> iSeed;
				rng.seed(iSeed);
			}
			else if (parId == "initialKSeed")
			{
				ifs >> iKSeed;
			}
			else if (parId == "fitnessSeed")
			{
				ifs >> iFSeed;
			}
			else if (parId == "zeroSeed")
			{
				ifs >> iZSeed;
			}
			else if(parId == "iPopSize")
			{
				ifs >> iPopSize;
			}
			else if(parId == "iNumGenerations")
			{
				ifs >> iNumGenerations;
			}
			//gene parameters
			else if(parId == "iNumGenePairs")
			{
				knowNumInitialGenepairs = true;
				ifs >> iNumGenePairs;
				if (knowNumReactions == true)
				{
					setKVectorSizes();
				}
			}
			else if (parId == "identicalFunction")
			{
				ifs >> identicalFunction;
			}
			else if(parId == "dBaseConcentration")
			{
				ifs >> dBaseConcentration;
			}
			//mutations
			else if(parId == "dLossOdds")
			{
				ifs >> dLossOdds;
			}
			else if(parId == "dDuplicationOdds")
			{
				ifs >> dDuplicationOdds;
			}
			else if(parId == "iMaxNumLoci")
			{
				ifs >> iMaxNumLoci;
			}
			else if(parId == "dMutationOdds")
			{
				ifs >> dMutationOdds;
			}
			else if(parId == "dMaxConcentration")
			{
				ifs >> dMaxConcentration;
			}
			else if(parId == "dMaxConcentrationChange")
			{
				ifs >> dMaxConcentrationChange;
			}
			else if(parId == "dKMu")
			{
				ifs >> dKMu;
			}
			else if(parId == "dKSigma")
			{
				ifs >> dKSigma;
			}
			else if(parId == "iNumReactions")
			{
				ifs >> iNumReactions;
				knowNumReactions = true;
				if (knowNumInitialGenepairs == true) //if both iNumReactions and iNumInitialCodingGenes are known
				{
					setKVectorSizes();
				}

				if (knowNumGenerations == true) //if both iNumReactions and iNumPhases are known
				{
					vecMetaboliteFitnessEffects.resize(iNumGenerations);
					for (int i = 0; i < iNumGenerations; ++i)
					{
						vecMetaboliteFitnessEffects[i].resize(std::pow(2, iNumReactions));
					}
				}
			}
			//kinetic parameters
			else if (parId == "setInitialK")
			{
				ifs >> setInitialK;
			}
			else if(parId == "vecKCat"+ std::to_string(iEnzyme) + std::to_string(iMetabolite))
			{
				if ((knowNumInitialGenepairs == true) & (knowNumReactions == true))
				{
					ifs >> vecKCat[iEnzyme][iMetabolite];
					if(iMetabolite < (std::pow(2, iNumReactions)-1))
					{
						++iMetabolite;
					}
					else
					{
						iMetabolite = 0; ++iEnzyme;
						if (iEnzyme == iNumGenePairs)
						{
							iEnzyme = 0;
						}
					}
				}
				else
				{
					std::cerr << "error: please place parameter '" <<  parId << "'after parameters iNumInitialCodingGenes and iNumReactions  " << inputFileName << '\n';
					exit(EXIT_FAILURE);
				}
			}
			else if(parId == "vecKm"+ std::to_string(iEnzyme) + std::to_string(iMetabolite))
			{
				if ((knowNumInitialGenepairs == true) & (knowNumReactions == true))
				{
					ifs >> vecKm[iEnzyme][iMetabolite];
					if(iMetabolite < (std::pow(2, iNumReactions)-1))
					{
						++iMetabolite;
					}
					else
					{
						iMetabolite = 0; ++iEnzyme;
						if (iEnzyme == iNumGenePairs)
						{
							iEnzyme = 0;
						}
					}
				}
				else
				{
					std::cerr << "error: please place parameter '" <<  parId << "'after parameters iNumInitialCodingGenes and iNumReactions  " << inputFileName << '\n';
					exit(EXIT_FAILURE);
				}
			}
			//other ODE parameters
			else if(parId == "dInitialMetConcentration")
			{
				ifs >> dInitialMetConcentration;
			}
			else if(parId == "dInflux")
			{
				ifs >> dInflux;
			}
			else if(parId == "dEfflux")
			{
				ifs >> dEfflux;
			}
			//ODE settings
			else if(parId == "dODELength")
			{
				ifs >> dODELength;
			}
			else if(parId == "ODEstepper")
			{
				ifs >> ODEstepper;
			}
			else if(parId == "dSmallestODEStepSize")
			{
				ifs >> dSmallestODEStepSize;
			}
			//fitness
			else if (parId == "fitnessHasEffect")
			{
				ifs >> fitnessHasEffect;
			}
			else if (parId == "iNumPhases")
			{
				ifs >> iNumPhases;
				knowNumPhases = true;
				vecPhaseTransitionTimes.resize(iNumPhases - 1);
				if (knowNumReactions == true) //if both iNumReactions and iNumPhases are known
				{
					vecMetaboliteFitnessEffects.resize(iNumGenerations);
					for (int i = 0; i < iNumGenerations; ++i)
					{
						vecMetaboliteFitnessEffects[i].resize(std::pow(2, iNumReactions));
					}
				}
			}
			else if (parId == "vecPhaseTransitionTimes" + std::to_string(iPhaseTransition) )
			{
				if (knowNumPhases == true)
				{
					ifs >> vecPhaseTransitionTimes[iPhaseTransition - 1];
					++iPhaseTransition;
				}
				else
				{
					std::cerr << "error: please place parameter '" <<  parId << "'after parameter iNumPhases " << inputFileName << '\n';
					exit(EXIT_FAILURE);
				}
			}
//			else if (parId == "scramblePhases")
//			{
//				ifs >> scramblePhases;
//			}
			else if(parId == "dFitnessCostPerEnzyme")
			{
				ifs >> dFitnessCostPerEnzyme;
			}
			else if (parId == "fitnessIsRandom")
			{
				ifs >> fitnessIsRandom;
			}

			else if(parId == "iFitnessEffectMet"+ std::to_string(iPhase) + std::to_string(iMetabolite) )
			{
				if ((knowNumReactions == true) & (knowNumGenerations == true))
				{
					if (iPhase >= iNumPhases)
					{
						std::cerr << "error: please place parameter '" <<  parId << "'after parameter iNumPhases " << inputFileName << '\n';
						exit(EXIT_FAILURE);
					} //error if trying to enter more values than there are phases
					ifs >> vecMetaboliteFitnessEffects[iPhase][iMetabolite];
					++iMetabolite;
					if (iMetabolite == pow(2, iNumReactions) )
					{
						iMetabolite = 0;
						++iPhase;
					}
				}
				else
				{
					std::cerr << "error: please place parameter '" <<  parId << "'after parameters iNumReactions and iNumPhases " << inputFileName << '\n';
					exit(EXIT_FAILURE);
				}
			}
			else if(parId == "dMinimumFitnessEffect")
			{
				ifs >> dMinimumFitnessEffect;
			}
			else if(parId == "dMaximumFitnessEffect")
			{
				ifs >> dMaximumFitnessEffect;
			}
			else if (parId == "iNumZeroFitness")
			{
				ifs >> iNumZeroFitness;
			}
			//output
			else if(parId == "outputODE")
			{
				ifs >> outputODE;
			}
			else if(parId == "outputMetabolites")
			{
				ifs >> outputMetabolites;
			}
			else if(parId == "outputFilesLocation")
			{
				ifs >> outputFilesLocation;
			}

			else if(parId == "iOutputInterval")
			{
				ifs >> iOutputInterval;
			}
			else if (parId == "file")
			{
				ifs >> iFileID;
			}
			else
			{
				std::cerr << "error: unknown parameter name '" <<  parId << "'in file " << inputFileName << '\n';
				exit(EXIT_FAILURE);
			}
//			outputLog << "parameter " << parId << " set to " << ifs << '\n';

		}
		else {
			outputLog << "last parameter read " << parId << std::endl;
			break; // to exit when EOF is reached or
		}
	}
	randomInitialK();
	createRandomFitnessAndMetaboliteCodes();
}

void Input::randomInitialK()
{
	std::mt19937_64 rngK; //
	rngK.seed(iKSeed);
	boost::random::lognormal_distribution<double>  amount(dKMu, dKSigma);
	if (setInitialK == false) //we set random values of K if this is TRUE even if values of K are supplied
	{
		for (int gene = 0; gene < iNumGenePairs; ++gene)
		{
			for (int metabolite = 0; (unsigned)metabolite < vecKCat[gene].size(); ++metabolite)
			{
				double dC = amount(rngK);
				double dM = amount(rngK);
				while (dC/dM > pow(10, 9)) //retry if kCat/kM would become higher than thermodynamically possible
				{
					dC = amount(rngK);
					dM = amount(rngK);
				}
				vecKm[gene][metabolite] = dM;
				vecKCat[gene][metabolite] = dC;
			}
		}
	}
//	outputLog << "initial K set" << std::endl;
}

void Input::setKVectorSizes()
{
	vecKCat.resize(iNumGenePairs);
	vecKm.resize(iNumGenePairs);
	for (int i = 0; i < (iNumGenePairs); ++ i)
	{
		vecKCat[i].resize(std::pow(2, iNumReactions));
		vecKm[i].resize(std::pow(2, iNumReactions));
	}
//	outputLog << "vector sizes set " << std::endl;
}

void Input::createRandomFitnessAndMetaboliteCodes()
{
	// Open file to record the randomly generated fitness values
	std::ofstream fitnessVals;
	fitnessVals.open("random_fitness.csv", std::ios_base::app);
	//prepare random distributions
	boost::uniform_real<> dist(dMinimumFitnessEffect, dMaximumFitnessEffect); //random fitness
	std::mt19937_64 rngFit; //random fitness rng
	rngFit.seed(iFSeed);
	std::mt19937_64 rngZero; //random zero fitness rng
	rngZero.seed(iZSeed);
//	std::vector<int> vecPhaseScramble(iNumPhases); //vector of positions to randomize phase order
//	std::iota(std::begin(vecPhaseScramble), std::end(vecPhaseScramble), 0); //every value of the vector is its position
//	if (scramblePhases == true) {std::shuffle(std::begin(vecPhaseScramble), std::end(vecPhaseScramble), rngPhase);} //shuffle the values if appropriate: now it can be used to shuffle the phases

	vecMetaboliteCodes.resize(std::pow(2, iNumReactions)); //vector of vectors to recieve the metabolite codes

	for (int i = 0; i < std::pow(2, iNumReactions); ++i) //for every metabolite
	{
		//initialize metabolite codes
		vecMetaboliteCodes[i].resize(iNumReactions); //ready the vector to recieve the binary code for the metabolite
		int j = i;
		for (int k = (iNumReactions - 1); k > -1; --k) //for every position in the vector
		{
			if(j >= std::pow(2, k)) //if the decimal value of the metabolite (or remnant) is higher than the decimal value of the binary bit
			{
				vecMetaboliteCodes[i][k] = 1;//add a 1
				j -= std::pow(2, k); //and subtract the value of the bit, to then evaluate the remnant for the next bit
			}
			else
			{
				vecMetaboliteCodes[i][k] = 0;//else add a 0 and move on
			}
		}
		//create random fitness effects (and possibly randomize phase order, only works for random fitness)
		if (fitnessIsRandom == true){
			if (i == 0) //if most basic metabolite
			{
				for (int g = 0; g < iNumGenerations; ++g) //in every phase
				{
					vecMetaboliteFitnessEffects[g][i] = 0; //fitness value is 0
				}
			}
			else
			{
				for (int g = 0; g < iNumGenerations; ++g)
				{
					vecMetaboliteFitnessEffects[g][i] = dist(rngFit); //put the random fitness value in the correct phase and metabolite
				}
			}
		}
	}
	//set as many values to zero each phase as iNumZeroFitness
//	outputLog << "metabolites set- vector length " << vecMetaboliteFitnessEffects[1].size() << " " << iNumZeroFitness << std::endl;
	int iPhase = 0;
	std::vector<int> vecScramble(std::pow(2, iNumReactions)-1); //Vector of size metabolites -1 (we exclude the basic metabolite 0)
	std::iota(std::begin(vecScramble), std::end(vecScramble), 1); //initialize starting at 1
	for (int g = 0; g < iNumGenerations; ++g){
//		outputLog << "test 1 " << std::endl;
		if (g == 0){
			std::shuffle(std::begin(vecScramble), std::end(vecScramble), rngZero); //shuffle)
		}
//		outputLog << "test 2 " << g << std::endl;
//		outputLog << vecPhaseTransitionTimes[iPhase] << std::endl;
//		outputLog << rngZero << std::endl;
		if (g == vecPhaseTransitionTimes[iPhase]){
			++iPhase;
			std::shuffle(std::begin(vecScramble), std::end(vecScramble), rngZero); //shuffle)
		}
//		outputLog << "test 3 " << std::endl;
		for (int i = 0; i < iNumZeroFitness; ++i) //for as many metabolites are to be set to zero
		{
			vecMetaboliteFitnessEffects[g][vecScramble[i]] = 0;
		}
//		outputLog << "test 4 " << std::endl;
	}
	outputLog << "zeroes set " << std::endl;
	iPhase = 0;
	for (int g = 0; g < iNumGenerations; ++g) //enter final fitness values into file
	{
		if (g == vecPhaseTransitionTimes[iPhase] || g == 0){
			if (g != 0){++iPhase;}
			for (int i = 0; i < std::pow(2, iNumReactions); ++i)
			{
				fitnessVals << iSeed << ", " << iFSeed << ", " << iPhase << ", " << iZSeed << ", " << g << ", " << i << ", " << vecMetaboliteFitnessEffects[g][i] << std::endl;
			}
		}
	}
//	outputLog << "metabolite codes set" << std::endl;
}
```
