## Supplementary material for "A metabolic model to investigate the evolution of chemodiversity": Simulation code and files: Input.h

```
/*
 * Input.h
 *      Author: Frans
 */

#ifndef INPUT_H_
#define INPUT_H_

struct Input
{
	 std::string logFile 	= "log.txt";					// Filename of the log file

//population parameters
	 int    iSeed;
	 int    iKSeed = 1;										// Random seed if the initial kinetic parameters are picked from a random distribution
	 int  	iFSeed;											//random fitness seed
	 int  	iZSeed;											//random zero zeed
	 int    iPopSize = 0;									// Population size
	 int    iNumGenerations = 0;							// Number of generations

//gene parameters
	 int    iNumGenePairs = 0;   							// Initial number of gene pairs
	 bool   identicalFunction = false;                      // If multiple coding gene pairs: Should these genes all have the same function or different ones?
	 double dRegulatoryEffect = 0;                         	// Regulatory effect (up, down) of initial regulatory genes
	 double dBaseConcentration = 0;						// Initial transcription rate of genes at start of simulation

//mutations
	 double dLossOdds = 0;									// Odds of loss
	 double dDuplicationOdds = 0;							// Odds of gene duplication
	 int 	iMaxNumLoci = 100;                              // Maximum length of the genomePairs vector
	 double dMutationOdds = 0;							    // Odds of mutations (apart from duplication and loss) happening
	 double dMaxConcentration = 0;							// Maximum enzyme concentration
	 double dMaxConcentrationChange = 0;					// Maximum change of enzyme concentration of gene
	 double dKMu;  											// Mu of the lognormal distribution that randomly determines Ks
	 double dKSigma;										// Sigma of the lognormal distribution that
	 int    iNumReactions = 0;                          	// Number of possible modifications to a metabolite

//metabolism
	 //general metabolite parameters
	 //kinetic parameters
	 bool setInitialK = true;								// Boolean of whether the initial kinetic parameters are set  manually or picked from a random distribution
	 std::vector<std::vector<double>> vecKCat {{0}};   		// Vector of catalizing rates for each metabbolite
	 std::vector<std::vector<double>> vecKm {{0}}; 			// Vector of the michaelis-menten constant
	 //other ODE parameters
	 double dInitialMetConcentration = 0;					// Initial concentration of the unmodified metabolite
	 double dInflux = 0;                                    // Rate of influx (production?) of unmodified metabolite into metabolic system
	 double dEfflux = 0;                                    // Rate of efflux of all metabolites from metabolic system
	 std::vector<std::vector<bool>> 	vecMetaboliteCodes {{0}}; 		// Vector of codes of possible metabolites
	 //ODE settings
	 double dODELength = 0; 								// Length of the ODE- number of timesteps
	 std::string ODEstepper = "runge-kutta";				// Stepper used to solve the ODE
	 double dSmallestODEStepSize = 0.0001;                  // Smallest step size the ODE can take before returning everything as 0.

//fitness
	 bool fitnessHasEffect = 1;                             // Boolean whether the fitness has effect on reproduction
	 //phase parameters
	 int	iNumPhases = 0;									// Number of phases under particular fitness effects in the simulation
	 std::vector<double> vecPhaseTransitionTimes = {0};				// Vector of times when the phase transitions take place
//	 bool	scramblePhases = false;							// boolean whether the phases should be scrabled or not
	 //non-metabolite effects
	 double dFitnessCostPerEnzyme = 0;                      // Fitness cost of every enzyme (will be multiplied with enzyme concentration)
	 //metabolite effects
	 bool   fitnessIsRandom = false;							// Boolean whether fitness is random or set by user
	 std::vector<std::vector<double>> 	vecMetaboliteFitnessEffects; 	// Vector (phases) of vectors (metabolites) of fitness effects of possible metabolites
	 double dMinimumFitnessEffect = 1;						// Minimum fitness effect of a metabolite
	 double dMaximumFitnessEffect = 1;						// Maximum fitness effect of a metabolite
	 int 	iNumZeroFitness = 0;							// Number of random metabolites that should have their fitness set to zero

//output
	 bool 	outputODE = true;								// Determines whether the ODE gets put out
	 bool 	outputMetabolites = true;						// Determines whether metabolite concentrations get put out
	 std::string outputFilesLocation = "output";			// File location of the output files
	 int 	iOutputInterval = 1;							// Number of generations that should be between writing to the output files
	 int 	iFileID = 0;										// File ID

	 void readInput(const std::string &inputFileName);
	 void randomInitialK();
	 void setKVectorSizes();
	 void createRandomFitnessAndMetaboliteCodes();
};

#endif /* INPUT_H_ */
```
