## Supplementary material for "A metabolic model to investigate the evolution of chemodiversity": Simulation code and files: Metabolism.cpp

```
/*
 * Metabolism.cpp
 *      Author: Frans
 */

#include <fstream>
#include <boost/numeric/odeint.hpp>

#include "Metabolism.h"

//extern std::ofstream outputLog;
std::ofstream outputFileStream;

void writeODESystem( const state_type &x , double t)
{
	// Function to write the state of the ODE to the output file
	// Takes:
	// - x:	Vector of values that describes the state of the ODE in the current timestep
	// - t: Value of the current timestep
	// Puts out:
	// - Nothing, but it writes the state of the ODE to the output file.

	outputFileStream << t << ',' ;                // Write the timestep
	for (int i = 0; (unsigned)i < x.size(); ++i)  // Write the value of each part of the ODE
	{
		outputFileStream << x[i] << ',';
	}
	outputFileStream << "\n"; 				      // End the line
}

std::vector<double> runMetabolism(const std::vector<Gene>& geneList, const std::vector<int>& relevants, const std::vector<int>& decreaseFunctionValue, const std::vector<std::vector<bool>>& vecMetaboliteCodes, const int iGeneration, const int iPlant, const Input& input, double dStepSize)
{
	// Function to run the metabolism of the plant.
	// Takes:
    // - genome plant:	The genome of the plant
	// - filenumber:	The number that marks the specific output file of the ODE
	// - input:			A struct of user-defined static input values
	// - dStepSize:		Step size of the odesolver
	// Puts out:
	// - The end state of the ODE as a vector.
	// - Informally, the states of the ODE through time.

	// Prepare the parameters for the ODE
	// Struct of static variables
	odeParam 					  tStruct;

	tStruct.dGamma  			= input.dInflux;
	tStruct.dEpsilon 			= input.dEfflux;
	tStruct.iGenomeSize 	  	= geneList.size();
	tStruct.iNumReactions 		= input.iNumReactions;
	tStruct.vecMetaboliteCodes  = vecMetaboliteCodes;

	// Vector of metabolites that should be skipped - false for now, then set to true for all metabolites that cannot be made
	tStruct.skips.resize(std::pow(2, input.iNumReactions), false);
	std::vector<bool> functionPresent(input.iNumReactions, false);
	tStruct.KCat.resize(tStruct.iGenomeSize); 				//matrix of KCats. Every row an enzyme, every column a metabolite
	tStruct.KM.resize(tStruct.iGenomeSize);   				//matrix of KMs. Every row an enzyme, every column a metabolite
	tStruct.concentration.resize(tStruct.iGenomeSize);     	//vector of concentrations of each enzyme
	tStruct.function.resize(tStruct.iGenomeSize);          	//vector of functions of each enzyme
	tStruct.relevants = relevants;
	// first determine which functions are there, and copy gene information in a more accessible way

	for (int i = 0; i < tStruct.iGenomeSize; ++i)
	{
		tStruct.KCat[i].resize(relevants.size());
		tStruct.KM[i].resize(relevants.size());
		std::vector<double> KCat = geneList[i].outputKCat();
		std::vector<double> KM   = geneList[i].outputMichaelis();
		for (int j = 0; (unsigned)j < relevants.size(); ++j)
		{
			tStruct.KCat[i][j] = KCat[relevants[j]];
			tStruct.KM[i][j]   = KM[relevants[j]];
		}
		tStruct.concentration[i] 	= geneList[i].outputConcentration();
		tStruct.function[i] 		= geneList[i].outputFunction();
		tStruct.function[i] 		-= decreaseFunctionValue[tStruct.function[i]];
//		outputLog << "function " << tStruct.function[i] << std::endl;
	}

	state_type x;
	// Vector of the state of the Metabolites present, initialized as all starting at 0 for now
	x.resize(tStruct.relevants.size(), 0.0);
	x[0] = input.dInitialMetConcentration; // Initial concentration of unmodified Metabolite

    // Prepare output file
    if (input.outputODE == true)
    {
    	std::string string1  = "/data_metabolism";
    	std::string string2  = ".csv";
    	std::string filename = input.outputFilesLocation + string1 + "g" + std::to_string(iGeneration) + "p" + std::to_string(iPlant) + string2;
    	outputFileStream.open(filename);
    	// Create column headers in the output file
    	outputFileStream << "t" << ',' ;             					// Timestep
    	for (int i = 0; (unsigned)i < tStruct.relevants.size(); ++i)	// Metabolites
    	{
    		outputFileStream << "M";
    		std::vector<bool> code = input.vecMetaboliteCodes[relevants[i]];
    		for (int j = 0; (unsigned)j < code.size(); ++j)
    		{
    			outputFileStream << code[j];
    		}
    		outputFileStream << ',';
    	}
    	outputFileStream << std::endl;
    }

    // Define the stepper for the integration
    boost::numeric::odeint::runge_kutta4< state_type > stepper;
    if (input.ODEstepper == "runge-kutta") {}
    else if (input.ODEstepper == "euler")			{boost::numeric::odeint::euler< state_type > stepper;}
    else if (input.ODEstepper == "bulirsch-stoer")	{boost::numeric::odeint::bulirsch_stoer< state_type > stepper;}
    else if (input.ODEstepper == "runge-kutta-54")  {boost::numeric::odeint::runge_kutta_cash_karp54< state_type > stepper;}
    else {std::cerr << "error: invalid stepper '" <<  input.ODEstepper << '\n';}

    // Create the ODE with all information from tStruct
    ode ho(tStruct);
    // Integrate/run the ODE
   if (input.outputODE == true)
    {
    	boost::numeric::odeint::integrate_const(stepper, ho, x, 0.0, input.dODELength, dStepSize, writeODESystem);
    	outputFileStream.close(); // Close the output file
    }
    else {
    	boost::numeric::odeint::integrate_const(stepper, ho, x, 0.0, input.dODELength, dStepSize);
    }

    std::vector<double> out;
    out = {x.begin(), x.begin() + x.size()};

    return out;
}
```
