## Supplementary material for "A metabolic model to investigate the evolution of chemodiversity": Simulation code and files: Metabolism.h

```
/*
 * Metabolism.h
 *      Author: Frans
 */

#ifndef METABOLISM_H_
#define METABOLISM_H_

#include "Gene.h"

#include <iostream>
#include <fstream>
extern std::ofstream outputLog;

typedef std::vector< double > state_type;

// Functions to run the metabolism
std::vector<double> runMetabolism(const std::vector<Gene>& geneList, const std::vector<int>& relevants, const std::vector<int>& decreaseFunctionValue, const std::vector<std::vector<bool>>& vecMetaboliteCodes, const int iGeneration, const int iPlant, const Input& input, double dStepSize);
void 				writeODESystem(const state_type &x , double t);

struct odeParam
{
	// Structure that holds parameters for the ODE.
//    std::vector<Gene> K; 	// From the genome we need the catalyzing rates and michaelist constants, which is why the genome is called K here.
    std::vector<std::vector<double>> KCat; //matrix of KCats. Every row an enzyme, every column a metabolite
    std::vector<std::vector<double>> KM;   //matrix of KMs. Every row an enzyme, every column a metabolite
    std::vector<double> concentration;     //vector of concentrations of each enzyme
    std::vector<double> function;          //vector of functions of each enzyme
    std::vector<bool> skips;// vector of booleans: true if the metabolite cannot be created in the current plant, false if they can. (speeds up the ODE if some gene functions are missing)
    std::vector<int> relevants; // vector of the metabolites that can actually be created
    double dGamma;    		// Influx into the system of unmodified metabolite
    double dEpsilon;		// Efflux from the system of all metabolites
    int iGenomeSize;
    int iNumReactions;
    std::vector<std::vector<bool>> vecMetaboliteCodes;
};


class ode
{
	// Structure that holds the ODE
	struct odeParam K; // The structure that holds information on the static parameters of the ODE
	public:
	    ode(const struct odeParam myParams) : K(myParams) {} // Function to load parameters into the ode class

    // System of ODEs
	void operator()( const state_type &x , state_type &dxdt , const double /* t */)
    {
//		for (int i = 0; i < std::pow(2, K.iNumReactions); ++i)//metabolite dxdt[i]: one for each metabolite.

		for (int i = 0; (unsigned)i < x.size(); ++i)
		{
			dxdt[i] = 0;
//			outputLog << "dxdt[" << i << "]=";
			if (i == 0) //if initial metabolite add influx
			{
				dxdt[i] = dxdt[i] + K.dGamma;
//				outputLog << dxdt[i] << " " << K.dGamma << " ";
			}
			if (K.iGenomeSize) //if there's genes
			{
				for(int j = 0; j < K.iGenomeSize; ++j)//Per gene
				{
					if (K.vecMetaboliteCodes[i][K.function[j]] == 0) // if current metabolite can be modified by current gene
					{
						dxdt[i] = dxdt[i] - ((K.KCat[j][i] * x[i])/ (K.KM[j][i] + x[i]) * K.concentration[j]);//- michaelis menten of the current metabolite
//						outputLog << "((K.KCat[" << j << "][K.relevants[" << i << "]] * x[" << i << "])/ (K.KM[" << j << "][K.relevants[" << i << "]] + x[" << i << "]) * K.concentration[" << j << "])";
					}else //if the gene can produce this metabolite
					{
						dxdt[i] = dxdt[i] + ((K.KCat[j][i - std::pow(2, K.function[j])] * x[i - std::pow(2, K.function[j])])/ (K.KM[j][i - std::pow(2, K.function[j])] + x[i - std::pow(2, K.function[j])]) * K.concentration[j]);//+ michaelis menten of the precursor
//						outputLog << "";
					}
				}
			}
			dxdt[i] = dxdt[i] -K.dEpsilon * x[i]; //remove efflux
//			outputLog << dxdt[i] << " " << x[i] << " ";
//			outputLog << -K.dEpsilon << "*" << x[i] << std::endl;
		}
//		outputLog << std::endl;
    }
};

#endif /* METABOLISM_H_ */
```
