## Supplementary material for "A metabolic model to investigate the evolution of chemodiversity": Simulation code and files: Plant.cpp

```
/*
 * Plant.cpp
 *      Author: Frans
 */

#include <fstream>

#include "Plant.h"

extern std::ofstream outputLog;

Plant::Plant(bool initializeGenome, const Input input) : genomePlant(initializeGenome, input){}

void Plant::calculatePlantFitness(int iGeneration, const Input& input)
{
	// Function to calculate the fitness of a single plant from the fitness effects of gene transcription, enzyme production, and metabolites.
	// Input:
	// - input: Structure of user-defined variables
	// - iPhase: The phase the simulation is in. Determines metabolite fitness effects, which can change over time
	// Puts out:
	// Nothing, but the fitness of the plant (dFitnessPlant) is calculated.

	dFitnessPlant = 0; //Set Fitness to 0

	std::vector<Gene> geneList = genomePlant.outputGeneList();

	for (int i = 0; (unsigned)i < geneList.size(); ++i)
	{
		dFitnessPlant -= geneList[i].outputConcentration() * input.dFitnessCostPerEnzyme; 	//Fitness effect of enzymes
	}

//	if (iGeneration > 240 )
//	{
//		outputLog << "final concentrations: ";
//		for (int i = 0; (unsigned)i < vecMetaboliteConcentrations.size(); ++i)
//		{
//			outputLog << vecMetaboliteConcentrations[i] << ", ";
//		}
//		outputLog << std::endl;
//	}

	for (int i = 0; (unsigned)i < std::pow(2, input.iNumReactions); ++i)
	{
		dFitnessPlant += vecMetaboliteConcentrations[i] * input.vecMetaboliteFitnessEffects[iGeneration][i];			//Fitness effect of metabolites - calculated by runPlantMetabolism()
	}
	if(dFitnessPlant < 0){outputLog << "plant fitness " << dFitnessPlant << ", set to 0" << std::endl; dFitnessPlant = 0;}
}

void Plant::runPlantMetabolism(int iGeneration, int iPlant, const Input& input)
{
	// Function to run the plant metabolism
	// Input:
	// - iGeneration: 	The generation the simulation is in
	// - iPlant: 		The plant ID in the population
	// - input:			The input object
	// Returns:
	// Nothing, but the metabolism is run and the metabolite concentrations are calculated.

	double dStepSize = 1;
	bool   anyErr;

	// Vector of metabolites that should be skipped - false for now, then set to true for all metabolites that cannot be made
	std::vector<bool>    skips(std::pow(2, input.iNumReactions), false);
	std::vector<bool>    functionPresent(input.iNumReactions, false);
	std::vector<int>     relevants;
	std::vector<double>  vecMetaboliteConcentrationsTemp;
	std::vector<int>  decreaseFunctionValue(input.iNumReactions, 0);
	// first determine which functions are there, and copy gene information in a more accessible way

	std::vector<Gene> geneList = genomePlant.outputGeneList();

	for (int i = 0; (unsigned)i < geneList.size(); ++i) //list which gene functions are present
	{
		functionPresent[geneList[i].outputFunction()] = true;
	}
	for (int i = 0; (unsigned)i < std::pow(2, input.iNumReactions); ++i) //list which metabolites can be made
	{
		std::vector<bool> code = input.vecMetaboliteCodes[i];
		for (int j = 0; j < input.iNumReactions; ++j)
		{
			if ((code[j] == 1) & (functionPresent[j] != true))
			{
				skips[i] = true;
			}
		}
		if (skips[i] == false){
		//			outputLog << i << " ";
			relevants.push_back(i);
		}
	}
	int nrRelevants = relevants.size();

	int decrease = 0;
	for (int i = 0; i < input.iNumReactions; ++i) //list how much each 'function' needs to be decreased considering the missing metabolites
	{
		if (functionPresent[i] != true)
		{
			++decrease;
		}
		decreaseFunctionValue[i] = decrease;
	}

	std::vector<std::vector<bool>> vecMetaboliteCodes;
	vecMetaboliteCodes.resize(nrRelevants);
	for (int i = 0; i < nrRelevants; ++i)
	{
		vecMetaboliteCodes[i].reserve(input.iNumReactions-decrease);
		for (int j = 0; j < input.iNumReactions; ++j)
		{
			if (functionPresent[j] == true)
			{
				vecMetaboliteCodes[i].push_back(input.vecMetaboliteCodes[relevants[i]][j]);
			}
		}
	}

//	outputLog << "ready to run metabolism " << iPlant << std::endl;

	do
	{
		anyErr = false;
		vecMetaboliteConcentrationsTemp = runMetabolism(geneList, relevants, decreaseFunctionValue, vecMetaboliteCodes, iGeneration, iPlant, input, dStepSize); //run the metabolism
	    for(int i = 0; i < nrRelevants; ++i) //check whether the ODE collapses to nan and needs a smaller step size.
	    {
		    if (std::isnan(vecMetaboliteConcentrationsTemp[i]) || vecMetaboliteConcentrationsTemp[i] < 0)
		    {
		    	outputLog << "offending values ";
	    		for (int j = 0; j < nrRelevants; ++j) //record what values causes issues
	    		{
	    			outputLog << vecMetaboliteConcentrationsTemp[j] << ", ";
	    		}
	    		outputLog << std::endl;
		    	if (dStepSize <= input.dSmallestODEStepSize)
		    	{
		    		outputLog << "ODE cannot be resolved. Generation " << iGeneration << " Plant " << iPlant << " Metabolite " << i << ": " << vecMetaboliteConcentrationsTemp[i] << " - all values set to 0" << std::endl;
		    		for (int j = 0; (unsigned)j < vecMetaboliteConcentrationsTemp.size(); ++j)
		    		{
		    			vecMetaboliteConcentrationsTemp[j] = 0;
		    		}
		    	} else
		    	{
		    		anyErr = true;

		    		dStepSize /= 10;
		    		outputLog << "Generation " << iGeneration << " Plant " << iPlant << " Stepsize decreased to " << dStepSize << std::endl;
		    		break;
		    	}
		    }

	    }

	    vecMetaboliteConcentrations.resize(std::pow(2, input.iNumReactions), 0.0);
	    for (int i = 0; i < nrRelevants; ++i)
	    {
	    	vecMetaboliteConcentrations[relevants[i]] = vecMetaboliteConcentrationsTemp[i];
	    }

	} while(anyErr == true); //If there are any nan, run the loop again with a smaller stepsize
}
```
