## Supplementary material for "A metabolic model to investigate the evolution of chemodiversity": Simulation code and files: Plant.h

```
/*
 * Plant.h
 *      Author: Frans
 */

#ifndef PLANT_H_
#define PLANT_H_

#include "Metabolism.h"

//extern std::ofstream outputLog;

class Plant{
public:
	         Plant(bool initializeGenome, const Input input);

	// Functions  to put out member variables
    Genome   outputPlantGenome()           			{return genomePlant;}
    double   outputPlantFitness()     	   			{return dFitnessPlant;}
    std::vector<double> 	outputMetaboliteConcentrations()		 			{return vecMetaboliteConcentrations;}
    void setMetaboliteConcentrations(std::vector<double> vecConcentrations)		{vecMetaboliteConcentrations = vecConcentrations;}


    // Functions to pass down changes to the genome
    void     setGenomePairs(std::vector<std::vector<Gene>> newGenome) 	{genomePlant.setGenomePairs(newGenome);}
    void     setGene(Gene gene, int iPair, int iAlele)                  {genomePlant.setGene(gene, iPair, iAlele);}
    void     listGenes()                                                {genomePlant.listGenes();}
    void     removeLocus(int iLocation)                                 {genomePlant.removeLocus(iLocation);}
    void 	 addEmptyGenepair()            								{genomePlant.addEmptyGenepair();}
    void 	 duplicateGene(int iPair, int iGene, int iNumGenes) 		{genomePlant.duplicateGene(iPair, iGene, iNumGenes);}

    // Function that update the plant
    void     runPlantMetabolism(int iGeneration, int iPlant, const Input& input);
    void     calculatePlantFitness(int iGeneration, const Input& input);

private:
	Genome   genomePlant;
	double   dFitnessPlant = 0;
	std::vector<double> vecMetaboliteConcentrations; //Output of runPlantMetabolism - the concentrations of the different metabolites after the ODE has run.
};


#endif /* PLANT_H_ */
```
