## Supplementary material for "A metabolic model to investigate the evolution of chemodiversity": Simulation code and files: Population.cpp

```
/*
 * Population.cpp
 *      Author: Frans
 */
#include "Population.h"

#include <random>
#include <sys/stat.h>
#include <boost/random.hpp>


extern std::ofstream outputLog;
struct stat info;

PlantPopulation::PlantPopulation(const Input input)
{
	// Function to initialize the plant population at the start of the timeseries
	// Takes:
	// - input: Input read from user-provided input file
	// Puts out:
	// - Nothing, but the plant population is initialized.

	outputLog << "initialize first plant" << std::endl;
	Plant plantFirst(true, input);               // Initialize a plant that will be cloned to make up the whole population
	vPlants.reserve(input.iPopSize);
	for(int i = 0; i < input.iPopSize; ++i)      // Copy the plant into the vector of plants
	{
		vPlants.push_back(plantFirst);
	}
	vPopulationFitnesses.resize(input.iPopSize);
	outputLog << "first population set" << std::endl;
	//Update the population
	update(0, input);

	outputLog << "first population metabolisms run" << std::endl;
	//Initialize the output files: open files and add headers
	if (mkdir(input.outputFilesLocation.c_str(), 0777) == -1) //check if output folder exists- if yes, return error and end program
	{
		std::cerr << "Error :  " << strerror(errno) << " " << input.outputFilesLocation.c_str() << std::endl;
		exit(EXIT_FAILURE);
	}

//	phylogenyFile.open(input.outputFilesLocation + "/phylogeny"); //open output file for registering every mutation
//	phylogenyFile << "'timestep', 'parent gene', 'descendant gene', 'changed parameter', 'new value'" << std::endl;

	iNumGenes = input.iNumGenePairs; //Initialize the code of the next gene that will come into existence from mutation

	metaboliteFile.open(input.outputFilesLocation + "/metabolite"); //open output file for registering metabolite concentrations at end of ODE
	metaboliteFile << "'timestep', 'plant', ";
	for (int i = 0; (unsigned)i < std::pow(2, input.iNumReactions); ++i)
	{
		for (int j = 0; j < input.iNumReactions; ++j)
		{
			metaboliteFile << input.vecMetaboliteCodes[i][j];
		}
		metaboliteFile << ", ";
	}
	metaboliteFile << std::endl;

	richnessFile.open(input.outputFilesLocation + "/richness"); //open output file for registering every mutation
	richnessFile << "'timestep', 'plant', 'richness'" << std::endl;

	ratioFile.open(input.outputFilesLocation + "/ratio"); //open output file for registering every mutation
	ratioFile << "'timestep', 'plant', 'ratio'" << std::endl;

	genomeFile.open(input.outputFilesLocation + "/genome"); //open output file for registering genes of every plant
	genomeFile << "'timestep', ";
	for (int i = 0; i < input.iPopSize; ++i){
		genomeFile << i << " ,";
	}
	genomeFile << std::endl;

	fitnessFile.open(input.outputFilesLocation + "/fitness"); //open output file for registering fitness of every plant
	fitnessFile << "'timestep', ";
	for (int i = 0; i < input.iPopSize; ++i){
		fitnessFile << i << " ,";
	}
	fitnessFile << std::endl;

	geneFile.open(input.outputFilesLocation + "/genes"); //open output file for registering parameters of every gene
	geneFile << "'timestep', 'ID', 'ancestral IDs', 'Function', 'TranscriptionRate', ";
	for (int i = 0; i < std::pow(2, input.iNumReactions); ++i)
	{
		geneFile << "Kcat";
		for (int j = 0; j < input.iNumReactions; ++j)
		{
			geneFile << input.vecMetaboliteCodes[i][j];
		}
		geneFile << ", ";
	}
	for (int i = 0; (unsigned)i < std::pow(2, input.iNumReactions); ++i)
	{
		geneFile << "KM";
		for (int j = 0; j < input.iNumReactions; ++j)
		{
			geneFile << input.vecMetaboliteCodes[i][j];
		}
		geneFile << ", ";
	}
	geneFile << std::endl;

	outputLog << "output initialized" << std::endl;
	for (int gene = 0; gene < input.iNumGenePairs; ++gene) //record the initial genes
	{
		recordGene(input, 0, vPlants[0].outputPlantGenome().outputGene(gene, 0));
	}

	iNumLociLast = input.iNumGenePairs;

	outputLog << "first genes recorded" << std::endl;
	outputStateToFiles(input.outputFilesLocation, input, 0);
	outputLog << "first output recorded" << std::endl;
}

void PlantPopulation::calculatePopulationFitness(int iGeneration, const Input input)
{
	// Function to calculate the fitness of the plants in the population
	// Takes:
	// - input: Input read from user-provided input file
	// - iPhase: The phase the simulation is in. Determines metabolite fitness effects, which can change over time
	// Puts out:
	// - Nothing, but the fitness of each plant in the population is calculated.

	for (int i = 0; i < input.iPopSize; ++i) //For each plant
	{
//		if (iGeneration > 240 )
//		{
//			outputLog << "plant: " << i << std::endl;
//		}
		vPlants[i].calculatePlantFitness(iGeneration, input);                        	//Calculate the fitness of the plant
	    vPopulationFitnesses[i] = vPlants[i].outputPlantFitness(); 		//Copy each plant fitness into the vector of fitnesses
	}
}

void PlantPopulation::mutate(int iTimestep, const Input input)
{
	// Function to mutate the plants of the population after reproduction
	// Takes:
	// - rng: 	The random number generator that is used everywhere in the program where random numbers are used
	// - input: Input read from user-provided input file
	// Puts out:
	// Nothing, but all plants in the population undergo random mutations.

	extern std::mt19937_64 rng;

	// Coinflip for gene duplication
	std::bernoulli_distribution duplicationCoinFlip(input.dDuplicationOdds);
	// Coinflip for gene Loss
	std::bernoulli_distribution lossCoinFlip(input.dLossOdds);
	// Coinflip for other mutations
	std::bernoulli_distribution biasedCoinFlip(input.dMutationOdds);

	int iGenomePairNr = vPlants[0].outputPlantGenome().outputGenePairs().size();

	for(int j = 0; j < input.iPopSize; ++j) //For each plant
	{
		for (int k = 0; k < iGenomePairNr * 2; k++) 	// For every gene
		{
			Gene geneMutate = vPlants[j].outputPlantGenome().outputGene(k/2, k%2); // Copy the gene to mutate for the following mutations
			if (geneMutate.proveExistence() == false){continue;} 	// You cannot mutate a non-existent gene

			if (duplicationCoinFlip(rng)) 						// Gene duplication
			{
				if (iGenomePairNr > input.iMaxNumLoci)
				{
					std::ofstream maxGenesHit;
					maxGenesHit.open("max_loci.csv", std::ios_base::app);
					maxGenesHit << input.iSeed << "," << iTimestep << std::endl;
					std::cout << "Gene not duplicated - maximum number of loci reached sim = " << input.iSeed << std::endl;
				} else
				{
					vPlants[j].duplicateGene(k/2, k%2, iNumGenes); 				//Duplicate the gene and add it to the end of the genome together with an empty gene
					for (int l = 0; l  < input.iPopSize; ++l)
					{
						if (l == j){continue;}
						else
						{
							vPlants[l].addEmptyGenepair(); //Add empty genepairs to all other plants
						}
					}
					++iGenomePairNr;
//					phylogenyFile << iTimestep << ", " << vPlants[j].outputPlantGenome().outputGene(k/2, k%2).cGeneID << ", ";
					geneMutate.vecHistory.push_back(geneMutate.cGeneID); //update the gene history
//					phylogenyFile << vPlants[j].outputPlantGenome().outputGene( vPlants[j].outputPlantGenome().outputGenePairs().size()-1, 0).cGeneID << ", Duplicated, " << 0 << std::endl;
					recordGene(input, iTimestep, vPlants[j].outputPlantGenome().outputGene( vPlants[j].outputPlantGenome().outputGenePairs().size()-1, 0));
					++iNumGenes;
				}
			}
			if (lossCoinFlip(rng)) 								// Gene loss
			{
//				phylogenyFile << iTimestep << ", " << vPlants[j].outputPlantGenome().outputGene(k/2, k%2).cGeneID << ", ";
//				phylogenyFile << 0 << ", lost , " << 0 << std::endl;
				geneMutate.loseGene();
//				std::vector<std::vector<Gene>> mutatedGenePairs = vPlants[j].outputPlantGenome().outputGenePairs();
//				mutatedGenePairs[k/2][k%2] = geneMutate;
//				vPlants[j].setGenomePairs(mutatedGenePairs);
				vPlants[j].setGene(geneMutate, k/2, k%2); //replace the old gene with the new one
				continue; //You cannot further mutate a non-existent gene
			}
			if (biasedCoinFlip(rng))							// Change dPromotor
			{
				boost::uniform_real<> dist(-input.dMaxConcentrationChange, input.dMaxConcentrationChange);
				double dChange = dist(rng);
				if (geneMutate.outputConcentration()+dChange > 0)
					{
					if (geneMutate.outputConcentration()+dChange < input.dMaxConcentration){geneMutate.setConcentration(geneMutate.outputConcentration()+dChange);}
					else {geneMutate.setConcentration(input.dMaxConcentration);};
					}
				else {geneMutate.setConcentration(0);};
//				phylogenyFile << iTimestep << ", " << geneMutate.cGeneID << ", ";
				geneMutate.vecHistory.push_back(geneMutate.cGeneID); //update the gene history
				geneMutate.cGeneID = "C" + std::to_string(iNumGenes);
//				phylogenyFile << geneMutate.cGeneID << ", Concentration, " << geneMutate.outputConcentration() << std::endl;
				recordGene(input, iTimestep, geneMutate);
				++iNumGenes;
			}

			if (biasedCoinFlip(rng))						// Change iEnzymeFunction
			{
				boost::random::uniform_int_distribution<> dist(0, input.iNumReactions - 1);
				geneMutate.setFunction(dist(rng));
//				phylogenyFile << iTimestep << ", " << geneMutate.cGeneID << ", ";
				geneMutate.vecHistory.push_back(geneMutate.cGeneID); //update the gene history
				geneMutate.cGeneID = "C" + std::to_string(iNumGenes);
//				phylogenyFile << geneMutate.cGeneID << ", EnzymeFunction, " << geneMutate.outputFunction() << std::endl;
				recordGene(input, iTimestep, geneMutate);
				++iNumGenes;
			}
			if (biasedCoinFlip(rng))						// Change values of VecK-C
			{
				boost::random::lognormal_distribution<double>  amount(input.dKMu, input.dKSigma);
				geneMutate.vecHistory.push_back(geneMutate.cGeneID); //update the gene history
				for (int metabolite = 0; metabolite < std::pow(2, input.iNumReactions); ++metabolite)
				{
					double dChange = amount(rng);
					while (dChange/geneMutate.outputMichaelis()[metabolite] > pow(10, 9)) //stay the same if kCat/kM would become higher than thermodynamically possible
					{
						dChange = amount(rng);
					}
					geneMutate.setKCatVal(metabolite, dChange);
//					phylogenyFile << iTimestep << ", " << geneMutate.cGeneID << ", ";
					geneMutate.cGeneID = "C" + std::to_string(iNumGenes);
//					phylogenyFile << geneMutate.cGeneID << ", KCat in position " << metabolite << " to:, " << geneMutate.outputKCat()[metabolite] << std::endl;
					}
				recordGene(input, iTimestep, geneMutate);
				++iNumGenes;
			}
			if (biasedCoinFlip(rng))						// Change values of VecK-M
			{
				boost::random::lognormal_distribution<double> amount(input.dKMu, input.dKSigma);
				geneMutate.vecHistory.push_back(geneMutate.cGeneID); //update the gene history
				for (int metabolite = 0; metabolite < std::pow(2, input.iNumReactions); ++metabolite)
				{
					double dChange = amount(rng);
					while (geneMutate.outputKCat()[metabolite]/dChange > pow(10, 9)) //stay the same if kCat/kM would become higher than thermodynamically possible
					{
						dChange = amount(rng);
					}
					geneMutate.setKMichaelisVal(metabolite, dChange);
//					phylogenyFile << iTimestep << ", " << geneMutate.cGeneID << ", ";
					geneMutate.cGeneID = "C" + std::to_string(iNumGenes);
//					phylogenyFile << geneMutate.cGeneID << ", KM in position " << metabolite << " to:, " << geneMutate.outputMichaelis()[metabolite] << std::endl;
				}
				recordGene(input, iTimestep, geneMutate);
				++iNumGenes;
			}
			vPlants[j].setGene(geneMutate, k/2, k%2); //replace the old gene with the new one
		}
		vPlants[j].listGenes(); //add all genes to the gene list
	}
}

void PlantPopulation::reproduce(const Input input)
{
	// Function for reproduction of the population based on fitness of the plants
	// Takes:
	// - rng: 	The random number generator that is used everywhere in the program where random numbers are used
	// - input: Input read from user-provided input file
	// Puts out:
	// - Nothing, but the population reproduces.

	extern std::mt19937_64 rng;

	std::vector<double> vLotteryNumbers;

	// If fitness has effect make the lottery dependent on fitness, otherwise not
	if(input.fitnessHasEffect == true){
		vLotteryNumbers = vPopulationFitnesses;
	} else {
		std::vector<double> vNoFitness(input.iPopSize, 1.0);
		vLotteryNumbers = vNoFitness;
	}

	// Load probability weights (fitnesses) into a std::discrete_distribution to create a lottery to pull the next generation from
	std::discrete_distribution<int> reproductionLottery(vLotteryNumbers.begin(), vLotteryNumbers.end());

	// Create a fair coinflipper to determine which allele of each genepair is sampled
	std::uniform_int_distribution<int> coinflip(0,1);

	std::vector<Plant> vecNewPopulation;
	vecNewPopulation.reserve(input.iPopSize); 		// Create an empty population vector to recieve the offspring

	Plant              pPlant(false, input); 	// Create an uninitialized plant to copy the genome into

	// Sample genotypes for next generation
	for(int j = 0; j < input.iPopSize; ++j)         // For the size of the population
	{
		int iPlant1 = reproductionLottery(rng);		// Pick a random plant
		std::vector<std::vector<Gene>> vecGenomeParentOne = vPlants[iPlant1].outputPlantGenome().outputGenePairs();
		int iPlant2 = reproductionLottery(rng);		// Pick another random plant
		std::vector<std::vector<Gene>> vecGenomeParentTwo = vPlants[iPlant2].outputPlantGenome().outputGenePairs();

		std::vector<std::vector<Gene>> vecGenomeOffspring(vecGenomeParentOne.size()); 	// Reception vector for sampled genes
		for (int k = 0; (unsigned)k < vecGenomeParentOne.size(); ++k)					// For each Genepair, take one allele of each parent randomly
		{
			vecGenomeOffspring[k].push_back( vecGenomeParentOne[k][coinflip(rng)] );
			vecGenomeOffspring[k].push_back( vecGenomeParentTwo[k][coinflip(rng)] );
		}
		pPlant.setGenomePairs(vecGenomeOffspring); 	// Sets vecGenePairs but not yet geneNetwork - that should happen after mutation (in case regulatory regions change)
		vecNewPopulation.push_back(pPlant);			// Add the plant to the new population
	}
	vPlants = vecNewPopulation;                    	// Replace the old population by the new one
//	outputLog << "Reproduction finished" << std::endl;
}

void PlantPopulation::update(int iGeneration, const Input input)
{
	// Function to update all parts of the population after reproduction and mutation
	// Takes:
	// - iGeneration: 	Generation number of the population
	// - input: 		Input read from user-provided input file
	// Puts out:
	// - Nothing, but it prints the gene networks to a file and updates the gene network, proteome, ODE and fitness values after reproduction

	for (int j = 0; j < input.iPopSize; ++j) //For each plant in the population
	{
		//3.4.3 Create and run ODE
		vPlants[j].runPlantMetabolism(iGeneration, j, input);
	}
	//3.4.4 Calculate fitness
	calculatePopulationFitness(iGeneration, input);
	if (vPlants[0].outputPlantGenome().outputGenePairs().size() > (unsigned)(input.iMaxNumLoci / 3 * 2)
			|| vPlants[0].outputPlantGenome().outputGenePairs().size() > (unsigned) iNumLociLast * 2
			) {cleanup(input);} //Cleanup universally empty gene pairs whenever the number of loci > 2/3 of the maximum
}

void PlantPopulation::outputStateToFiles(std::string outputLocation, const Input input, int iGeneration)
{
	// Function to write information about the population state to output files
	// Takes:
	// - outputLocation: 	The output folder to write information to.
	// Puts out:
	// - Nothing, but it writes information about the population state to output files.

	//3.1.1 Write information about each plant to the output file
//	fitnessFile << iGeneration << ", ";
	genomeFile 	<< iGeneration << ", ";
	for (int i = 0; i < input.iPopSize; i++)
	{
		std::vector<Gene> geneList = vPlants[i].outputPlantGenome().outputGeneList();
//		fitnessFile << vPopulationFitnesses[i] << ", ";
		for (int j = 0; (unsigned)j < geneList.size(); ++j)
		{
			genomeFile << geneList[j].cGeneID << " ";
		}
		genomeFile << ", ";

		int iRichness = 0;
		double dTotalConcentration = 0;
		double dBeneficialConcentration = 0;
		for (int j = 1; (unsigned)j < std::pow(2, input.iNumReactions); ++j)
		{
			double dFocalConc = vPlants[i].outputMetaboliteConcentrations()[j];
			if (dFocalConc > 0)
			{
				++iRichness;
				dTotalConcentration += dFocalConc;
				if (input.vecMetaboliteFitnessEffects[iGeneration][j] > 0)
				{
					dBeneficialConcentration += dFocalConc;
				}
			}
		}
		richnessFile << iGeneration << ", " << i << ", " << iRichness << std::endl;
		ratioFile    << iGeneration << ", " << i << ", " << dBeneficialConcentration/dTotalConcentration << std::endl;
		if(input.outputMetabolites == true){
			metaboliteFile << iGeneration << ", " << i << ", ";
			for (int j = 0; (unsigned)j < std::pow(2, input.iNumReactions); ++j)
			{
				metaboliteFile << vPlants[i].outputMetaboliteConcentrations()[j] << ",";
			}
			metaboliteFile 	<< "\n";
		}
	}
	genomeFile 		<< "\n";
//	fitnessFile 	<< "\n";
}

void PlantPopulation::recordGene(const Input input, int iGeneration, Gene gGene)
{
	geneFile << iGeneration << ", " << gGene.cGeneID << ", ";
	for (int i = 0; (unsigned)i < gGene.vecHistory.size(); ++i)
	{
		geneFile << gGene.vecHistory[i] << " ";
	}

	geneFile << ", " << gGene.outputFunction() << ", " << gGene.outputConcentration() << ", ";
	for (int i = 0; (unsigned)i < std::pow(2, input.iNumReactions); ++i)
	{
		geneFile << gGene.outputKCat()[i] << ", ";
	}
	for (int i = 0; (unsigned)i < std::pow(2, input.iNumReactions); ++i)
	{
		geneFile << gGene.outputMichaelis()[i] << ", ";
	}
	geneFile << std::endl;
}

void PlantPopulation::cleanup(Input input)
{
    // Function to remove loci that don't have any genes in them from the genome
	// Takes:
	// - nothing
	// Returns:
	// - nothing, but the genomes of all plants are rid of loci where none have any genes, making the object smaller.

	bool removeLocus;

	std::vector<std::vector<Gene>> plantGenome = vPlants[0].outputPlantGenome().outputGenePairs();

	for (int i = 0; (unsigned)i < plantGenome.size(); ++i) //for each gene in the first plant
	{
		if (plantGenome[i][0].proveExistence() == false && plantGenome[i][1].proveExistence() == false) //check whether the locus is empty
		{
			removeLocus = true;
			for (int j = 1; j < input.iPopSize; j++) //if it is, for each plant
			{
				if (vPlants[j].outputPlantGenome().outputGenePairs()[i][0].proveExistence() == true || vPlants[j].outputPlantGenome().outputGenePairs()[i][1].proveExistence() == true) //check whether any plant has anything on that locus
				{
					removeLocus = false;
					break;
				}
			}
			if (removeLocus == true) //If it is empty in every single plant
			{
//				outputLog << "remove locus " << i << std::endl;
				for (int k = 0; k < input.iPopSize; k++) //for each plant
				{
				  vPlants[k].removeLocus(i);//remove the locus
				}
				plantGenome = vPlants[0].outputPlantGenome().outputGenePairs();
				--i;
			}
		}

	}
	iNumLociLast = plantGenome.size();
}
```
