## Supplementary material for "A metabolic model to investigate the evolution of chemodiversity": Simulation code and files: Population.h

```
/*
 * Population.h
 *      Author: Frans
 */
#include "Plant.h"

#include <fstream>

#ifndef POPULATION_H_
#define POPULATION_H_

class PlantPopulation{
public:
	// Function to initialize
						PlantPopulation(const Input input);

	// Functions to return member variables
	std::vector<Plant>  outputPopulation(){return vPlants;}
	std::vector<double> outputPopulationFitness(){return vPopulationFitnesses;}

	// Functions to change the plant population
	void 				reproduce(Input input); 						// Function to use the fitnesses of the population to create a new population
	void 				mutate(int iTimestep, const Input input);		// Function to mutate plant genomes after reproduction
	void 				update(int iGeneration, const Input input); 	// Function to update the plants of the population after reproduction and mutation
	void                cleanup(const Input input);										// Function to remove gene pairs that are empty in the whole population to limit the size the genome takes
	void 				calculatePopulationFitness(int iGeneration, const Input input);

	// Functions to output information to output files
	void 				outputStateToFiles(std::string outputLocation, const Input input, int iGeneration);
	void 				recordGene(const Input input, int iGeneration, Gene gGene);

private:
	std::vector<Plant> 	vPlants;				// Vector of plants that make up the population
	std::vector<double> vPopulationFitnesses; 	// Vector of fitnesses of the plants that make up the population
	std::ofstream 		metaboliteFile; 		// File where, for each generation and each plant, the concentrations of each metabolite are stored.
	std::ofstream 		richnessFile; 			// File where, for each generation and each plant, the number of derived metabolites produced is stored.
	std::ofstream 		ratioFile;    	 		// File where, for each generation and each plant, the ratio of beneficial metabolites to total derived metabolites is stored.
	std::ofstream		genomeFile;				// File where, for each generation and each plant, geneID's of all genes in all plants are stored
	std::ofstream		fitnessFile;			// File where, for each generation and plant, their fitness is stored.
	std::ofstream		phylogenyFile;			// File where the descent of every gene is stored.
	std::ofstream		geneFile;				// File where the parameters of every unique gene are stored.
	int 				iNumGenes;				// Integer that tracks how many mutations have occurred in total
	int					iNumLociLast;			// Integer that tracks how many loci there were the last time loci were reduced
};


#endif /* POPULATION_H_ */
```
