## Supplementary material for "A metabolic model to investigate the evolution of chemodiversity": Simulation code and files: program.cpp

```
//============================================================================
// Name        : program.cpp
// Author      : Frans Thon
// Version     : Full model with all functionalities as of 2024
// Description : Main execution file
//============================================================================


#include <iostream>
#include <vector>
#include <chrono>
#include <fstream>
#include <random>

#include "Input.h"
#include "Population.h"

std::ofstream outputLog;
std::mt19937_64 rng;

int main(int argc, char* argv[]) {
	// Setup the program
	// Extract parameterfilename from program arguments or use default
	std::cout << "start program ";
	std::string inputFileName ("modelparameters.csv"); // For this to actually work, it's nescessary that the program is called from within the folder that contains modelparameters.txt

	if(argc > 1)
	{
		inputFileName = argv[1];
	}

	//1 Read input
	Input inputVals;
	inputVals.readInput(inputFileName);
	std::cout << inputVals.iFileID;
	std::cout << std::endl;
//	outputLog.open(inputVals.logFile, std::ios::app);

	//Set seed
	unsigned seed = inputVals.iSeed;
	rng.seed(seed);

    //Save starting time  of simulation
	std::ofstream timeFile;
	timeFile.open("time.csv", std::ios::app);
	auto startingTime = std::chrono::system_clock::to_time_t(std::chrono::system_clock::now());

	outputLog << "finished reading input" << std::endl;

	outputLog << "starting program " << argv[0] << " seed " << seed << std::endl; // Save also what the input parameter file is if there is one.

	// Initialize the plant population
	PlantPopulation ppPlantPopulation(inputVals); //2 Initialize population

	//3 Run the timeseries
	for (int i = 1; i < inputVals.iNumGenerations; ++i)
	{
		outputLog << std::endl << "Generation " << i << std::endl;
		//3.2 Create plants in new generation based on fitness
		ppPlantPopulation.reproduce(inputVals);
		outputLog << "Reproduction done " << std::endl;
		//3.3 Mutate plant genes
		ppPlantPopulation.mutate(i, inputVals);
		outputLog << "Mutations done " << std::endl;
		//3.4 Create and run metabolism ODE, calculate fitness of new generation
		ppPlantPopulation.update(i, inputVals);
		outputLog << "Updating done " << std::endl; //Generation nr for now
		//3.1 Write the state of the timeseries to output files
		if (i % inputVals.iOutputInterval == 0)
		{
//				std::cout << "Write state to file \n";
			ppPlantPopulation.outputStateToFiles(inputVals.outputFilesLocation, inputVals, i);
		}
		outputLog << "output done" << std::endl;
	}

	//Save ending time of simulation
	auto endingTime = std::chrono::system_clock::to_time_t(std::chrono::system_clock::now());
	timeFile << inputVals.iFileID << "," << startingTime << "," << endingTime << std::endl;
	return 0;
}
```
