## Supplementary material for "A metabolic model to investigate the evolution of chemodiversity": Simulation code and files: chemotyping.R

```
chemotyping <- function(metabolitedata, nrPlants, reactions) {
  #function to determine which metabolites are made by which plants in the simulation
  types <- vector("list", (2^reactions-1)) #which metabolites exist?
  for (i in 1:length(types)){
    types[[i]] <- c(1, 2^(which(intToBits(i) == 1)-1)+ 1) #metabolites with one combination present in a chemotype
  }
  temp  <- metabolitedata[,3:(3+length(types))] #subset dataframe
  temp2 <- temp > 0 #presence of metabolites dataframe
  frame <- as.data.frame(matrix(nrow = nrow(metabolitedata)/nrPlants, ncol = length(types) + 2)) #set up dataframe size receiving dataframe
  names(frame) <- c("t", '0', paste(types)) #colnames
  frame$t <- unique(metabolitedata$X.timestep.) #time
  for (gen in 1:nrow(frame)){ #per generation
    t <- 0
    for(type in rev(types)){ #per type (from most complex to least complex)
      temp3 <- temp2[((gen-1) * nrPlants + 1):(gen * nrPlants),type] #subset current generation for type
      temp4 <- as.data.frame(temp2[((gen-1) * nrPlants + 1):(gen * nrPlants),setdiff(types[[length(types)]], type)]) #subset current generation for *not* type
      val <- sum(rowSums(temp3) == length(type) & rowSums(temp4) == 0) #sum nr of plants where type is present but other metabolites are not
      frame[gen, ncol(frame) - t] <- val #add value to output dataframe
      t <- t + 1
    }
    frame[gen, 2] <- nrPlants - sum(frame[gen, 3:ncol(frame)]) #unmodified metabolite
  }
  return(frame)
}
```
