## Supplementary material for "A metabolic model to investigate the evolution of chemodiversity": Simulation code and files: kcat_km.R

```
kcat_km <- function(genome, genes, reactions, tables){
  #lower is more specific!
  K <- list()
  K$meanmedian <- vector(length = nrow(genome))
  K$se <- vector(length = nrow(genome))
  for (l in 1:nrow(genome)){
    table1 <- tables[[l]]
    if(nrow(table1) == 1) { #if no genes
      K$meanmedian[l:length(K$meanmedian)]  <- NA
      K$se[l:length(K$se)]  <- NA
      print( "error: no genes")
      return(K)
    }
    table1 <- table1[2:nrow(table1),]
    table1 <- table1[order(nchar(as.character(table1$genelist)), table1$genelist),] #order of age
    genelist2 <- sort(as.numeric(sub('.', '', table1$genelist)))
    genetable <- as.data.frame(matrix(nrow = length(genelist2), ncol = ncol(genes)))
    genetable <- genes[genelist2 + 1,]
    
    kcat_km    <- genetable[6:(5+2^reactions)]/genetable[(6+2^reactions):(5+2*2^reactions)] #kcat/km: specificity constant
    kcat_km    <- kcat_km[2:length(kcat_km)-1]
    maxes      <- as.vector(apply(kcat_km,1,max)) #maximum kcat/km
    mean_max_k <- mean(maxes)
    
    normalized <- kcat_km / maxes #normalized kcat/km: lower values for most entries if one kcat/km is much higher than others
    medians    <- apply(normalized, 1, median, na.rm=TRUE) #...which makes the normalized median an indicator of promiscuity
    counts     <- table1[,2]
    
    K$meanmedian[l] <- sum(medians*counts)/sum(counts)
    sd <- sqrt((sum((medians-K$meanmedian[l])^2*counts))/sum(counts))
    K$se[l] <- sd/sqrt(sum(counts))
  }
  return(K)
}
```
