## Supplementary material for "A metabolic model to investigate the evolution of chemodiversity": Simulation code and files: lttstats.R

```
lttstats <- function(genome, tables){
  lin <- list()
  lin$ltt$`1`<- lin$ltt$`2` <- lin$ltt$`3` <- lin$ltt$`4` <- lin$ltt$`5` <- vector(length = nrow(genome))
  lin$rltt   <- lin$ltt
  lin$mnatt  <- vector(length = nrow(genome))
  lin$tot    <- vector(length = nrow(genome))
  for (j in 1:nrow(genome)){
    table <- tables[[j]]
    table <- table[2:nrow(table),] #remove what isn't genes
    
    #lineages through time
    lin$ltt$`1`[j] <- sum(table[,2] > 0)
    lin$ltt$`2`[j] <- sum(table[,2] > 1)
    lin$ltt$`3`[j] <- sum(table[,2] > 2)
    lin$ltt$`4`[j] <- sum(table[,2] > 3)
    lin$ltt$`5`[j] <- sum(table[,2] > 4)
    
    #relative lineages through time: ltt/total lineages
    tot   <- sum(table$Freq)
    lin$tot[j]      <- tot
    
    lin$rltt$`1`[j] <- sum(table[,2] > 0)/tot
    lin$rltt$`2`[j] <- sum(table[,2] > 1)/tot
    lin$rltt$`3`[j] <- sum(table[,2] > 2)/tot
    lin$rltt$`4`[j] <- sum(table[,2] > 3)/tot
    lin$rltt$`5`[j] <- sum(table[,2] > 4)/tot
    
    #mean normalized abundance through time: mean of (abundance of each lineage / most common gene)
    lin$mnatt[j] <- mean(table[,2]/ max(table[,2]))
    
  }
  return(lin)
}
```
