## Supplementary material for "A metabolic model to investigate the evolution of chemodiversity": Simulation code and files: make_tables.R

```
make_tables <- function(genome){
  #tabulate gene frequencies by generation
  tables <- list()
  for (i in 1:nrow(genome)){
    genelist <- c()  
    for (k in 2:ncol(genome)){
      genelist <- c(genelist, str_split(genome[i,k], " ")[[1]]) 
    }
    tables[[i]] <- as.data.frame(table(genelist))
  }
  return(tables)
}
```
