## Supplementary material for "A metabolic model to investigate the evolution of chemodiversity": Simulation code and files: morris_trajectories.R

```
pick_trajectories <- function(parameters, levels, nr_trajectories, samplesize){
  #generate trajectories
  startpoints <- as.data.frame(matrix(nrow = samplesize, ncol = length(parameters))) #1000 trajectory starting points
  changeorder <- as.data.frame(matrix(nrow = samplesize, ncol = length(parameters)))
  to          <- as.data.frame(matrix(nrow = samplesize, ncol = length(parameters)))
  levelvec    <- 1:levels
  
  all_timepoints <- list()
  
  for (i in 1:samplesize){
    startpoints[i,] <- sample(levelvec, length(parameters), replace = TRUE) #sample random values
    changeorder[i,] <- sample(1:length(parameters)) #pick a random order for the parameters to be changed
    for (j in 1:length(parameters)){
      to[i,j] <- sample(levelvec[! levelvec %in% startpoints[i,j]], 1) #pick a second set of random values for the parameters to be change to
    }
  }
  
  all_timepoints[[1]] <- startpoints
  
  for (i in 1:length(parameters)){
    all_timepoints[[i+1]] <- startpoints
    all_timepoints[[i+1]][changeorder < (i + 1)] <- to[changeorder < i + 1]
  }
  trajectories <- all_timepoints
  return(trajectories)
}
```
