## Supplementary material for "A metabolic model to investigate the evolution of chemodiversity": Simulation code and files: posEffectRatio.R

```
posEffectRatio     <- function(metabolitedata, fitTable, nrPlants){ 
  #Calculates the the mean ratio of the concentration of 
  #beneficial metabolite and the total metabolite produced over all plants in each generation

  out <- list()
  
  total <- as.data.frame(matrix(nrow = nrPlants, ncol = nrow(metabolitedata)/nrPlants)) #dataframe with each row a plant and each column a timestep: total amount of metabolite
  
  beneficial <- as.data.frame(matrix(nrow = nrPlants, ncol = nrow(metabolitedata)/nrPlants)) #dataframe with each row a plant and each column a timestep: amount of beneficial metabolite
  
  genswitches <- c(unique(fitTable$generation), max(metabolitedata$X.timestep.) + 1)
  currentgen  <- 1
  
  for(i in 1:(nrow(metabolitedata)/nrPlants)){ #Per recording instance
    if (metabolitedata[((i-1)*nrPlants + 1),1] == genswitches[currentgen]){
      usefulmets <- fitTable[fitTable$phase == (currentgen - 1),]$fitnessval
      usefulmets <- usefulmets>0
      currentgen <- currentgen + 1
    }
    total[i]      <- rowSums(metabolitedata[((i-1)*nrPlants + 1):(i * nrPlants),4:ncol(metabolitedata)])
    if (sum(usefulmets) > 1){
      beneficial[i] <- rowSums(metabolitedata[((i-1)*nrPlants + 1):(i * nrPlants),which(usefulmets == TRUE) + 2]) 
    } else {beneficial[i] <- metabolitedata[((i-1)*nrPlants + 1):(i * nrPlants),which(usefulmets == TRUE) + 2]}
  }
  
  
  posMetRatio <- colMeans(beneficial/total, na.rm = TRUE) #vector with the mean ratio of beneficial to total metabolites: one row per generation, with the rows being different timesteps
  
  out$posMetRatio <- posMetRatio
  
  return(out)
}
```
