## Supplementary material for "A metabolic model to investigate the evolution of chemodiversity": Simulation code and files: richness.R

```
met_richness <-function(metabolitedata, nrPlants) {
  #Calculates average metabolite richness
  averages <- rowSums(metabolitedata[,4:ncol(metabolitedata)]>0, na.rm = TRUE) #richness per plant
  timeseries2 <- numeric(length = (nrow(metabolitedata)/nrPlants)) #prepare vector
  for (i in 0:(nrow(metabolitedata)/nrPlants - 1)){ #average for every generation
     timeseries2[i + 1] <- sum(averages[1:nrPlants+i*nrPlants])/nrPlants
  }
  return(timeseries2)
}
```
