## Supplementary material for "A metabolic model to investigate the evolution of chemodiversity": Simulation code and files: README.txt

This archive contains all files nescessary to recreate the model and all simulations used for the publication this is a supplement to.

Contents of the archive:

simulations
	batch 1
	batch 2
	robustness
R
program
	src

To recreate the model and simulations, first build the c++ model for which the source files are found in the 'program' directory.

Then place the built model, found in 'Debug', inside the 'simulations' folder.


#Main simulations
First adjust the 'CORES' parameter in the 'create_jobs_experiments.sh' files in the 'batch 1' and 'batch 2' directories for your machine. Then run these files to create the parameter files.

Start the simulations with 'start_run.txt' on a linux SLURM cluster.

Use the figures.R file in the 'simulations' directory to visualize the output of 'batch 1' and 'batch 2'.


#Robustness analysis
Unfortunately, we did not specify a random seed when we generated the random samples for the robustness analysis. For this reason, we have included a trajectories.RData file with the samples that we generated in the directory. If you want to create your own random samples for the robustness analysis, run the function in the make_files.R file in the 'robustness' directory inside the 'simulations' directory. Run the rest of that file to make the input files for the simulations.

To run the simulations for the the robustness analysis, set the number of cores in both the start_run.txt and start_simulations.R files, and then run the start_run.txt file on a linux SLURM cluster.

Use the variance_rank.R file to visualize the results of the robustness analysis.

If you cannot, or do not want to run all the simulations, the 'indices.RData' files contain the results of the analyses, which can be visualized with the figures.R and variance_rank.R files.
