## Supplementary material for "A metabolic model to investigate the evolution of chemodiversity": Simulation code and files: create_jobs_experiments.sh

```
#!/bin/bash
#Adjust for your machine
CORES=20

SCRIPTS_DIR="scripts_tmp"
OUTPUT_DIR="output"
LOG_DIR="logs"
BASHFILE="start_run.txt"
RMASTERFILE="start_simulations.R"
RINDICESFILE="extract_indices.R"
RANDOM_FITNESSES_FILE="random_fitness.csv"
MAX_LOCI_FILE="max_loci.csv"
TIME_FILE="time.csv"
REPLICATES=10

# Enter the variable parameters here:

#population parameters
i=1 #iseed
i2=${i}
j=0 #experiment/fitness seed
k=1 #zero seed first
l=1 #phase order seed first
m=1 #initial K seed 
iPopSizeVec=(1000)
iNumGenerationsVec=(1001)

#gene parameters
iNumGeneExps=1 #iterator for iNumGenePairs and iNumInitialCodingGenes
iNumGenePairs=(3)
identicalFunction=1
dBaseConcentrationVec=(0.001)

#mutations
dLossOddsVec=(0.01)
dDuplicationOddsVec=(0.0001)
iMaxNumLociVec=(80)
dMutationOddsVec=(0.0001)
dMaxConcentrationVec=(0.1)
dMaxConcentrationChangeVec=(0.001)
numK=1 #iterator for mu and sigma
dKMu=(-3.45)
dKSigma=(0.68)
iNumReactions=(3)

#metabolism
SetInitialKVec=(0)
iNumMetConcentrations=1 #iterator through dInitialMetConcentrationVec and Influx
dInitialMetConcentrationVec=(0.01)
dInfluxVec=(0.001)
dEffluxVec=(0.1)
dODELength=100
ODEstepperVec=("runge-kutta")
dSmallestODEStepSizeVec=(0.01)

#fitness
fitnessHasEffect=(1)
phaseTransitionAtVec=(250)
dFitnessCostsPerEnzyme=(1 10)
fitnessIsRandom=1
numExperiments=4 #number of different random fitness landscapes
dMinimumFitnessEffectVec=(0)
dMaximumFitnessEffectVec=(500)
numZeroEx=(10) #iterator: number of zero seeds (random patterns of fitness effects put to 0)
numZeroVec=(0 1 2 3 4 5 6)


outputODE=0
outputIntervals=(10)

#remove files that are about to be created
rm -rf ${SCRIPTS_DIR}
rm -rf ${OUTPUT_DIR}
rm -rf ${LOG_DIR}
rm -rf ${BASHFILE}
rm -rf ${RMASTERFILE}
rm -rf ${RINDICESFILE}
rm -rf ${RANDOM_FITNESSES_FILE}
rm -rf ${MAX_LOCI_FILE}
rm -rf ${TIME_FILE}

#create directories
mkdir ${SCRIPTS_DIR}
mkdir ${OUTPUT_DIR}
mkdir ${LOG_DIR}

#create Rscript that connects program to input files
echo "#!/usr/bin/Rscript --vanilla" >> ${RMASTERFILE}
echo "" >> ${RMASTERFILE}
echo "args <- commandArgs(TRUE)" >> ${RMASTERFILE}
echo "library(foreach)" >> ${RMASTERFILE}
echo "library(doMC)" >> ${RMASTERFILE}
echo "detectCores()" >> ${RMASTERFILE}
echo "registerDoMC(${CORES})" >> ${RMASTERFILE}
echo "" >> ${RMASTERFILE}
echo "files <- list.files(\"${SCRIPTS_DIR}/\")" >> ${RMASTERFILE}
echo "" >> ${RMASTERFILE}
echo "randorder <- sample(${i}:(${i}-1+length(files)))" >> ${RMASTERFILE}
echo "" >> ${RMASTERFILE}
echo "foreach (i=randorder)%dopar%{" >> ${RMASTERFILE}
echo "  file <- files[which(grepl(paste0(\"file_\", i, \"_\"), files, fixed=TRUE) == TRUE)]" >> ${RMASTERFILE}
echo "  if (file.exists(paste0(\"output/\", file)) == FALSE){" >> ${RMASTERFILE}
echo "    system(paste0('echo \"starting file \"', i))" >> ${RMASTERFILE}
echo "    system(paste0(\"./../program ${SCRIPTS_DIR}/\", file))" >> ${RMASTERFILE}
echo "  }" >> ${RMASTERFILE}
echo "}" >> ${RMASTERFILE}

#create bash file to start on cluster
echo "#!/bin/bash" >> ${BASHFILE}
echo "" >> ${BASHFILE}
echo "#SBATCH -J $(pwd | rev | awk -F / '{print $1}' | rev)" >> ${BASHFILE}
echo "#SBATCH -N 1" >> ${BASHFILE}
echo "#SBATCH --ntasks-per-node=${CORES}" >> ${BASHFILE}
echo "#SBATCH --ntasks-per-core=1" >> ${BASHFILE}
echo "" >> ${BASHFILE}
echo "Rscript ${RMASTERFILE}" >> ${BASHFILE}
echo "Rscript ${RINDICESFILE}" >> ${BASHFILE}

#create global output files
echo "simnum, fseed, phase, zero, generation, metabolite, fitnessval" >> ${RANDOM_FITNESSES_FILE}
echo "simnum, generation" >> ${MAX_LOCI_FILE}
echo "simnum, start, end" >> ${TIME_FILE}

                                                                                                  for dMinimumFitnessEffect in ${dMinimumFitnessEffectVec[*]}
                                                                                                  do
                                                                                               for dMaximumFitnessEffect in ${dMaximumFitnessEffectVec[*]}
                                                                                               do
												for ((geneExp=0; geneExp <(${iNumGeneExps}); ++geneExp))
												do
											     for SetInitialK in ${SetInitialKVec[*]}
											     do
                                                                                       for ((Kval=0; Kval<${numK}; ++Kval))
                                                                                       do
										       for fiteffect in ${fitnessHasEffect[*]}
										       do
									         for ((zero=k; zero <(${numZeroEx} + k); ++zero))
		 		                                               do 
                                                                           for ((position=j; position<(${numExperiments} + j); ++position))
                                                                           do                 
                                                                  for dSmallestODEStepSize in ${dSmallestODEStepSizeVec[*]}
                                                                  do
                                                               for ODEstepper in ${ODEstepperVec[*]}
                                                               do
                                                            for iMaxNumloci in ${iMaxNumLociVec[*]}
                                                            do
                                                      for dFitnessCostPerEnzyme in ${dFitnessCostsPerEnzyme[*]}
                                                      do
                                                   for outputInterval in ${outputIntervals[*]}
                                                   do
                                          for numZero in ${numZeroVec[*]}
                                          do
                                       for dEfflux in ${dEffluxVec[*]}
                                       do
                                     for phaseTransitionAt in ${phaseTransitionAtVec[*]}
                                     do
                                 for ((conc=0; conc<(${iNumMetConcentrations}); ++conc))
                                 do
                              for dDuplicationOdds in ${dDuplicationOddsVec[*]}
                              do
                           for dLossOdds in ${dLossOddsVec[*]}
                           do
                  for dMaxConcentrationChange in ${dMaxConcentrationChangeVec[*]}
                  do
               for dMaxConcentration in ${dMaxConcentrationVec[*]}
               do
            for dMutationOdds in ${dMutationOddsVec[*]}
            do
         for dBaseConcentration in ${dBaseConcentrationVec[*]}
         do
      for iNumGenerations in ${iNumGenerationsVec[*]}
      do
   for iPopSize in ${iPopSizeVec[*]}
   do
for ((replicate=1; replicate<=${REPLICATES}; ++replicate))
do
   filename=file_${i}_eCost_${dFitnessCostPerEnzyme}_numZero_${numZero}_zeroEx_${zero}_Ex_${position}_rep_${replicate}
   echoFile=${SCRIPTS_DIR}/${filename}
   echo "logFile ${LOG_DIR}/log_${i}.txt" >> ${echoFile}
   echo "" >> ${echoFile}
 #population parameters
   echo "iSeed ${i}">> ${echoFile}
   echo "initialKSeed ${m}" >> ${echoFile}   
   echo "fitnessSeed ${position}" >> ${echoFile}
   echo "zeroSeed ${zero}" >> ${echoFile}
   echo "iPopSize ${iPopSize}" >> ${echoFile}
   echo "iNumGenerations ${iNumGenerations}" >> ${echoFile}
   echo "" >> ${echoFile}
         
 #gene parameters 
   echo "iNumGenePairs ${iNumGenePairs[${geneExp}]}" >> ${echoFile} 
   echo "identicalFunction ${identicalFunction}" >> ${echoFile}
   echo "dBaseConcentration ${dBaseConcentration}" >> ${echoFile}   
   echo "" >> ${echoFile}

#mutations
   echo "dLossOdds ${dLossOdds}" >> ${echoFile}
   echo "dDuplicationOdds ${dDuplicationOdds}" >> ${echoFile}
   echo "iMaxNumLoci ${iMaxNumloci}" >> ${echoFile}
   echo "dMutationOdds ${dMutationOdds}" >> ${echoFile}   
   echo "dMaxConcentration ${dMaxConcentration}" >> ${echoFile}
   echo "dMaxConcentrationChange ${dMaxConcentrationChange}" >> ${echoFile}
   echo "dKMu ${dKMu[${Kval}]}" >> ${echoFile}
   echo "dKSigma ${dKSigma[${Kval}]}" >> ${echoFile}
   echo "iNumReactions ${iNumReactions}" >> ${echoFile}   
   echo "" >> ${echoFile}
   
#metabolism
   echo "setInitialK ${SetInitialK}" >> ${echoFile}
   echo "dInitialMetConcentration ${dInitialMetConcentrationVec[${conc}]}" >> ${echoFile}   
   echo "dInflux ${dInfluxVec[${conc}]}" >> ${echoFile}
   echo "dEfflux ${dEfflux}" >> ${echoFile}
   echo "dODELength ${dODELength}" >> ${echoFile}       
   echo "ODEstepper ${ODEstepper}" >> ${echoFile}
   echo "dSmallestODEStepSize ${dSmallestODEStepSize}" >> ${echoFile}
   echo "" >> ${echoFile}
   
#fitness
   echo "fitnessHasEffect ${fiteffect}" >> ${echoFile}
   declare -i iNumPhases2=${iNumGenerations}%${phaseTransitionAt}
   if [ ${iNumPhases2} = 0 ]; then
   	declare -i iNumPhases=${iNumGenerations}/${phaseTransitionAt}
   	else
   	declare -i iNumPhases=${iNumGenerations}/${phaseTransitionAt}+1
   fi 
   echo "iNumPhases ${iNumPhases}" >> ${echoFile}
   for ((phase=1; phase <(${iNumPhases}); ++phase))
   do
     declare -i time=${phaseTransitionAt}*${phase}
     echo "vecPhaseTransitionTimes${phase} ${time}" >> ${echoFile}
   done
   echo "dFitnessCostPerEnzyme ${dFitnessCostPerEnzyme}" >> ${echoFile}
   echo "fitnessIsRandom ${fitnessIsRandom}" >> ${echoFile}   
   echo "dMinimumFitnessEffect ${dMinimumFitnessEffect}" >> ${echoFile}   
   echo "dMaximumFitnessEffect ${dMaximumFitnessEffect}" >> ${echoFile}         
   echo "iNumZeroFitness ${numZero}" >> ${echoFile}
   echo "" >> ${echoFile}

#output   
   echo "outputODE ${outputODE}" >> ${echoFile}
   echo "outputFilesLocation ${OUTPUT_DIR}/${filename}" >> ${echoFile}
   echo "iOutputInterval ${outputInterval}" >> ${echoFile} 
   echo "file ${i}" >> ${echoFile}
  
   ((i=${i}+1))
done 
   done
      done
         done
            done
               done
                  done
                     done
                        done
                           done
                              done
                                 done
                                    done
                                       done
                                          done
                                             done
                                                done
                                                   done
                                                      done
                                                         done
                                                            done
                                                               done
                                                                  done
                                                                     done
                                                                        done
                                                                           done
                                                                                    
echo "#!/usr/bin/Rscript --vanilla" >> ${RINDICESFILE}
echo "" >> ${RINDICESFILE}
echo "args <- commandArgs(TRUE)" >> ${RINDICESFILE}
echo "library(foreach)" >> ${RINDICESFILE}
echo "library(doMC)" >> ${RINDICESFILE}
echo "detectCores()" >> ${RINDICESFILE}
echo "registerDoMC(${CORES})" >> ${RINDICESFILE}
echo "setwd(paste0(\"../../R\"))" >> ${RINDICESFILE}
echo "" >> ${RINDICESFILE}
echo "source(\"richness.R\")" >> ${RINDICESFILE}
echo "source(\"chemotyping.R\")" >> ${RINDICESFILE}
echo "source(\"kcat_km.R\")" >> ${RINDICESFILE}
echo "source(\"lttstats.R\")" >> ${RINDICESFILE}
echo "source(\"make_tables.R\")" >> ${RINDICESFILE}
echo "source(\"posEffectRatio.R\")" >> ${RINDICESFILE}
echo "" >> ${RINDICESFILE}
echo "require(stringr)" >> ${RINDICESFILE}
echo "require(reshape2)" >> ${RINDICESFILE}
echo "" >> ${RINDICESFILE}
echo "sessionInfo()" >> ${RINDICESFILE}	
echo "" >> ${RINDICESFILE}
echo "setwd(paste0(\"../simulations/$(pwd | rev | awk -F / '{print $1}' | rev)\"))" >> ${RINDICESFILE}
echo "" >> ${RINDICESFILE}
echo "reactions     <- ${iNumReactions}" >> ${RINDICESFILE}
echo "simlength     <- ${iNumGenerations}" >> ${RINDICESFILE}
echo "recordingfreq <- ${outputIntervals}" >> ${RINDICESFILE}
echo "popsize       <- ${iPopSize}" >> ${RINDICESFILE}
echo "" >> ${RINDICESFILE}
echo "files <- list.files(path = \"output/\")" >> ${RINDICESFILE}
echo "files2 <- vector(length = length(files))" >> ${RINDICESFILE}
echo "" >> ${RINDICESFILE}
echo "randFitness <- read.csv(paste0(\"random_fitness.csv\"), sep=",", header=TRUE)" >> ${RINDICESFILE}
echo "maxhit      <- read.csv(paste0(\"max_loci.csv\"), sep=",", header=TRUE)" >> ${RINDICESFILE}
echo "" >> ${RINDICESFILE}
echo "numSims              <- length(files)" >> ${RINDICESFILE}
echo "" >> ${RINDICESFILE}
echo "diversities          <- list()" >> ${RINDICESFILE}
echo "diversities\$richness <- diversities\$simpson <- diversities\$whittaker <- as.data.frame(matrix(nrow = numSims, ncol = simlength/recordingfreq + 1))" >> ${RINDICESFILE}
echo "K                    <- list()" >> ${RINDICESFILE}
echo "K\$meanmedian         <- K\$se <- as.data.frame(matrix(nrow = numSims, ncol = simlength/recordingfreq + 1))" >> ${RINDICESFILE}
echo "fitnessmeans         <- matrix(nrow = numSims, ncol = simlength/recordingfreq+1)" >> ${RINDICESFILE}
echo "chemotypes           <- list(length(numSims))" >> ${RINDICESFILE}
echo "lins                 <- list()" >> ${RINDICESFILE}
echo "lins\$ltt             <- list()" >> ${RINDICESFILE}
echo "lins\$rltt            <- list()" >> ${RINDICESFILE}
echo "lins\$ltt$\`1\` <- lins\$ltt$\`2\` <- lins\$ltt$\`3\` <- lins\$ltt$\`4\` <- lins\$ltt$\`5\` <- lins\$mnatt <- lins\$tot <- as.data.frame(matrix(nrow = numSims, ncol = simlength/recordingfreq + 1))" >> ${RINDICESFILE}
echo "lins\$rltt <- lins\$ltt" >> ${RINDICESFILE}
echo "ratio                <- list()" >> ${RINDICESFILE}
echo "ratio\$posMetRatio    <- ratio\$richratio <- as.data.frame(matrix(nrow = numSims, ncol = simlength/recordingfreq + 1))" >> ${RINDICESFILE}
echo "" >> ${RINDICESFILE}
echo "comb <- function(x, ...) {" >> ${RINDICESFILE}
echo "  lapply(seq_along(x)," >> ${RINDICESFILE}
echo "         function(i) c(x[[i]], lapply(list(...), function(y) y[[i]])))" >> ${RINDICESFILE}
echo "}" >> ${RINDICESFILE}
echo "" >> ${RINDICESFILE}
echo "rez <- foreach(i=${i2}:(${i2}-1 + length(files)), .combine='comb', .multicombine=TRUE," >> ${RINDICESFILE}
echo "               .init=list(list(), list(), list(), list(), list(), list(), list()," >> ${RINDICESFILE}
echo "                          list(), list(), list()))%dopar%{" >> ${RINDICESFILE}
echo "  print(i)" >> ${RINDICESFILE}
echo "  files2[i]                 <- files[which(grepl(paste0(\"file_\", i, \"_\"), files, fixed=TRUE) == TRUE)]" >> ${RINDICESFILE}
echo "" >> ${RINDICESFILE}
echo "  metabolitedata            <- read.csv(paste0(\"output/\", files2[i], \"/metabolite\"), sep=\",\", header=TRUE)" >> ${RINDICESFILE}
echo "  genome                    <- read.csv(paste0(\"output/\", files2[i], \"/genome\"), sep=\",\", header=TRUE)" >> ${RINDICESFILE}
echo "  genes                     <- read.csv(paste0(\"output/\", files2[i], \"/genes\"), sep=\",\", header=TRUE)" >> ${RINDICESFILE}
echo "  fitTable                  <- randFitness[randFitness\$simnum == i,]   " >> ${RINDICESFILE}
echo "  fitTable                  <- fitTable[!duplicated(fitTable), ]" >> ${RINDICESFILE}
echo "" >> ${RINDICESFILE}
echo "  chemotypes2               <- chemotyping(metabolitedata, popsize, reactions)" >> ${RINDICESFILE}
echo "" >> ${RINDICESFILE}
echo "  metabolitedata            <- metabolitedata[1:(length(metabolitedata)-1)]" >> ${RINDICESFILE}
echo "" >> ${RINDICESFILE}
echo "  richness     <- met_richness(metabolitedata, nrPlants = popsize)" >> ${RINDICESFILE}
echo "" >> ${RINDICESFILE}
echo "  tables       <- make_tables(genome)" >> ${RINDICESFILE}
echo "" >> ${RINDICESFILE}
echo "  posRatio     <- posEffectRatio(metabolitedata, fitTable, popsize)" >> ${RINDICESFILE}
echo "  posMetRatio  <- posRatio\$posMetRatio" >> ${RINDICESFILE}
echo "" >> ${RINDICESFILE}
echo "  Ks           <- kcat_km(genome, genes, reactions, tables) # mean median normalized Kcat/Km" >> ${RINDICESFILE}
echo "  meanmedian   <- Ks\$meanmedian" >> ${RINDICESFILE}
echo "" >> ${RINDICESFILE}
echo "  lin  <- lttstats(genome, tables)" >> ${RINDICESFILE}
echo "  ltt1 <- lin\$ltt$\`1\` #lineages through time" >> ${RINDICESFILE}
echo "  ltt2 <- lin\$ltt$\`2\`" >> ${RINDICESFILE}
echo "  ltt3 <- lin\$ltt$\`3\`" >> ${RINDICESFILE}
echo "  ltt4 <- lin\$ltt$\`4\`" >> ${RINDICESFILE}
echo "  ltt5 <- lin\$ltt$\`5\`" >> ${RINDICESFILE}
echo "" >> ${RINDICESFILE}
echo "  tot <- lin\$tot" >> ${RINDICESFILE}
echo "" >> ${RINDICESFILE}
echo "" >> ${RINDICESFILE}
echo "  list(chemotypes2, richness, meanmedian, ltt1, ltt2, ltt3, ltt4, ltt5, posMetRatio, tot)" >> ${RINDICESFILE}
echo "}" >> ${RINDICESFILE}
echo "" >> ${RINDICESFILE}
echo "for(i in 1:length(files)){" >> ${RINDICESFILE}
echo "  chemotypes[[i]]              <- rez[[1]][[i]]" >> ${RINDICESFILE}
echo "" >> ${RINDICESFILE}
echo "  diversities\$richness[i,]     <- rez[[2]][[i]]" >> ${RINDICESFILE}
echo "" >> ${RINDICESFILE}
echo "  K\$meanmedian[i,] <- rez[[3]][[i]]" >> ${RINDICESFILE}
echo "" >> ${RINDICESFILE}
echo "  lins\$ltt\$\`1\`[i,]  <- rez[[4]][[i]]" >> ${RINDICESFILE}
echo "  lins\$ltt\$\`2\`[i,]  <- rez[[5]][[i]]" >> ${RINDICESFILE}
echo "  lins\$ltt\$\`3\`[i,]  <- rez[[6]][[i]]" >> ${RINDICESFILE}
echo "  lins\$ltt\$\`4\`[i,]  <- rez[[7]][[i]]" >> ${RINDICESFILE}
echo "  lins\$ltt\$\`5\`[i,]  <- rez[[8]][[i]]" >> ${RINDICESFILE}
echo "" >> ${RINDICESFILE}
echo "  ratio\$posMetRatio[i,] <- rez[[9]][[i]]" >> ${RINDICESFILE}
echo "" >> ${RINDICESFILE}
echo "  lins\$tot[i,]    <- rez[[10]][[i]]" >> ${RINDICESFILE}
echo "}" >> ${RINDICESFILE}
echo "" >> ${RINDICESFILE}
echo "allhits <- as.data.frame(matrix(nrow = numSims, ncol = simlength, data = FALSE))" >> ${RINDICESFILE}
echo "" >> ${RINDICESFILE}
echo "for (i in 1:numSims){" >> ${RINDICESFILE}
echo "  if (any(maxhit\$simnum == ${i2} -1 +i)){" >> ${RINDICESFILE}
echo "    places <- unique(maxhit\$generation[which(maxhit\$simnum == ${i2} -1 +i)])" >> ${RINDICESFILE}
echo "    allhits[i,places] <- TRUE" >> ${RINDICESFILE}
echo "  }" >> ${RINDICESFILE}
echo "}" >> ${RINDICESFILE}
echo "" >> ${RINDICESFILE}
echo "save.image(file='indices.RData')" >> ${RINDICESFILE}

echo "done ..."
```
