## Supplementary material for "A metabolic model to investigate the evolution of chemodiversity": Simulation code and files: figures2.R

```
rm(list = ls())

require(reshape2)
require(ggplot2)
require(ggpubr)
require(patchwork)
require(forcats)
require(readr)
require(viridis)

names                  <- c("run", "set", "exp", "ecost", "zero", "zexp", "idFunc")
bigdescriptions        <- as.data.frame(matrix(nrow = 1200, ncol = length(names)))
names(bigdescriptions) <- names
cbbPalette <- c("#000000", "#56B4E9", "#F0E442", "#009E73", "#D55E00", "#CC79A7", "#E69F00", "#0072B2", "#999999", "#FFFFFF")

trial <- 1
repNum        <- 10
reactions     <- 10
recordingfreq <- 10
simlength     <- 1000
numSims       <- 60
popsize       <- 1000
experiments   <- c(0)
zexpvec       <- c(1:10)
enzymeCostVec <- c(1, 10)
numZeroVec    <- c(923, 1013, 1022)
idFunc        <- c(0)

seed <- 1
for (zex in zexpvec){
  for (exp in experiments){
    for (ecost in enzymeCostVec){
      for (nZero in numZeroVec){
        for (rep in 1:repNum){
          print(seed) 
          bigdescriptions[seed,] <- c(seed + 11200, trial, exp+1, ecost, nZero, zex, idFunc)
          seed <- seed + 1
        }
      }
    }
  }
}

trial <- 2

repNum        <- 10
reactions     <- 10
recordingfreq <- 10
simlength     <- 1000
numSims       <- 60
popsize       <- 1000
experiments   <- c(0)
zexpvec       <- c(1:10)
enzymeCostVec <- c(1, 10)
numZeroVec    <- c(923, 1013, 1022)
idFunc        <- c(1)

for (zex in zexpvec){
  for (exp in experiments){
    for (ecost in enzymeCostVec){
      for (nZero in numZeroVec){
        for (rep in 1:repNum){
          print(seed)
          bigdescriptions[seed,] <- c(seed + 11200, trial, exp+1, ecost, nZero, zex, idFunc)
          seed <- seed + 1
        }
      }
    }
  }
}

setwd(paste0("screening1"))

load("indices.RData")

Ktab            <- K$meanmedian
lttab           <- lins$ltt$`1`
totaltab        <- lins$tot
richness        <- diversities$richness
maxgeneshit     <- allhits
posMetRatio     <- ratio$posMetRatio

setwd(paste0("../screening2"))

load("indices.RData")

Ktab            <- rbind(Ktab, K$meanmedian)
lttab           <- rbind(lttab, lins$ltt$`1`)
totaltab        <- rbind(totaltab, lins$tot)
richness        <- rbind(richness, diversities$richness)
maxgeneshit     <- rbind(maxgeneshit, allhits)
posMetRatio     <- rbind(posMetRatio, ratio$posMetRatio)

relevants <- randFitness[randFitness$generation == 0 | randFitness$generation == 250 | randFitness$generation == 500 | randFitness$generation == 750, ]

bigdescriptions2 <- unique(bigdescriptions[3:7])
df               <- as.data.frame(matrix(nrow = nrow(bigdescriptions2), ncol = 101))

#richness
bigplotlist <- list()
bigplotlist2 <- list()

richtable          <- cbind(bigdescriptions, richness)
richtable[8:ncol(richtable)] <- richtable[8:ncol(richtable)] - 1 #exclude unmodified metabolite

richtable2         <- cbind(bigdescriptions2, df)

for(i in 1:nrow(bigdescriptions2)){
  richtable2[i,6:ncol(richtable2)] <- colMeans(richtable[richtable[4] == bigdescriptions2[i,2] & 
                       richtable[5] == bigdescriptions2[i,3] & 
                       richtable[6] == bigdescriptions2[i,4] &
                       richtable[7] == bigdescriptions2[i,5] ,8:ncol(richtable)], na.rm = TRUE) 
}

richplot           <- melt(richtable2, id.vars = names(bigdescriptions2))
richplot$variable  <- (as.numeric(richplot$variable)-1)*recordingfreq
names(richplot)    <- c(names(richplot)[1:length(names(bigdescriptions2))], "generations", "richness")
richplot$ecost     <- paste("enzyme cost ", richplot$ecost)
richplot$zero      <- 1023 - richplot$zero
richplot$idFunc[richplot$idFunc == 1] <- "identical"
richplot$idFunc[richplot$idFunc == 0] <- "different"


bigplotlist[[1]] <- ggplot(richplot, aes(x = generations, y = richness
                    , linetype = interaction(idFunc)
                    , colour = interaction(zero)
)) +
  labs(title = paste("E Richness summary"), linetype = "starting function", colour = "number of beneficial metabolites")+
  theme_classic()+
  facet_wrap(vars(ecost))+
  coord_cartesian(ylim=c(0, 10))+
  theme(strip.placement = "outside", legend.position="bottom")+
  geom_vline(xintercept = c(1:3*250), linetype='dashed', color='red', size=0.5)+
  scale_colour_manual(values = cbbPalette)+
  stat_summary( geom = "smooth", se = TRUE,
  )
# ggsave("richness_summary.pdf")

richplot2 <- richplot[(richplot$generations + 10) %% 250 == 0,]
richplot2$generations <- (richplot2$generations + 10)/250

bigplotlist2[[1]] <- ggplot(richplot2, aes(x = zero, y = richness
                                           , linetype = interaction(idFunc)
                                           , colour = as.factor(generations)
                                           , group = interaction(idFunc, generations))) +
  labs(title = paste("E Richness summary"), linetype = "starting function", colour = "phase", x = "number of beneficial metabolites")+
  theme_classic()+
  facet_wrap(vars(ecost))+
  theme(strip.placement = "outside", legend.position="bottom")+
  # geom_vline(xintercept = c(1:3*250), linetype='dashed', color='red', size=0.5)+
  scale_colour_manual(values = cbbPalette)+
  stat_summary( geom = "smooth", se = TRUE)

for (j in c(1, 10)){
  
  richtable2         <- richtable[richtable$ecost == j,]
  richplot           <- melt(richtable2, id.vars = names)
  richplot$variable  <- (as.numeric(richplot$variable)-1)*recordingfreq
  names(richplot)    <- c(names(richplot)[1:length(names(bigdescriptions))], "generations", "richness")
  richplot$zero      <- 1023 - richplot$zero
  richplot$zexp      <- paste("scenario", richplot$zexp)
  richplot$idFunc[richplot$idFunc == 1] <- "identical"
  richplot$idFunc[richplot$idFunc == 0] <- "different"
  
  ggplot(richplot, aes(x = generations, y = richness
                      , linetype = interaction(idFunc)
                      , colour = interaction(zero)
                      , group = run
  )) +
    labs(title = paste("Richness, Enzyme cost ", j), linetype = "starting function", colour = "number of beneficial metabolites")+
    theme_classic()+
    coord_cartesian(ylim=c(0, 25))+
    facet_wrap(~forcats::fct_relevel(zexp, paste("scenario", 1:10)))+
    theme(strip.placement = "outside", legend.position="right")+
    geom_vline(xintercept = c(1:3*250), linetype='dashed', color='red', size=0.5)+
    scale_colour_manual(values = cbbPalette)+
    stat_summary( geom = "smooth", se = TRUE,
    )
  ggsave(paste0("screening_richness_ecost_", j, ".pdf"), width = 22, height = 13, units = "cm")
}


#ltt
ltttable  <- cbind(bigdescriptions, lttab)
lttplots1         <- list()
lttplots2         <- list()

ltttable2       <- cbind(bigdescriptions2, df)

for(i in 1:nrow(bigdescriptions2)){
  ltttable2[i,6:ncol(ltttable2)] <- colMeans(ltttable[ltttable[4] == bigdescriptions2[i,2] & 
                                                           ltttable[5] == bigdescriptions2[i,3] & 
                                                           ltttable[6] == bigdescriptions2[i,4] &
                                                           ltttable[7] == bigdescriptions2[i,5] ,8:ncol(ltttable)], na.rm = TRUE) 
}

  lttplot           <- melt(ltttable2, id.vars = names(bigdescriptions2))
  lttplot$variable  <- (as.numeric(lttplot$variable)-1)*recordingfreq
  names(lttplot)    <- c(names(lttplot)[1:length(names(bigdescriptions2))], "generations", "ltt")
  lttplot$ecost     <- paste("enzyme cost ", lttplot$ecost)
  lttplot$zero      <- 1023 - lttplot$zero
  lttplot$idFunc[lttplot$idFunc == 1] <- "identical"
  lttplot$idFunc[lttplot$idFunc == 0] <- "different"
  
  
  bigplotlist[[2]] <- ggplot(lttplot, aes(x = generations, y = ltt
                                           , linetype = interaction(idFunc)
                                           , colour = interaction(zero)
  )) +
    labs(title = paste("B Sum of alleles summary"), linetype = "starting function", colour = "number of beneficial metabolites", y = "sum of alleles")+
    theme_classic()+
    facet_wrap(vars(ecost))+
    theme(strip.placement = "outside", legend.position="none")+
    geom_vline(xintercept = c(1:3*250), linetype='dashed', color='red', size=0.5)+
    scale_colour_manual(values = cbbPalette)+
    stat_summary( geom = "smooth", se = TRUE,
    )
  # ggsave("ltt_summary.jpg")
  
  lttplot2 <- lttplot[(lttplot$generations + 10) %% 250 == 0,]
  lttplot2$generations <- (lttplot2$generations + 10)/250
  
  bigplotlist2[[2]] <- ggplot(lttplot2, aes(x = zero, y = ltt
                                            , linetype = interaction(idFunc)
                                            , colour = as.factor(generations)
                                            , group = interaction(idFunc, generations))) +
    labs(title = paste("B Sum of alleles summary"), linetype = "starting function", colour = "phase", x = "number of beneficial metabolites", y = "sum of alleles")+
    theme_classic()+
    facet_wrap(vars(ecost))+
    theme(strip.placement = "outside", legend.position="none")+
    # geom_vline(xintercept = c(1:3*250), linetype='dashed', color='red', size=0.5)+
    scale_colour_manual(values = cbbPalette)+
    stat_summary( geom = "smooth", se = TRUE)

  for (j in c(1, 10)){

    ltttable2         <- ltttable[ltttable$ecost == j,]
    lttplot           <- melt(ltttable2, id.vars = names)
    lttplot$variable  <- (as.numeric(lttplot$variable)-1)*recordingfreq
    names(lttplot)    <- c(names(lttplot)[1:length(names(bigdescriptions))], "generations", "ltt")
    lttplot$zero      <- 1023 - lttplot$zero
    lttplot$zexp      <- paste("scenario", lttplot$zexp)
    lttplot$idFunc[lttplot$idFunc == 1] <- "identical"
    lttplot$idFunc[lttplot$idFunc == 0] <- "different"

    ggplot(lttplot, aes(x = generations, y = ltt
                                           , linetype = interaction(idFunc)
                                           , colour = interaction(zero)
                                           , group = run
    )) +
      labs(title = paste("Sum of alleles, Enzyme cost ", j), linetype = "starting function", colour = "number of beneficial metabolites", y = "sum of alleles")+
      theme_classic()+
      facet_wrap(~forcats::fct_relevel(zexp, paste("scenario", 1:10)))+
      theme(strip.placement = "outside", legend.position="right")+
      geom_vline(xintercept = c(1:3*250), linetype='dashed', color='red', size=0.5)+
      scale_colour_manual(values = cbbPalette)+
      stat_summary( geom = "smooth", se = TRUE,
      )
    ggsave(paste("screening_ltt_all_ecost", j, ".pdf"), width = 22, height = 13, units = "cm")
  }

  
  #total
  totaltable  <- cbind(bigdescriptions, totaltab)
  totalplots1         <- list()
  totalplots2         <- list()
  
  totaltable2       <- cbind(bigdescriptions2, df)
  
  for(i in 1:nrow(bigdescriptions2)){
    totaltable2[i,6:ncol(totaltable2)] <- colMeans(totaltable[totaltable[4] == bigdescriptions2[i,2] & 
                                                          totaltable[5] == bigdescriptions2[i,3] & 
                                                          totaltable[6] == bigdescriptions2[i,4] &
                                                          totaltable[7] == bigdescriptions2[i,5] ,8:ncol(totaltable)], na.rm = TRUE) 
  }

  totalplot           <- melt(totaltable2, id.vars = names(bigdescriptions2))
  totalplot$variable  <- (as.numeric(totalplot$variable)-1)*recordingfreq
  names(totalplot)    <- c(names(totalplot)[1:length(names(bigdescriptions2))], "generations", "total")
  totalplot$ecost     <- paste("enzyme cost ", totalplot$ecost)
  totalplot$zero      <- 1023 - totalplot$zero
  totalplot$idFunc[totalplot$idFunc == 1] <- "identical"
  totalplot$idFunc[totalplot$idFunc == 0] <- "different"
  
  
  bigplotlist[[3]] <- ggplot(totalplot, aes(x = generations, y = total
                      , linetype = interaction(idFunc)
                      , colour = interaction(zero)
  )) +
    labs(title = paste("C Number of gene copies summary"), linetype = "starting function", colour = "number of beneficial metabolites", y = "total gene copies")+
    theme_classic()+
    facet_wrap(vars(ecost))+
    theme(strip.placement = "outside", legend.position="none")+
    geom_vline(xintercept = c(1:3*250), linetype='dashed', color='red', size=0.5)+
    scale_colour_manual(values = cbbPalette)+
    stat_summary( geom = "smooth", se = TRUE,
    )
  # ggsave("total_summary.jpg")
  
  totalplot2 <- totalplot[(totalplot$generations + 10) %% 250 == 0,]
  totalplot2$generations <- (totalplot2$generations + 10)/250
  
  bigplotlist2[[3]] <- ggplot(totalplot2, aes(x = zero, y = total
                                              , linetype = interaction(idFunc)
                                              , colour = as.factor(generations)
                                              , group = interaction(idFunc, generations))) +
    labs(title = paste("C Number of gene copies summary"), linetype = "starting function", colour = "phase", x = "number of beneficial metabolites", y = "number of gene copies")+
    theme_classic()+
    facet_wrap(vars(ecost))+
    theme(strip.placement = "outside", legend.position="none")+
    # geom_vline(xintercept = c(1:3*250), linetype='dashed', color='red', size=0.5)+
    scale_colour_manual(values = cbbPalette)+
    stat_summary( geom = "smooth", se = TRUE)
  
  for (j in c(1, 10)){
    
    totaltable2         <- totaltable[totaltable$ecost == j,]
    totalplot           <- melt(totaltable2, id.vars = names)
    totalplot$variable  <- (as.numeric(totalplot$variable)-1)*recordingfreq
    names(totalplot)    <- c(names(totalplot)[1:length(names(bigdescriptions))], "generations", "total")
    totalplot$zero      <- 1023 - totalplot$zero
    totalplot$zexp      <- paste("scenario", totalplot$zexp)
    totalplot$idFunc[totalplot$idFunc == 1] <- "identical"
    totalplot$idFunc[totalplot$idFunc == 0] <- "different"
    
    ggplot(totalplot, aes(x = generations, y = total
                        , linetype = interaction(idFunc)
                        , colour = interaction(zero)
                        , group = run
    )) +
      labs(title = paste("Number of gene copies, Enzyme cost ", j), linetype = "starting function", colour = "number of beneficial metabolites", y = "total gene copies")+
      theme_classic()+
      facet_wrap(~forcats::fct_relevel(zexp, paste("scenario", 1:10)))+
      theme(strip.placement = "outside", legend.position="right")+
      geom_vline(xintercept = c(1:3*250), linetype='dashed', color='red', size=0.5)+
      scale_colour_manual(values = cbbPalette)+
      stat_summary( geom = "smooth", se = TRUE,
      )
    ggsave(paste("screening_total_all_ecost", j, ".pdf"), width = 22, height = 13, units = "cm")
  } 

#promiscuity
Ktable   <- cbind(bigdescriptions, Ktab)

Ktable2       <- cbind(bigdescriptions2, df)

for(i in 1:nrow(bigdescriptions2)){
  Ktable2[i,6:ncol(Ktable2)] <- colMeans(Ktable[Ktable[4] == bigdescriptions2[i,2] & 
                                                        Ktable[5] == bigdescriptions2[i,3] & 
                                                        Ktable[6] == bigdescriptions2[i,4] &
                                                        Ktable[7] == bigdescriptions2[i,5] ,8:ncol(Ktable)], na.rm = TRUE) 
}


    Kplot <- melt(Ktable2, id.vars = names(bigdescriptions2))

    Kplot$variable  <- (as.numeric(Kplot$variable)-1)*recordingfreq
    names(Kplot)    <- c(names(Kplot)[1:length(names(bigdescriptions2))], "generations", "promiscuity index")
    Kplot$zero  <- 1023 - Kplot$zero
    Kplot$ecost     <- paste("enzyme cost ", Kplot$ecost)
    Kplot$idFunc[Kplot$idFunc == 1] <- "identical"
    Kplot$idFunc[Kplot$idFunc == 0] <- "different"
    
    
    bigplotlist[[4]] <- ggplot(Kplot, aes(x = generations, y = `promiscuity index`
                      , linetype = interaction(idFunc)
                      , colour = interaction(zero)
    )) +
      labs(title = paste0("A Promiscuity index summary"), linetype = "starting function", colour = "number of beneficial metabolites")+
      theme_classic()+
      facet_wrap(vars(ecost))+
      theme(strip.placement = "outside", legend.position="none")+
      geom_vline(xintercept = c(1:3*250), linetype='dashed', color='red', size=0.5)+
      scale_colour_manual(values=cbbPalette)+
      stat_summary( geom = "smooth", se = TRUE, alpha = 0.1
      )
    
# ggsave(paste("Kplots_summary.pdf"))

    Kplot2 <- Kplot[(Kplot$generations + 10) %% 250 == 0,]
    Kplot2$generations <- (Kplot2$generations + 10)/250
    
    bigplotlist2[[4]] <- ggplot(Kplot2, aes(x = zero, y = `promiscuity index`
                                            , linetype = interaction(idFunc)
                                            , colour = as.factor(generations)
                                            , group = interaction(idFunc, generations))) +
      labs(title = paste("A Promiscuity index summary"), linetype = "starting function", colour = "phase", x = "number of beneficial metabolites")+
      theme_classic()+
      facet_wrap(vars(ecost))+
      theme(strip.placement = "outside", legend.position="none")+
      # geom_vline(xintercept = c(1:3*250), linetype='dashed', color='red', size=0.5)+
      scale_colour_manual(values = cbbPalette)+
      stat_summary( geom = "smooth", se = TRUE)    
    
for (j in c(1, 10)){
  
  Ktable22            <- Ktable[Ktable$ecost == j,]
  Kplot               <- melt(Ktable22, id.vars = names)
  totalplot           <- Kplot
  totalplot$variable  <- (as.numeric(totalplot$variable)-1)*recordingfreq
  names(totalplot)    <- c(names(totalplot)[1:length(names(bigdescriptions))], "generations", "promiscuity index")
  totalplot$zero      <- 1023 - totalplot$zero
  totalplot$zexp      <- as.factor(paste("scenario", totalplot$zexp))
  totalplot$idFunc[totalplot$idFunc == 1] <- "identical"
  totalplot$idFunc[totalplot$idFunc == 0] <- "different"
  
  ggplot(totalplot, aes(x = generations, y = `promiscuity index`
                        , linetype = interaction(idFunc)
                        , colour = interaction(zero)
                        , group = run
  )) +
    labs(title = paste("Promiscuity index, Enzyme cost ", j), linetype = "starting function", colour = "number of beneficial metabolites")+
    theme_classic()+
    facet_wrap(~forcats::fct_relevel(zexp, paste("scenario", 1:10)))+
    theme(strip.placement = "outside", legend.position="right")+
    geom_vline(xintercept = c(1:3*250), linetype='dashed', color='red', size=0.5)+
    scale_colour_manual(values = cbbPalette)+
    stat_summary( geom = "smooth", se = TRUE,
    )
  ggsave(paste("screening_Ktable_all_ecost", j, ".pdf"), width = 22, height = 13, units = "cm")
}  


#ratio
Rtable  <- cbind(bigdescriptions, posMetRatio)
Rtable2       <- cbind(bigdescriptions2, df)

for(i in 1:nrow(bigdescriptions2)){
  Rtable2[i,6:ncol(Rtable2)] <- colMeans(Rtable[Rtable[4] == bigdescriptions2[i,2] & 
                                                  Rtable[5] == bigdescriptions2[i,3] & 
                                                  Rtable[6] == bigdescriptions2[i,4] &
                                                  Rtable[7] == bigdescriptions2[i,5] ,8:ncol(Rtable)], na.rm = TRUE) 
}
  Rplot           <- melt(Rtable2, id.vars = names(bigdescriptions2))
  Rplot$variable  <- (as.numeric(Rplot$variable)-1)*recordingfreq
  names(Rplot)    <- c(names(Rplot)[1:length(names(bigdescriptions2))], "generations", "ratio")
  Rplot$ecost     <- paste("enzyme cost ", Rplot$ecost)
  Rplot$zero      <- 1023 - Rplot$zero
  Rplot$idFunc[Rplot$idFunc == 1] <- "identical"
  Rplot$idFunc[Rplot$idFunc == 0] <- "different"

  bigplotlist[[5]] <- ggplot(Rplot, aes(x = generations, y = ratio
                                       , linetype = interaction(idFunc)
                                       , colour = interaction(zero)
  )) +
    labs(title = paste("D Ratio between beneficial and total metabolite concentration summary"),
         linetype = "starting function",
         colour = "number of beneficial metabolites", 
         y = "beneficial ratio")+
    theme_classic()+
    facet_wrap(vars(ecost))+
    theme(strip.placement = "outside", legend.position="none")+
    geom_vline(xintercept = c(1:3*250), linetype='dashed', color='red', size=0.5)+
    scale_colour_manual(values=cbbPalette)+
    stat_summary(geom = "smooth", se = TRUE,
    )
  # ggsave(paste("Rplot_summary.pdf"))
  
  Rplot2 <- Rplot[(Rplot$generations + 10) %% 250 == 0,]
  Rplot2$generations <- (Rplot2$generations + 10)/250
  
  bigplotlist2[[5]] <- ggplot(Rplot2, aes(x = zero, y = ratio
                                          , linetype = interaction(idFunc)
                                          , colour = as.factor(generations)
                                          , group = interaction(idFunc, generations))) +
    labs(title = paste("D Ratio between beneficial and total metabolite concentration summary"), linetype = "starting function", colour = "phase", x = "number of beneficial metabolites", y = "beneficial ratio")+
    theme_classic()+
    facet_wrap(vars(ecost))+
    theme(strip.placement = "outside", legend.position="none")+
    # geom_vline(xintercept = c(1:3*250), linetype='dashed', color='red', size=0.5)+
    scale_colour_manual(values = cbbPalette)+
    stat_summary( geom = "smooth", se = TRUE)  
  
  for (j in c(1, 10)){
    Rtable2 <-  Rtable[Rtable$ecost == j,]
    Rplot           <- melt(Rtable2, id.vars = names)
    Rplot$variable  <- (as.numeric(Rplot$variable)-1)*recordingfreq
    names(Rplot)    <- c(names(Rplot)[1:length(names(bigdescriptions))], "generations", "ratio")
    Rplot$zero      <- 1023 - Rplot$zero
    Rplot$zexp      <- paste("scenario", Rplot$zexp)
    Rplot$idFunc[Rplot$idFunc == 1] <- "identical"
    Rplot$idFunc[Rplot$idFunc == 0] <- "different"
    
    ggplot(Rplot, aes(x = generations, y = ratio
                      , linetype = interaction(idFunc)
                      , colour = interaction(zero)
                      , group = run
    )) +
      labs(title = paste("Ratio enzyme cost ", j), , linetype = "starting function", colour = "number of beneficial metabolites", y = "beneficial ratio")+
      theme_classic()+
      facet_wrap(~forcats::fct_relevel(zexp, paste("scenario", 1:10)))+
      theme(strip.placement = "outside", legend.position="right")+
      geom_vline(xintercept = c(1:3*250), linetype='dashed', color='red', size=0.5)+
      scale_colour_manual(values=cbbPalette)+
      stat_summary( geom = "smooth", se = TRUE,
      )
    ggsave(paste("screening_Rplot_ALL_ecost", j, ".pdf"), width = 22, height = 13, units = "cm")
  }

  patchplots <- bigplotlist[[4]] / bigplotlist[[2]] / bigplotlist[[3]] / bigplotlist[[5]] / bigplotlist[[1]] + plot_layout(guides = 'collect')
  ggarrange(plotlist = patchplots, common.legend = TRUE, legend = "top")

  ggsave("screening_common_plot.pdf", width = 20, height = 40, units = "cm")

  patchplots2 <- bigplotlist2[[4]] / bigplotlist2[[2]] / bigplotlist2[[3]] / bigplotlist2[[5]] / bigplotlist2[[1]] + plot_layout(guides = 'collect')
  ggarrange(plotlist = patchplots2, common.legend = TRUE, legend = "top")
  
  ggsave("zerox_screening_common_plot.pdf", width = 20, height = 40, units = "cm")  
  
#maximum hit/absence of genes

hist <- cbind(bigdescriptions, !is.na(Ktable[(length(names) + 1):ncol(Ktable)]))

names(hist)[(length(names) + 1):ncol(hist)] <- (0:100 * 10)

hist3 <- melt(hist, id.vars = names)

hist3$ecost  <- paste(" ec.", hist3$ecost)
hist3$idFunc[hist3$idFunc == 1] <- "id"
hist3$idFunc[hist3$idFunc == 0] <- "diff"
hist3$zero   <- paste(1023 - hist3$zero, "ben.")

hist4 <- hist3[hist3$variable == 1000,]

ggplot(hist3, aes(x = as.numeric(variable)*10, fill = value)) +
  labs(title = "Allele presence in simulation", fill = "alleles present")+
  xlab("generations")+
  ylab("simulations")+
  facet_grid(zero~interaction(idFunc, ecost))+
  theme_bw() +
  scale_fill_manual(values=c("#D55E00", "#009E73"))+
  geom_bar() +
  theme(axis.text.x = element_text(angle = 90, vjust = 0.5, hjust=1))

ggsave("screening_gene_presence_through_time.pdf", units = "cm", width = 13, height = 8)


times_hit <- rowSums(maxgeneshit)
times_hit4 <- cbind(bigdescriptions,times_hit)

hitbar <- list()
  
names(times_hit4)[8] <- "hitcount"
  
hitbar[[1]]   <- ggplot(times_hit4, aes(x = hitcount, fill = interaction(ecost))) +
    labs(title = paste("Nr. times max loci hit per replicate"), fill = "enzyme cost")+
    xlab("times max loci hit")+
    ylab("nr. simulations")+
    theme_bw() +
  scale_fill_manual(values=c("#D55E00", "#009E73"))+
    geom_histogram(bins = 20, #breaks = c(0, 1:20 * 34)
    ) +
     # scale_y_log10()+
    theme(axis.text.x = element_text(angle = 90, vjust = 0.5, hjust=1))

ggarrange(plotlist = hitbar, common.legend = TRUE, legend = "bottom", ncol = 1, nrow = 1)
ggsave("screening_loci_maxhit.pdf", units = "cm", width = 13, height = 8)

#phaseplot

unique(bigdescriptions$zexp)

relevants2 <- relevants[!duplicated(relevants),]

relevants2 <- relevants[order(relevants$simnum),]
relevants3 <- relevants2[relevants2$simnum%%10 == 0 & (relevants2$simnum - 11800)%%60 >= 30,]
relevants3 <- relevants3[!duplicated(relevants3),]

relevants3$fitnessval[relevants3$fitnessval > 0] <- 'yes'
relevants3$fitnessval[relevants3$fitnessval == 0] <- 'no'
relevants3$zero <- as.factor(relevants3$zero)

relevants4 <- cbind(relevants3, rep(0, times = nrow(relevants3)))
names(relevants4)[8] <- "complexity"

relevants5 <- relevants4
values <- c(1, 2, 4, 8, 16, 32, 64, 128, 256, 512)
values2 <- c(512, 256, 128, 64, 32, 16, 8, 4,2,1)
dummy <- relevants4$metabolite

for(i in values2){
  relevants5[dummy%/%i >= 1, ]$complexity <- relevants5[dummy%/%i >= 1, ]$complexity + 1
  dummy[dummy%/%i >= 1 ] <- dummy[dummy%/%i >= 1 ] - i
}

relevants6 <- relevants5[relevants5$fitnessval == "yes",]
names(relevants6)[1] <- "run"

relevants7        <- merge(relevants6, bigdescriptions[unique(relevants6$run)-11200,], by = "run")
relevants7$zexp   <- paste("scen.", relevants7$zexp)
relevants7$zexp   <- factor(relevants7$zexp, levels=unique(as.character(relevants7$zexp)) )
relevants7$zero.y <- paste(1023 - relevants7$zero.y, "beneficial")
relevants8 <- relevants7[relevants7$zero.y == "100 beneficial",]
relevants8$complexity <- as.integer(relevants8$complexity)
relevants8$phase <- relevants8$phase+1

A <- ggplot(relevants8, aes(x = as.factor(phase), fill = as.factor(complexity)))+
  geom_bar() +
  labs(fill = "complexity", x = "phase", y = "metabolite")+
  scale_y_continuous(breaks = c(0, 50, 100))+
  scale_fill_viridis_d(option = "turbo")+
  facet_grid(zexp~zero.y)+
  theme_classic() +
  theme(strip.background.y = element_blank(),
        strip.text.y = element_blank()
        )

relevants8 <- relevants7[relevants7$zero.y == "10 beneficial",]
relevants8$complexity <- as.integer(relevants8$complexity)
relevants8$phase <- relevants8$phase+1

B <- ggplot(relevants8, aes(x = as.factor(phase), fill = as.factor(complexity)))+
  geom_bar() +
  # scale_fill_manual(values = cbbPalette)+
  labs(fill = "complexity", x = "phase")+
  scale_y_continuous(breaks = c(0, 5, 10))+
  scale_fill_viridis_d(option = "turbo")+
  facet_grid(zexp~zero.y)+
  theme_classic() +
  theme(strip.background.y = element_blank(),
        strip.text.y = element_blank(),
        axis.title.y=element_blank())

relevants8 <- relevants7[relevants7$zero.y == "1 beneficial",]
relevants8$complexity <- as.integer(relevants8$complexity)
relevants8$phase <- relevants8$phase+1

C <- ggplot(relevants8, aes(x = as.factor(phase), fill = as.factor(complexity)))+
  geom_bar() +
  labs(fill = "complexity", x = "phase")+
  scale_y_continuous(breaks = c(0, 1))+
  scale_fill_viridis_d(option = "turbo", guide = FALSE)+
  facet_grid(zexp~zero.y)+
  theme_classic() +
  theme(axis.title.y=element_blank())

A + B + C + plot_layout(guides = 'collect') & plot_annotation(title = 'Phase effects')

ggsave("screening_phaseplot.pdf", width = 14, height = 25, units = "cm")
```
