## Supplementary material for "A metabolic model to investigate the evolution of chemodiversity": Simulation code and files: figures.R

```
#I used R version 4.5.3

#load packages
require(reshape2)  #version1.4.5
require(ggplot2)   #version 4.0.2
require(ggpubr)    #version 0.6.3
require(patchwork) #version 1.3.2
require(forcats)   #version 1.0.1

sessionInfo()

cbbPalette    <- c("#000000", "#56B4E9", "#F0E442", "#009E73", "#D55E00", "#CC79A7", "#E69F00", "#0072B2") #colourblind-friendly palette used throughout

#prepare dataframe with information about the simulation parameters
names                  <- c("run", "set", "exp", "ecost", "zero", "zexp", "idFunc")
bigdescriptions        <- as.data.frame(matrix(nrow = 11200, ncol = length(names)))
names(bigdescriptions) <- names

recordingfreq <- 10 #number of generations between recordings of the state of the simulation


#fill dataframe with parameters and load data
batch <- 1
setwd(paste0("batch ", batch))

repNum        <- 10
numZeroVec    <- c(0, 1, 2, 3, 4, 5, 6)
enzymeCostVec <- c(1, 10)
experiments   <- c(0:3)
zexpvec       <- c(1:10)
idFunc        <- c(1)

seed <- 1
for (zex in zexpvec){
  for (exp in experiments){
    for (ecost in enzymeCostVec){
      for (nZero in numZeroVec){
        for (rep in 1:repNum){
          print(seed)
          bigdescriptions[seed,] <- c(seed, batch, exp+1, ecost, nZero, zex, idFunc)
          seed <- seed + 1
        }
      }
    }
  }
}

load("indices.RData")

Ktab        <- K$meanmedian #promiscuity
lttab       <- lins$ltt$`1` #sum of alleles across loci
totaltab    <- lins$tot     #total gene copies across loci
richness    <- diversities$richness #metaboilte richness
posMetRatio <- ratio$posMetRatio    #ratio of metabolites with beneficial effects
maxgeneshit <- allhits      #all occurences of simulations hitting the maximum number of loci
chemotypelist <- chemotypes

batch <- 2
setwd(paste0("../batch ", batch))

repNum        <- 10
numZeroVec    <- c(0, 1, 2, 3, 4, 5, 6)
enzymeCostVec <- c(1, 10)
experiments   <- c(0:3)
zexpvec       <- c(1:10)
idFunc        <- c(0)

for (zex in zexpvec){
  for (exp in experiments){
    for (ecost in enzymeCostVec){
      for (nZero in numZeroVec){
        for (rep in 1:repNum){
          print(seed)
          bigdescriptions[seed,] <- c(seed, batch, exp+1, ecost, nZero, zex, idFunc)
          seed <- seed + 1
        }
      }
    }
  }
}

load("indices.RData")

Ktab        <- rbind(Ktab, K$meanmedian)
lttab       <- rbind(lttab, lins$ltt$`1`)
totaltab    <- rbind(totaltab, lins$tot)
richness    <- rbind(richness, diversities$richness)
posMetRatio <- rbind(posMetRatio, ratio$posMetRatio)
maxgeneshit <- rbind(maxgeneshit, allhits)

#extract relevant data about which metabolites are beneficial from a file that records the exact fitness effects of the metabolites
relevants <- randFitness[randFitness$generation == 0 | randFitness$generation == 250 | randFitness$generation == 500 | randFitness$generation == 750, ]

#prepare plotlist for summary plots
bigplotlist <- list()
bigplotlist2 <- list()

#Prepare dataframe for phase-scenario plots
bigdescriptions2 <- unique(bigdescriptions[3:7])
df               <- as.data.frame(matrix(nrow = nrow(bigdescriptions2), ncol = 101))

#plot richness
richtable                    <- cbind(bigdescriptions, richness)
richtable[8:ncol(richtable)] <- richtable[8:ncol(richtable)] - 1 #exclude unmodified metabolite

richtable2 <- cbind(bigdescriptions2, df)

for(i in 1:nrow(bigdescriptions2)){
  richtable2[i,6:ncol(richtable2)] <- colMeans(richtable[richtable[4] == bigdescriptions2[i,2] & 
                                                           richtable[5] == bigdescriptions2[i,3] & 
                                                           richtable[6] == bigdescriptions2[i,4] &
                                                           richtable[7] == bigdescriptions2[i,5] ,8:ncol(richtable)], na.rm = TRUE) 
}
richplot           <- melt(richtable2, id.vars = names(bigdescriptions2))
richplot$variable  <- (as.numeric(richplot$variable)-1)*recordingfreq
names(richplot)    <- c(names(richplot)[1:length(names(bigdescriptions2))], "generations", "richness")
richplot$ecost     <- paste("enzyme cost ", richplot$ecost)
richplot$zero      <- 7 - richplot$zero
richplot$idFunc[richplot$idFunc == 1] <- "identical"
richplot$idFunc[richplot$idFunc == 0] <- "different"

#summary plot
bigplotlist[[1]] <- ggplot(richplot, aes(x = generations, y = richness
                    , linetype = interaction(idFunc)
                    , colour = interaction(zero))) +
  labs(title = paste("E Richness summary"), linetype = "starting function", colour = "number of beneficial metabolites")+
  theme_classic()+
  facet_wrap(vars(ecost))+
  theme(strip.placement = "outside", legend.position="bottom")+
  geom_vline(xintercept = c(1:3*250), linetype='dashed', color='red', size=0.5)+
  scale_colour_manual(values = cbbPalette)+
  stat_summary( geom = "smooth", se = TRUE)

richplot2 <- richplot[(richplot$generations + 10) %% 250 == 0,]
richplot2$generations <- (richplot2$generations + 10)/250

bigplotlist2[[1]] <- ggplot(richplot2, aes(x = zero, y = richness
                                         , linetype = interaction(idFunc)
                                         , colour = as.factor(generations)
                                         , group = interaction(idFunc, generations))) +
  labs(title = paste("E Richness summary"), linetype = "starting function", colour = "phase", x = "number of beneficial metabolites")+
  theme_classic()+
  facet_wrap(vars(ecost))+
  theme(strip.placement = "outside", legend.position="bottom")+
  # geom_vline(xintercept = c(1:3*250), linetype='dashed', color='red', size=0.5)+
  scale_colour_manual(values = cbbPalette)+
  stat_summary( geom = "smooth", se = TRUE)

# ggsave("richness_summary.pdf")

#plots split out per phase scenario
for (j in c(1, 10)){
  richtable2         <- richtable[richtable$ecost == j,]
  richplot           <- melt(richtable2, id.vars = names)
  richplot$variable  <- (as.numeric(richplot$variable)-1)*recordingfreq
  names(richplot)    <- c(names(richplot)[1:length(names(bigdescriptions))], "generations", "richness")
  richplot$zero      <- 7 - richplot$zero
  richplot$zexp      <- paste("scenario", richplot$zexp)
  richplot$idFunc[richplot$idFunc == 1] <- "identical"
  richplot$idFunc[richplot$idFunc == 0] <- "different"
  
  ggplot(richplot, aes(x = generations, y = richness
                      , linetype = interaction(idFunc)
                      , colour = interaction(zero)
                      , group = interaction(idFunc, zero, exp))) +
    labs(title = paste("Richness, Enzyme cost ", j), linetype = "starting function", colour = "number of beneficial metabolites")+
    theme_classic()+
    facet_wrap(~forcats::fct_relevel(zexp, paste("scenario", 1:10)))+
    theme(strip.placement = "outside", legend.position="right")+
    geom_vline(xintercept = c(1:3*250), linetype='dashed', color='red', size=0.5)+
    scale_colour_manual(values = cbbPalette)+
    stat_summary( geom = "smooth", se = TRUE)
  
  ggsave(paste0("richness_ecost_", j, ".pdf"), width = 33, height = 19, units = "cm")
  
  richplot2 <- richplot[(richplot$generations + 10) %% 250 == 0,]
  richplot2$generations <- (richplot2$generations + 10)/250
  
  ggplot(richplot2, aes(x = zero, y = richness
                       , linetype = interaction(idFunc)
                       , colour = interaction(generations)
                       , group = interaction(idFunc, generations, exp)
                       )) +
    labs(title = paste("Richness, Enzyme cost ", j), linetype = "starting function", colour = "phase", x = "number of beneficial metabolites")+
    theme_classic()+
    facet_wrap(~forcats::fct_relevel(zexp, paste("scenario", 1:10)))+
    theme(strip.placement = "outside", legend.position="right")+
    # geom_vline(xintercept = c(1:3*250), linetype='dashed', color='red', size=0.5)+
    scale_colour_manual(values = cbbPalette)+
    stat_summary( geom = "smooth", se = TRUE)
  ggsave(paste0("zerox_richness_ecost_", j, ".pdf"), width = 33, height = 19, units = "cm")
}


#plot sum of alleles accross loci
ltttable        <- cbind(bigdescriptions, lttab)
ltttable2       <- cbind(bigdescriptions2, df)

for(i in 1:nrow(bigdescriptions2)){
  ltttable2[i,6:ncol(ltttable2)] <- colMeans(ltttable[ltttable[4] == bigdescriptions2[i,2] & 
                                                        ltttable[5] == bigdescriptions2[i,3] & 
                                                        ltttable[6] == bigdescriptions2[i,4] &
                                                        ltttable[7] == bigdescriptions2[i,5] ,8:ncol(ltttable)], na.rm = TRUE) 
}
lttplot           <- melt(ltttable2, id.vars = names(bigdescriptions2))
lttplot$variable  <- (as.numeric(lttplot$variable)-1)*recordingfreq
names(lttplot)    <- c(names(lttplot)[1:length(names(bigdescriptions2))], "generations", "ltt")
lttplot$ecost     <- paste("enzyme cost ", lttplot$ecost)
lttplot$zero      <- 7 - lttplot$zero
lttplot$idFunc[lttplot$idFunc == 1] <- "identical"
lttplot$idFunc[lttplot$idFunc == 0] <- "different"
  
#summary plot
bigplotlist[[2]] <- ggplot(lttplot, aes(x = generations, y = ltt
                                         , linetype = interaction(idFunc)
                                         , colour = interaction(zero))) +
    labs(title = paste("B Sum of alleles summary"), linetype = "starting function", colour = "number of beneficial metabolites", y = "sum of alleles")+
    theme_classic()+
    facet_wrap(vars(ecost))+
    theme(strip.placement = "outside", legend.position="none")+
    geom_vline(xintercept = c(1:3*250), linetype='dashed', color='red', size=0.5)+
    scale_colour_manual(values = cbbPalette)+
    stat_summary( geom = "smooth", se = TRUE)

lttplot2 <- lttplot[(lttplot$generations + 10) %% 250 == 0,]
lttplot2$generations <- (lttplot2$generations + 10)/250

bigplotlist2[[2]] <- ggplot(lttplot2, aes(x = zero, y = ltt
                                           , linetype = interaction(idFunc)
                                           , colour = as.factor(generations)
                                           , group = interaction(idFunc, generations))) +
  labs(title = paste("B Sum of alleles summary"), linetype = "starting function", colour = "phase", x = "number of beneficial metabolites", y = "sum of alleles")+
  theme_classic()+
  facet_wrap(vars(ecost))+
  theme(strip.placement = "outside", legend.position="none")+
  # geom_vline(xintercept = c(1:3*250), linetype='dashed', color='red', size=0.5)+
  scale_colour_manual(values = cbbPalette)+
  stat_summary( geom = "smooth", se = TRUE)

# ggsave("ltt_summary.pdf")

#plots split out by phase scenario
for (j in c(1, 10)){
  ltttable2         <- ltttable[ltttable$ecost == j,]
  lttplot           <- melt(ltttable2, id.vars = names)
  lttplot$variable  <- (as.numeric(lttplot$variable)-1)*recordingfreq
  names(lttplot)    <- c(names(lttplot)[1:length(names(bigdescriptions))], "generations", "ltt")
  lttplot$zero      <- 7 - lttplot$zero
  lttplot$zexp      <- paste("scenario", lttplot$zexp)
  lttplot$idFunc[lttplot$idFunc == 1] <- "identical"
  lttplot$idFunc[lttplot$idFunc == 0] <- "different"

  ggplot(lttplot, aes(x = generations, y = ltt
                                         , linetype = interaction(idFunc)
                                         , colour = interaction(zero)
                                         , group = interaction(idFunc, zero, exp))) +
      labs(title = paste("Sum of alleles, Enzyme cost ", j), linetype = "starting function", colour = "number of beneficial metabolites", y = "sum of alleles")+
      theme_classic()+
      facet_wrap(~forcats::fct_relevel(zexp, paste("scenario", 1:10)))+
      theme(strip.placement = "outside", legend.position="right")+
      geom_vline(xintercept = c(1:3*250), linetype='dashed', color='red', size=0.5)+
      scale_colour_manual(values = cbbPalette)+
      stat_summary( geom = "smooth", se = TRUE)
  
  ggsave(paste("ltt_all_ecost", j, ".pdf"), width = 33, height = 19, units = "cm")
  
  lttplot2 <- lttplot[(lttplot$generations + 10) %% 250 == 0,]
  lttplot2$generations <- (lttplot2$generations + 10)/250
  
  ggplot(lttplot2, aes(x = zero, y = ltt
                        , linetype = interaction(idFunc)
                        , colour = interaction(generations)
                        , group = interaction(idFunc, generations, exp)
  )) +
    labs(title = paste("Sum of alleles, Enzyme cost ", j), linetype = "starting function", colour = "phase", x = "number of beneficial metabolites", y = "sum of alleles")+
    theme_classic()+
    facet_wrap(~forcats::fct_relevel(zexp, paste("scenario", 1:10)))+
    theme(strip.placement = "outside", legend.position="right")+
    # geom_vline(xintercept = c(1:3*250), linetype='dashed', color='red', size=0.5)+
    scale_colour_manual(values = cbbPalette)+
    stat_summary( geom = "smooth", se = TRUE)
  ggsave(paste0("zerox_ltt_all_ecost_", j, ".pdf"), width = 33, height = 19, units = "cm")
}

  
#plot total gene copies across loci
totaltable  <- cbind(bigdescriptions, totaltab)
totaltable2       <- cbind(bigdescriptions2, df)
for(i in 1:nrow(bigdescriptions2)){
  totaltable2[i,6:ncol(totaltable2)] <- colMeans(totaltable[totaltable[4] == bigdescriptions2[i,2] & 
                                                              totaltable[5] == bigdescriptions2[i,3] & 
                                                              totaltable[6] == bigdescriptions2[i,4] &
                                                              totaltable[7] == bigdescriptions2[i,5] ,8:ncol(totaltable)], na.rm = TRUE) 
}
totalplot           <- melt(totaltable2, id.vars = names(bigdescriptions2))
totalplot$variable  <- (as.numeric(totalplot$variable)-1)*recordingfreq
names(totalplot)    <- c(names(totalplot)[1:length(names(bigdescriptions2))], "generations", "total")
totalplot$ecost     <- paste("enzyme cost ", totalplot$ecost)
totalplot$zero      <- 7 - totalplot$zero
totalplot$idFunc[totalplot$idFunc == 1] <- "identical"
totalplot$idFunc[totalplot$idFunc == 0] <- "different"
  
#summary plot
bigplotlist[[3]] <- ggplot(totalplot, aes(x = generations, y = total
                    , linetype = interaction(idFunc)
                    , colour = interaction(zero))) +
    labs(title = paste("C Number of gene copies summary"), linetype = "starting function", colour = "number of beneficial metabolites", y = "number of gene copies")+
    theme_classic()+
    facet_wrap(vars(ecost))+
    theme(strip.placement = "outside", legend.position="none")+
    geom_vline(xintercept = c(1:3*250), linetype='dashed', color='red', size=0.5)+
    scale_colour_manual(values = cbbPalette)+
    stat_summary( geom = "smooth", se = TRUE)

totalplot2 <- totalplot[(totalplot$generations + 10) %% 250 == 0,]
totalplot2$generations <- (totalplot2$generations + 10)/250

bigplotlist2[[3]] <- ggplot(totalplot2, aes(x = zero, y = total
                                        , linetype = interaction(idFunc)
                                        , colour = as.factor(generations)
                                        , group = interaction(idFunc, generations))) +
  labs(title = paste("C Number of gene copies summary"), linetype = "starting function", colour = "phase", x = "number of beneficial metabolites", y = "number of gene copies")+
  theme_classic()+
  facet_wrap(vars(ecost))+
  theme(strip.placement = "outside", legend.position="none")+
  # geom_vline(xintercept = c(1:3*250), linetype='dashed', color='red', size=0.5)+
  scale_colour_manual(values = cbbPalette)+
  stat_summary( geom = "smooth", se = TRUE)
  
# ggsave("total_summary.pdf")
  
#plots split out by phase scenario
for (j in c(1, 10)){
  totaltable2         <- totaltable[totaltable$ecost == j,]
  totalplot           <- melt(totaltable2, id.vars = names)
  totalplot$variable  <- (as.numeric(totalplot$variable)-1)*recordingfreq
  names(totalplot)    <- c(names(totalplot)[1:length(names(bigdescriptions))], "generations", "total")
  totalplot$zero      <- 7 - totalplot$zero
  totalplot$zexp      <- paste("scenario", totalplot$zexp)
  totalplot$idFunc[totalplot$idFunc == 1] <- "identical"
  totalplot$idFunc[totalplot$idFunc == 0] <- "different"
    
  ggplot(totalplot, aes(x = generations, y = total
                        , linetype = interaction(idFunc)
                        , colour = interaction(zero)
                        , group = interaction(idFunc, zero, exp))) +
      labs(title = paste("Number of gene copies, Enzyme cost ", j), linetype = "starting function", colour = "number of beneficial metabolites", y = "number of gene copies")+
      theme_classic()+
      facet_wrap(~forcats::fct_relevel(zexp, paste("scenario", 1:10)))+
      theme(strip.placement = "outside", legend.position="right")+
      geom_vline(xintercept = c(1:3*250), linetype='dashed', color='red', size=0.5)+
      scale_colour_manual(values = cbbPalette)+
      stat_summary( geom = "smooth", se = TRUE)
    ggsave(paste("total_all_ecost", j, ".pdf"), width = 33, height = 19, units = "cm")
    
    totalplot2 <- totalplot[(totalplot$generations + 10) %% 250 == 0,]
    totalplot2$generations <- (totalplot2$generations + 10)/250
    
    ggplot(totalplot2, aes(x = zero, y = total
                          , linetype = interaction(idFunc)
                          , colour = interaction(generations)
                          , group = interaction(idFunc, generations, exp)
    )) +
      labs(title = paste("Number of gene copies, Enzyme cost ", j), linetype = "starting function", colour = "phase", x = "number of beneficial metabolites", y = "number of gene copies")+
      theme_classic()+
      facet_wrap(~forcats::fct_relevel(zexp, paste("scenario", 1:10)))+
      theme(strip.placement = "outside", legend.position="right")+
      # geom_vline(xintercept = c(1:3*250), linetype='dashed', color='red', size=0.5)+
      scale_colour_manual(values = cbbPalette)+
      stat_summary( geom = "smooth", se = TRUE)
    ggsave(paste0("zerox_total_all_ecost_", j, ".pdf"), width = 33, height = 19, units = "cm")
} 

#promiscuity
Ktable        <- cbind(bigdescriptions, Ktab)
Ktable2       <- cbind(bigdescriptions2, df)

for(i in 1:nrow(bigdescriptions2)){
  Ktable2[i,6:ncol(Ktable2)] <- colMeans(Ktable[Ktable[4] == bigdescriptions2[i,2] & 
                                                  Ktable[5] == bigdescriptions2[i,3] & 
                                                  Ktable[6] == bigdescriptions2[i,4] &
                                                  Ktable[7] == bigdescriptions2[i,5] ,8:ncol(Ktable)], na.rm = TRUE) 
}
Kplot           <- melt(Ktable2, id.vars = names(bigdescriptions2))
Kplot$variable  <- (as.numeric(Kplot$variable)-1)*recordingfreq
names(Kplot)    <- c(names(Kplot)[1:length(names(bigdescriptions2))], "generations", "promiscuity index")
Kplot$zero      <- 7 - Kplot$zero
Kplot$ecost     <- paste("enzyme cost ", Kplot$ecost)
Kplot$idFunc[Kplot$idFunc == 1] <- "identical"
Kplot$idFunc[Kplot$idFunc == 0] <- "different"
    
#summary plot
bigplotlist[[4]] <- ggplot(Kplot, aes(x = generations, y = `promiscuity index`
                  , linetype = interaction(idFunc)
                  , colour = interaction(zero)
  )) +
    labs(title = paste0("A Promiscuity index summary"), linetype = "starting function", colour = "number of beneficial metabolites")+
    theme_classic()+
    facet_wrap(vars(ecost))+
    theme(strip.placement = "outside", legend.position="none")+
    geom_vline(xintercept = c(1:3*250), linetype='dashed', color='red', size=0.5)+
    scale_colour_manual(values=cbbPalette)+
    stat_summary( geom = "smooth", se = TRUE, alpha = 0.1)
    

Kplot2 <- Kplot[(Kplot$generations + 10) %% 250 == 0,]
Kplot2$generations <- (Kplot2$generations + 10)/250

bigplotlist2[[4]] <- ggplot(Kplot2, aes(x = zero, y = `promiscuity index`
                                           , linetype = interaction(idFunc)
                                           , colour = as.factor(generations)
                                           , group = interaction(idFunc, generations))) +
  labs(title = paste("A Promiscuity index summary"), linetype = "starting function", colour = "phase", x = "number of beneficial metabolites")+
  theme_classic()+
  facet_wrap(vars(ecost))+
  theme(strip.placement = "outside", legend.position="none")+
  # geom_vline(xintercept = c(1:3*250), linetype='dashed', color='red', size=0.5)+
  scale_colour_manual(values = cbbPalette)+
  stat_summary( geom = "smooth", se = TRUE)
# ggsave(paste("Kplots_summary.pdf"))

#plots split out by phase scenario
for (j in c(1, 10)){
  Ktable2         <- Ktable[Ktable$ecost == j,]
  Kplot           <- melt(Ktable2, id.vars = names)
  Kplot$variable  <- (as.numeric(Kplot$variable)-1)*recordingfreq
  names(Kplot)    <- c(names(Kplot)[1:length(names(bigdescriptions))], "generations", "promiscuity index")
  Kplot$zero      <- 7 - Kplot$zero
  Kplot$zexp      <- as.factor(paste("scenario", Kplot$zexp))
  Kplot$idFunc[Kplot$idFunc == 1] <- "identical"
  Kplot$idFunc[Kplot$idFunc == 0] <- "different"
  
  ggplot(Kplot, aes(x = generations, y = `promiscuity index`
                        , linetype = interaction(idFunc)
                        , colour = interaction(zero)
                        , group = interaction(idFunc, zero, exp)
  )) +
    labs(title = paste("Promiscuity index, Enzyme cost ", j), linetype = "starting function", colour = "number of beneficial metabolites")+
    theme_classic()+
    facet_wrap(~forcats::fct_relevel(zexp, paste("scenario", 1:10)))+
    theme(strip.placement = "outside", legend.position="right")+
    geom_vline(xintercept = c(1:3*250), linetype='dashed', color='red', size=0.5)+
    scale_colour_manual(values = cbbPalette)+
    stat_summary( geom = "smooth", se = TRUE)
  
  ggsave(paste("Ktable_all_ecost", j, ".pdf"), width = 33, height = 19, units = "cm")
  
  Kplot2 <- Kplot[(Kplot$generations + 10) %% 250 == 0,]
  Kplot2$generations <- (Kplot2$generations + 10)/250
  
  ggplot(Kplot2, aes(x = zero, y = `promiscuity index`
                        , linetype = interaction(idFunc)
                        , colour = interaction(generations)
                        , group = interaction(idFunc, generations, exp)
  )) +
    labs(title = paste("Promiscuity index, Enzyme cost ", j), linetype = "starting function", colour = "phase", x = "number of beneficial metabolites")+
    theme_classic()+
    facet_wrap(~forcats::fct_relevel(zexp, paste("scenario", 1:10)))+
    theme(strip.placement = "outside", legend.position="right")+
    # geom_vline(xintercept = c(1:3*250), linetype='dashed', color='red', size=0.5)+
    scale_colour_manual(values = cbbPalette)+
    stat_summary( geom = "smooth", se = TRUE)
  ggsave(paste0("zerox_ktable_all_ecost_", j, ".pdf"), width = 33, height = 19, units = "cm")
}  

#ratio of beneficial metabolites
Rtable  <- cbind(bigdescriptions, posMetRatio)
Rtable2       <- cbind(bigdescriptions2, df)

for(i in 1:nrow(bigdescriptions2)){
  Rtable2[i,6:ncol(Rtable2)] <- colMeans(Rtable[Rtable[4] == bigdescriptions2[i,2] & 
                                                  Rtable[5] == bigdescriptions2[i,3] & 
                                                  Rtable[6] == bigdescriptions2[i,4] &
                                                  Rtable[7] == bigdescriptions2[i,5] ,8:ncol(Rtable)], na.rm = TRUE) 
}
Rplot           <- melt(Rtable2, id.vars = names(bigdescriptions2))
Rplot$variable  <- (as.numeric(Rplot$variable)-1)*recordingfreq
names(Rplot)    <- c(names(Rplot)[1:length(names(bigdescriptions2))], "generations", "ratio")
Rplot$ecost     <- paste("enzyme cost ", Rplot$ecost)
Rplot$zero      <- 7 - Rplot$zero
Rplot$idFunc[Rplot$idFunc == 1] <- "identical"
Rplot$idFunc[Rplot$idFunc == 0] <- "different"

#summary plot
bigplotlist[[5]] <- ggplot(Rplot, aes(x = generations, y = ratio
                                      , linetype = interaction(idFunc)
                                      , colour = interaction(zero)
)) +
  labs(title = paste("D Ratio between beneficial and total metabolite concentration summary"),
       linetype = "starting function",
       colour = "number of beneficial metabolites", 
       y = "beneficial ratio")+
  theme_classic()+
  facet_wrap(vars(ecost))+
  theme(strip.placement = "outside", legend.position="none")+
  geom_vline(xintercept = c(1:3*250), linetype='dashed', color='red', size=0.5)+
  scale_colour_manual(values=cbbPalette)+
  stat_summary( geom = "smooth", se = TRUE)
# ggsave(paste("Rplot_summary.pdf"))

Rplot2 <- Rplot[(Rplot$generations + 10) %% 250 == 0,]
Rplot2$generations <- (Rplot2$generations + 10)/250

bigplotlist2[[5]] <- ggplot(Rplot2, aes(x = zero, y = ratio
                                           , linetype = interaction(idFunc)
                                           , colour = as.factor(generations)
                                           , group = interaction(idFunc, generations))) +
  labs(title = paste("D Ratio between beneficial and total metabolite concentration summary"), linetype = "starting function", colour = "phase", x = "number of beneficial metabolites", y = "beneficial ratio")+
  theme_classic()+
  facet_wrap(vars(ecost))+
  theme(strip.placement = "outside", legend.position="none")+
  # geom_vline(xintercept = c(1:3*250), linetype='dashed', color='red', size=0.5)+
  scale_colour_manual(values = cbbPalette)+
  stat_summary( geom = "smooth", se = TRUE)

#plots split out by phase scenario
for (j in c(1, 10)){
    Rtable2         <-  Rtable[Rtable$ecost == j,]
    Rplot           <- melt(Rtable2, id.vars = names)
    Rplot$variable  <- (as.numeric(Rplot$variable)-1)*recordingfreq
    names(Rplot)    <- c(names(Rplot)[1:length(names(bigdescriptions))], "generations", "ratio")
    Rplot$zero      <- 7 - Rplot$zero
    Rplot$zexp      <- paste("scenario", Rplot$zexp)
    Rplot$idFunc[Rplot$idFunc == 1] <- "identical"
    Rplot$idFunc[Rplot$idFunc == 0] <- "different"
    
    ggplot(Rplot, aes(x = generations, y = ratio
                                       , linetype = interaction(idFunc)
                                       , colour = interaction(zero)
                                       , group = interaction(idFunc, zero, exp)
    )) +
      labs(title = paste("Ratio enzyme cost ", j), , linetype = "starting function", colour = "number of beneficial metabolites", y = "beneficial ratio")+
      theme_classic()+
      facet_wrap(~forcats::fct_relevel(zexp, paste("scenario", 1:10)))+
      theme(strip.placement = "outside", legend.position="right")+
      geom_vline(xintercept = c(1:3*250), linetype='dashed', color='red', size=0.5)+
      scale_colour_manual(values=cbbPalette)+
      stat_summary( geom = "smooth", se = TRUE,
      )
  ggsave(paste("Rplot_ALL_ecost", j, ".pdf"), width = 33, height = 19, units = "cm")
  
  Rplot2 <- Rplot[(Rplot$generations + 10) %% 250 == 0,]
  Rplot2$generations <- (Rplot2$generations + 10)/250
  
  ggplot(Rplot2, aes(x = zero, y = ratio
                        , linetype = interaction(idFunc)
                        , colour = interaction(generations)
                        , group = interaction(idFunc, generations, exp)
  )) +
    labs(title = paste("Ratio Enzyme cost ", j), linetype = "starting function", colour = "phase", x = "number of beneficial metabolites", y = "beneficial ratio")+
    theme_classic()+
    facet_wrap(~forcats::fct_relevel(zexp, paste("scenario", 1:10)))+
    theme(strip.placement = "outside", legend.position="right")+
    # geom_vline(xintercept = c(1:3*250), linetype='dashed', color='red', size=0.5)+
    scale_colour_manual(values = cbbPalette)+
    stat_summary( geom = "smooth", se = TRUE)
  ggsave(paste0("zerox_Rplot_ALL_ecost_", j, ".pdf"), width = 33, height = 19, units = "cm")
}

#plot all the summary plots together
patchplots <- bigplotlist[[4]] / bigplotlist[[2]] / bigplotlist[[3]] / bigplotlist[[5]] / bigplotlist[[1]] + plot_layout(guides = 'collect')
ggarrange(plotlist = patchplots, common.legend = TRUE, legend = "top")

ggsave("common_plot.pdf", width = 20, height = 40, units = "cm")

patchplots2 <- bigplotlist2[[4]] / bigplotlist2[[2]] / bigplotlist2[[3]] / bigplotlist2[[5]] / bigplotlist2[[1]] + plot_layout(guides = 'collect')
ggarrange(plotlist = patchplots2, common.legend = TRUE, legend = "top")

ggsave("zerox_common_plot.pdf", width = 20, height = 40, units = "cm")
  
#chemotypes
alphas <- c(0.2, 0.4, 0.4, 0.47, 0.4, 0.7, 0.7, 1)
complexity <- rep(alphas, each = 101)
plist <- list()
for (i in 1:200){
  chemotypes[[i]][2] <- 1000 - rowSums(chemotypes[[i]][3:ncol(chemotypes[[i]])])
  names(chemotypes[[i]]) <- c("generations", "000", "100", "010", "110", "001", "101", "011", "111")
  f <- cbind( complexity, melt(chemotypes[[i]], "generations"))
  names(f)[3] <- "most complex metabolite"
  names(f)[4] <- "nr plants"
  plist[[i]] <- ggplot(f, aes(y = `nr plants`, fill = `most complex metabolite`, x = generations
                                   , alpha = complexity
  )) +
    theme(axis.text.x = element_text(angle = 45, vjust = 1, hjust=1)) +
    scale_fill_manual(values = cbbPalette)+
    geom_bar(stat = "identity")+
    theme_classic()+
    geom_vline(xintercept = c(1:3*250), linetype='dashed', color='red', size=0.5)+
    guides(alpha = "none", fill = guide_legend(override.aes = list(alpha =alphas)))
}

count <- 71
figure <- ggarrange(plotlist = plist[0:9+count], common.legend = TRUE, legend = "bottom", ncol = 5, nrow = 2)

annotate_figure(figure, top = text_grob(paste("Chemotypes" ), face = "bold", size = 14))
ggsave(paste("chemodiversity_example_1.pdf"), width = 19, height = 10, units = "cm")

count <- 101
figure <- ggarrange(plotlist = plist[0:9+count], common.legend = TRUE, legend = "bottom", ncol = 5, nrow = 2)

annotate_figure(figure, top = text_grob(paste("Chemotypes" ), face = "bold", size = 14))
ggsave(paste("chemodiversity_example_2.pdf"), width = 19, height = 10, units = "cm")

mets  <- read.csv("../batch 1/output/file_97_eCost_10_numZero_2_zeroEx_1_Ex_0_rep_7/metabolite")
mets2 <- mets[1:10] 
names(mets2) <- c("generation", "plant", "000", "100", "010", "110", "001", "101", "011", "111")
mets3 <- melt(mets2, id.vars = c("generation", "plant"))
names(mets3)[c(3,4)]    <- c("metabolite", "concentration")

#overview of a run
plots <- list()
chemotypelist[[97]][2] <- 1000 - rowSums(chemotypelist[[97]][3:ncol(chemotypelist[[97]])])
names(chemotypelist[[97]]) <- c("generations", "000", "100", "010", "110", "001", "101", "011", "111")
f <- cbind( complexity, melt(chemotypelist[[97]], "generations"))
names(f)[3] <- "most complex metabolite"
names(f)[4] <- "nr plants"
plots[[1]] <- ggplot(f, aes(y = `nr plants`, fill = `most complex metabolite`, x = generations, alpha = complexity)) +
  theme_bw() +
  labs(title = paste("A"))+
  scale_fill_manual(values = cbbPalette)+
  geom_bar(stat = "identity")+
  geom_vline(xintercept = c(1:3*250), linetype='dashed', color='red', size=0.5)+
  guides(alpha = "none", fill = guide_legend(override.aes = list(alpha =alphas)))

plots[[2]] <- ggplot(mets3, aes(x = generation, y = concentration
                                , colour = metabolite
)) +
  labs(title = paste("B"))+
  theme_classic() +
  geom_vline(xintercept = c(1:3*250), linetype='dashed', color='red', size=0.5)+
  scale_colour_manual(values=cbbPalette)+
  stat_summary( geom = "smooth", se = TRUE,
  )

k2 <- Ktable[97,]
names(k2)[(length(names) + 1):ncol(k2)] <- (0:100 * 10)
k3           <- melt(k2, id.vars = names)
k3$variable  <- (as.numeric(k3$variable)-1)*recordingfreq
names(k3)    <- c(names(k3)[1:length(names(bigdescriptions))], "generations", "promiscuity index")

plots[[3]] <- ggplot(k3, aes(x = generations, y = `promiscuity index`)) +
  geom_line()+
  labs(title = paste("C"))+
  theme_classic()+
  geom_vline(xintercept = c(1:3*250), linetype='dashed', color='red', size=0.5)+
  scale_colour_manual(values='black')

l2 <- ltttable[97,]
names(l2)[(length(names) + 1):ncol(l2)] <- (0:100 * 10)
l3           <- melt(l2, id.vars = names)
l3$variable  <- (as.numeric(l3$variable)-1)*recordingfreq
names(l3)    <- c(names(l3)[1:length(names(bigdescriptions))], "generations", "sum of alleles")

plots[[4]] <- ggplot(l3, aes(x = generations, y = `sum of alleles`)) +
  geom_line()+
  labs(title = paste("D"))+
  theme_classic()+
  geom_vline(xintercept = c(1:3*250), linetype='dashed', color='red', size=0.5)+
  scale_colour_manual(values='black')

r2 <- Rtable[97,]
names(r2)[(length(names) + 1):ncol(r2)] <- (0:100 * 10)
r3           <- melt(r2, id.vars = names)
r3$variable  <- (as.numeric(r3$variable)-1)*recordingfreq
names(r3)    <- c(names(r3)[1:length(names(bigdescriptions))], "generations", "ratio")

plots[[5]] <- ggplot(r3, aes(x = generations, y = ratio)) +
  geom_line()+
  labs(title = paste("E"), y = "beneficial ratio")+
  theme_classic()+
  geom_vline(xintercept = c(1:3*250), linetype='dashed', color='red', size=0.5)+
  scale_colour_manual(values='black')

ri2 <- richtable[97,]
names(ri2)[(length(names) + 1):ncol(ri2)] <- (0:100 * 10)
ri3           <- melt(ri2, id.vars = names)
ri3$variable  <- (as.numeric(ri3$variable)-1)*recordingfreq
names(ri3)    <- c(names(ri3)[1:length(names(bigdescriptions))], "generations", "richness")

plots[[6]] <- ggplot(ri3, aes(x = generations, y = richness)) +
  geom_line()+
  labs(title = paste("F"))+
  theme_classic()+
  geom_vline(xintercept = c(1:3*250), linetype='dashed', color='red', size=0.5)+
  scale_colour_manual(values='black')

g1 <- ggarrange(plotlist = plots[c(1:2)], legend = "bottom", ncol = 2, nrow = 1)
g2 <- ggarrange(plotlist = plots[c(3:6)], legend = "bottom", ncol = 4, nrow = 1)
ggarrange(plotlist = list(g1, g2), ncol = 1, nrow = 2, heights = c(3, 2))

ggsave("overview_of_a_run.pdf", width = 24, height = 13, units = "cm")

#presence of alleles

hist <- cbind(bigdescriptions, !is.na(Ktable[(length(names) + 1):ncol(Ktable)]))

names(hist)[(length(names) + 1):ncol(hist)] <- (0:100 * 10)

hist3 <- melt(hist, id.vars = names)

hist3$ecost  <- paste(" ec.", hist3$ecost)
hist3$idFunc[hist3$idFunc == 1] <- "id"
hist3$idFunc[hist3$idFunc == 0] <- "diff"
hist3$zero   <- paste(7 - hist3$zero, "ben.")

ggplot(hist3, aes(x = as.numeric(variable)*10, fill = value)) +
  labs(title = "Allele presence in simulation", fill = "alleles present")+
  xlab("generations")+
  ylab("simulations")+
  facet_grid(zero~interaction(idFunc, ecost))+
  theme_bw() +
  scale_fill_manual(values=c("#D55E00", "#009E73"))+
  geom_bar() +
  theme(axis.text.x = element_text(angle = 90, vjust = 0.5, hjust=1))

ggsave("gene_presence_through_time.pdf", units = "cm", width = 13, height = 13)

#max loci hit
times_hit  <- rowSums(maxgeneshit)
times_hit4 <- cbind(bigdescriptions,times_hit)
names(times_hit4)[7] <- "hitcount"  

ggplot(times_hit4, aes(x = times_hit, fill = interaction(ecost))) +
    labs(title = paste("Nr. times max loci hit per replicate"), fill = "enzyme cost")+
    xlab("times max loci hit")+
    ylab("nr. simulations")+
    theme(legend.position = "bottom")+
    theme_bw() +
    scale_fill_manual(values=c("#D55E00", "#009E73"))+
    geom_histogram(bins = 20, #breaks = c(0, 1:20 * 34)
    ) +
    theme(axis.text.x = element_text(angle = 90, vjust = 0.5, hjust=1))

ggsave("loci_maxhit.pdf", units = "cm", width = 13, height = 8)

#phaseplot
relevants2 <- relevants[(relevants$simnum %% 10 - 2) == 0 & (relevants$simnum %% 560 - 2) < 70 & relevants$fseed == 0,]
relevants3 <- relevants2[!duplicated(relevants2),]
relevants3$fitnessval[relevants3$fitnessval > 0] <- 'yes'
relevants3$fitnessval[relevants3$fitnessval == 0] <- 'no'
relevants3 <- relevants3[order(relevants3$simnum),]

relevants5 <- relevants3
relevants5$simnum <- rep(rep(7:1, each = 32), 10)
relevants5$phase  <- paste("phase", relevants5$phase + 1)
relevants5$zero   <- paste("scen.", relevants5$zero)
relevants5$zero   <- factor(relevants5$zero, levels=unique(as.character(relevants5$zero)) )
relevants5$metabolite[relevants5$metabolite == 0] <- "000"
relevants5$metabolite[relevants5$metabolite == 1] <- "100"
relevants5$metabolite[relevants5$metabolite == 2] <- "010"
relevants5$metabolite[relevants5$metabolite == 3] <- "110"
relevants5$metabolite[relevants5$metabolite == 4] <- "001"
relevants5$metabolite[relevants5$metabolite == 5] <- "101"
relevants5$metabolite[relevants5$metabolite == 6] <- "011"
relevants5$metabolite[relevants5$metabolite == 7] <- "111"

ggplot(relevants5, aes(reorder(relevants3$zero, zero), y = reorder(metabolite, relevants3$metabolite), fill = fitnessval, x = simnum)) +
  theme_classic()+
  facet_grid(zero~phase)+
  theme(strip.placement = "outside")+
  labs(title = "Phase effects", fill = "has beneficial effect")+
  xlab("nr. beneficial metabolites")+
  ylab("metabolite")+
  scale_x_continuous(breaks = c(0:7)) +
  scale_fill_manual(values=c("#D55E00", "#009E73"))+
  geom_tile()

ggsave("phase_tileplot.pdf", units = "cm", width = 13, height = 27)
```
