## Supplementary material for "A metabolic model to investigate the evolution of chemodiversity": Simulation code and files: extract_indices.R

```
#!/usr/bin/Rscript --vanilla

args <- commandArgs(TRUE)
library(foreach)
library(doMC)
detectCores()
registerDoMC(20) #set by user
setwd(paste0("../../R"))

source("richness.R")
source("kcat_km.R")
source("lttstats.R")
source("make_tables.R")
source("posEffectRatio.R")

require(stringr)
require(reshape2)
# require(ape)

setwd(paste0("../simulations/robustness"))

reactions     <- 3
simlength     <- 1000
recordingfreq <- 10
nrPhases      <- 5

files <- list.files(path = "output/")
files2 <- vector(length = length(files))

randFitness <- read.csv(paste0("random_fitness.csv"), sep=,, header=TRUE)

numSims              <- length(files)

diversities          <- list()
diversities$richness <- diversities$simpson <- diversities$whittaker <- as.data.frame(matrix(nrow = numSims, ncol = simlength/recordingfreq + 1))
K                    <- list()
K$meanmedian         <- as.data.frame(matrix(nrow = numSims, ncol = simlength/recordingfreq + 1))
lins                 <- list()
lins$ltt             <- list()
lins$ltt$`1` <- lins$ltt$`2` <- lins$ltt$`3` <- lins$ltt$`4` <- lins$ltt$`5` <- lins$tot <- as.data.frame(matrix(nrow = numSims, ncol = simlength/recordingfreq + 1))
ratio                <- list()
ratio$posMetRatio    <- as.data.frame(matrix(nrow = numSims, ncol = simlength/recordingfreq + 1))

comb <- function(x, ...) {
  lapply(seq_along(x),
         function(i) c(x[[i]], lapply(list(...), function(y) y[[i]])))
}

rez <- foreach(i=20001:(20001-1 + length(files)), .combine='comb', .multicombine=TRUE,
               .init=list(list(), list(), list(), list(), list(), list(), list(),
                          list(), list()))%dopar%{
  print(i)
  files2[i]                 <- files[which(grepl(paste0("file_", i), files, fixed=TRUE) == TRUE)]

  metabolitedata            <- read.csv(paste0("output/", files2[i], "/metabolite"), sep=",", header=TRUE)
  genome                    <- read.csv(paste0("output/", files2[i], "/genome"), sep=",", header=TRUE)
  genes                     <- read.csv(paste0("output/", files2[i], "/genes"), sep=",", header=TRUE)
  fitTable                  <- randFitness[randFitness$simnum == i,]   
  fitTable                  <- fitTable[!duplicated(fitTable), ]

  popsize                   <- nrow(metabolitedata)/(simlength/10 + 1)

  metabolitedata            <- metabolitedata[1:(length(metabolitedata)-1)]

  richness     <- met_richness(metabolitedata, nrPlants = popsize)

  tables       <- make_tables(genome)

  posRatio     <- posEffectRatio(metabolitedata, fitTable, popsize)
  posMetRatio  <- posRatio$posMetRatio

  Ks           <- kcat_km(genome, genes, reactions, tables) # mean median normalized Kcat/Km
  meanmedian   <- Ks$meanmedian

  lin  <- lttstats(genome, tables)
  ltt1 <- lin$ltt$`1` #lineages through time
  ltt2 <- lin$ltt$`2`
  ltt3 <- lin$ltt$`3`
  ltt4 <- lin$ltt$`4`
  ltt5 <- lin$ltt$`5`
 
  tot <- lin$tot

  list(richness, meanmedian, ltt1, ltt2, ltt3, ltt4, ltt5, posMetRatio, tot)
}

load(file='indices1.RData')

print("part 1 done")

for(i in 1:length(files)){
  if (length(rez[[3]][[i]]) < 101){
    print(20000+i)
    print(length(rez[[3]][[i]]))
  }
}

for(i in 1:length(files)){
  diversities$richness[i,]     <- rez[[1]][[i]]
  
  K$meanmedian[i,]  <- rez[[2]][[i]]
  
  lins$ltt$`1`[i,]  <- rez[[3]][[i]]
  lins$ltt$`2`[i,]  <- rez[[4]][[i]]
  lins$ltt$`3`[i,]  <- rez[[5]][[i]]
  lins$ltt$`4`[i,]  <- rez[[6]][[i]]
  lins$ltt$`5`[i,]  <- rez[[7]][[i]]

  ratio$posMetRatio[i,] <- rez[[8]][[i]]

  lins$tot[i,]    <- rez[[9]][[i]]
}

print("part 2 done")

save.image(file='indices.RData')
```
