## Supplementary material for "A metabolic model to investigate the evolution of chemodiversity": Simulation code and files: make_files.R

```
#!/usr/bin/Rscript --vanilla

setwd(paste0("../../R"))
source("morris_trajectories.R")
setwd(paste0("../simulations/robustness"))

set.seed(1)

parameters <- c("popsize", "idFunc", "baseConc", "loss", "dup", "maxloci", "mutation", "maxConcChange", "kmean", "kmode", "influx", "ecost", "maxfit", "zero")

trajectories <- pick_trajectories(parameters, 4, 50, 50)

save.image(file = "trajectories.RData")

print(paste0("nr files = ", length(trajectories) * nrow(trajectories[[1]])))

popsize       <- c(200, 600, 1000, 1400)
idFunc        <- c(0, 0, 1, 1)
baseConc      <- c(0.0005, 0.001, 0.0015, 0.002)
loss          <- c(0.005, 0.01, 0.015, 0.002)
dup           <- c(0.00002, 0.00006, 0.0001, 0.00014)
maxloci       <- c(40, 60, 80, 100)
mutation      <- c(0.00002, 0.00006, 0.0001, 0.00014)
maxConcChange <- c(0.0005, 0.001, 0.0015, 0.002)
kMean         <- c(-4.37, -4.14, -4.01, -3.91, -3.91, -3.68, -3.55, -3.45, -3.64, -3.41, -3.28, -3.18, -3.45, -3.22, -3.08, -2.99) #mu, four groups for four means
kMode         <- c(0.96, 0.68, 0.44, 0.0001, 1.18, 0.96, 0.81, 0.68, 1.29, 1.09, 0.96, 0.86, 1.36, 1.18, 1.06, 0.96) #sigma, four groups for four means:cycling through four modes four times
influx        <- c(0.0005, 0.001, 0.0015, 0.002)
ecost         <- c(1, 5, 10, 15)
maxfit        <- c(100, 500, 1000, 1500)
zero          <- c(0, 2, 4, 6)

seed <- 1
for (trajectory in 1:nrow(trajectories[[1]])){
  for (timestep in 1:length(trajectories)){
    cat(paste0("logFile logs/log_", 20000 + seed, "_trajectory_", trajectory, "_step_", timestep, ".txt"),
        paste0(""),
        paste0("iSeed ", 20000 + seed),
        paste0("initialKSeed 1"),
        paste0("fitnessSeed 0"),
        paste0("zeroSeed 1"),
        paste0("iPopSize ", popsize[trajectories[[timestep]][trajectory,1]]),
        paste0("iNumGenerations 1001"),
        paste0(""),
        paste0("iNumGenePairs 3"),
        paste0("identicalFunction ", idFunc[trajectories[[timestep]][trajectory,2]]),
        paste0("dBaseConcentration ", baseConc[trajectories[[timestep]][trajectory,3]]),
        paste0(""),
        paste0("dLossOdds ", loss[trajectories[[timestep]][trajectory,4]]),
        paste0("dDuplicationOdds ", dup[trajectories[[timestep]][trajectory,5]]),
        paste0("iMaxNumLoci ", maxloci[trajectories[[timestep]][trajectory,6]]),
        paste0("dMutationOdds ", mutation[trajectories[[timestep]][trajectory,7]]),
        paste0("dMaxConcentration 0.1"),
        paste0("dMaxConcentrationChange ", maxConcChange[trajectories[[timestep]][trajectory,8]]),
        paste0("dKMu ", kMean[(trajectories[[timestep]][trajectory,9] - 1) * 4 + trajectories[[timestep]][trajectory,10]]),
        paste0("dKSigma ", kMode[(trajectories[[timestep]][trajectory,9] - 1) * 4 + trajectories[[timestep]][trajectory,10]]),
        paste0("iNumReactions 3"),
        paste0(""),
        paste0("setInitialK 0"),
        paste0("dInitialMetConcentration ", influx[trajectories[[timestep]][trajectory,11]] * 10),
        paste0("dInflux ", influx[trajectories[[timestep]][trajectory,11]]),
        paste0("dEfflux 0.1"),
        paste0("dODELength 100"),
        paste0("ODEstepper runge-kutta"),
        paste0("dSmallestODEStepSize 0.01"),
        paste0(""),
        paste0("fitnessHasEffect 1"),
        paste0("iNumPhases 5"),
        paste0("vecPhaseTransitionTimes1 250"),
        paste0("vecPhaseTransitionTimes2 500"),
        paste0("vecPhaseTransitionTimes3 750"),
        paste0("vecPhaseTransitionTimes4 1000"),
        paste0("dFitnessCostPerEnzyme ", ecost[trajectories[[timestep]][trajectory,12]]),
        paste0("fitnessIsRandom 1"),
        paste0("dMinimumFitnessEffect 0"),
        paste0("dMaximumFitnessEffect ", maxfit[trajectories[[timestep]][trajectory,13]]),
        paste0("iNumZeroFitness ", zero[trajectories[[timestep]][trajectory,14]]),
        paste0(""),
        paste0("outputODE 0"),
        paste0("outputMetabolites 1"),
        paste0("outputFilesLocation output/file_", 20000 + seed, "_trajectory_", trajectory, "_step_", timestep),
        paste0("iOutputInterval 10"),
        paste0("file ", 20000 + seed),
        
        sep = "\n",
        file = paste0("scripts_tmp/file_", 20000 + seed, "_trajectory_", trajectory, "_step_", timestep))
    seed <- seed + 1
  }
}
```
