## Supplementary material for "A metabolic model to investigate the evolution of chemodiversity": Simulation code and files: start_simulations.R

```
#!/usr/bin/Rscript --vanilla

args <- commandArgs(TRUE)
library(foreach)
library(doMC)
detectCores()
registerDoMC(20) #set by user

files <- list.files("scripts_tmp/")

randorder <- sample(20001:(20000+length(files)))

foreach(i=randorder)%dopar%{
  file <- files[which(grepl(paste0("file_", i, "_"), files, fixed=TRUE) == TRUE)]
  if (file.exists(paste0("output/", file)) == FALSE){
    system(paste0('echo "starting file "', i))
    system(paste0("./../program scripts_tmp/", file))
  }
}
```
