## Supplementary material for "A metabolic model to investigate the evolution of chemodiversity": Simulation code and files: variance_rank.R

```
require(reshape2)
require(ggplot2)
require(ggpubr)

#setwd("robustness") #Set working directory if nescessarry

#parameters
popsize       <- c(200, 600, 1000, 1400)
idFunc        <- c(0, 0, 1, 1)
baseConc      <- c(0.0005, 0.001, 0.0015, 0.002)
loss          <- c(0.005, 0.01, 0.015, 0.002)
dup           <- c(0.00002, 0.00006, 0.0001, 0.00014)
maxloci       <- c(40, 60, 80, 100)
mutation      <- c(0.00002, 0.00006, 0.0001, 0.00014)
maxConcChange <- c(0.0005, 0.001, 0.0015, 0.002)
kMean         <- c(-4.37, -4.14, -4.01, -3.91, -3.91, -3.68, -3.55, -3.45, -3.64, -3.41, -3.28, -3.18, -3.45, -3.22, -3.08, -2.99) #mu, four groups for four means
kMode         <- c(0.96, 0.68, 0.44, 0.0001, 1.18, 0.96, 0.81, 0.68, 1.29, 1.09, 0.96, 0.86, 1.36, 1.18, 1.06, 0.96) #sigma, four groups for four means:cycling through four modes four times
influx        <- c(0.0005, 0.001, 0.0015, 0.002)
ecost         <- c(1, 5, 10, 15)
maxfit        <- c(100, 500, 1000, 1500)
zero          <- c(0, 2, 4, 6)

#loading data created in extract_indices.R and start_simulations.R
load("trajectories.RData")
load("indices.RData")

trajectories2 <- trajectories

#match parameters
for (i in 1:length(trajectories)){
  trajectories[[i]][1] <- popsize[trajectories[[i]][,1]]
  trajectories[[i]][2] <- idFunc[trajectories[[i]][,2]]
  trajectories[[i]][3] <- baseConc[trajectories[[i]][,3]]
  trajectories[[i]][4] <- loss[trajectories[[i]][,4]]
  trajectories[[i]][5] <- dup[trajectories[[i]][,5]]
  trajectories[[i]][6] <- maxloci[trajectories[[i]][,6]]
  trajectories[[i]][7] <- mutation[trajectories[[i]][,7]]
  trajectories[[i]][8] <- maxConcChange[trajectories[[i]][,8]]
  trajectories[[i]][9] <- kMean[trajectories[[i]][,9]]
  trajectories[[i]][10] <- kMode[trajectories[[i]][,10]]
  trajectories[[i]][11] <- influx[trajectories[[i]][,11]]
  trajectories[[i]][12] <- ecost[trajectories[[i]][,12]]
  trajectories[[i]][13] <- maxfit[trajectories[[i]][,13]]
  trajectories[[i]][14] <- zero[trajectories[[i]][,14]]
}

#add parameter descriptions to output
bigdescriptions <- as.data.frame(matrix(nrow = nrow(trajectories[[1]])*length(trajectories), ncol = length(parameters)))
names(bigdescriptions) <- parameters

bigdescriptions2 <- bigdescriptions

for (i in 1:length(trajectories)){
bigdescriptions[0:(nrow(trajectories[[1]])-1)*length(trajectories) + i,] <- trajectories[[i]]
}

for (i in 1:length(trajectories2)){
  bigdescriptions2[0:(nrow(trajectories2[[1]])-1)*length(trajectories2) + i,] <- trajectories2[[i]]
}

replacement = TRUE #since idFunc can actually only have two values, we consolidate the four values of idFunc into two and change the calculations accordingly

if (replacement == TRUE){
  bigdescriptions2$idFunc[which(bigdescriptions2$idFunc == 1)] <- 2
  bigdescriptions2$idFunc[which(bigdescriptions2$idFunc == 4)] <- 3 
}

#retrieve output
diversities2 <- list()
ratio2       <- list()

Ktab <- K$meanmedian
lttab <- lins$ltt$`1`
diversities2$richness  <- diversities$richness
ratio2$posMetRatio     <- ratio$posMetRatio

#prepare to calculate variance
richValsTable <- list()
mu     <- list(richness = 0, promiscuity = 0, ltt = 0, ratio = 0)
muStar <- list(richness = 0, promiscuity = 0, ltt = 0, ratio = 0)
sigma  <- list(richness = 0, promiscuity = 0, ltt = 0, ratio = 0)

#Take from end of simulation
pos <- 100

richValsTable[[1]] <- cbind(bigdescriptions2, diversities2$richness[pos])
richValsTable[[2]] <- cbind(bigdescriptions2, Ktab[pos])
richValsTable[[3]] <- cbind(bigdescriptions2, lttab[pos])
richValsTable[[4]] <- cbind(bigdescriptions2, ratio2$posMetRatio[pos])

metrics <- c("Richness", "Promiscuity", "SoA", "Ratio of beneficial metabolites")

for (metric in 1:4){
  nochangeIdFunc <- 0
  richValsTable[[metric]][is.na(richValsTable[[metric]][15]),15] <- 0

#calculate variance here:
  dfF <- as.data.frame(matrix(nrow = nrow(trajectories[[1]]), ncol = length(parameters)))

  dfF2 <- as.data.frame(matrix(nrow = nrow(trajectories[[1]]), ncol = length(parameters)))


  for (j in 0:(nrow(trajectories[[1]])-1)){
    for (i in 1:length(parameters)){
      calc <- bigdescriptions2[i + j * (length(parameters) + 1),] - bigdescriptions2[i + j * (length(parameters) + 1) + 1,]
      if (!any(calc != 0)){print(i)}
      if (!any(calc != 0) && replacement == TRUE){
        dfF[j+1,2] <- 0
        dfF2[j+1,2] <- 0
        nochangeIdFunc <- nochangeIdFunc + 1
      }
      dfF[j+1,which(calc != 0)] <- (richValsTable[[metric]][i + j * (length(parameters) + 1), ncol(richValsTable[[metric]])] - richValsTable[[metric]][i + j * (length(parameters) + 1) + 1, ncol(richValsTable[[metric]])])/sum(calc) #di(X)
      dfF2[j+1,which(calc != 0)] <- sum(calc) #delta
    
    }
  }

  for (i in 1:length(parameters)){
    mu[[metric]][i]     <- 1/nrow(trajectories[[1]]) * sum(dfF[i])
    muStar[[metric]][i] <- 1/nrow(trajectories[[1]]) * sum(abs(dfF[i]))
    sigma[[metric]][i]  <- sqrt(1/(nrow(trajectories[[1]]-1)) * sum((dfF[i] - mu[[metric]][i])^2))
  }
  if(replacement == TRUE){
    mu[[metric]][2] <- 1/(nrow(trajectories[[1]])-nochangeIdFunc) * sum(dfF[2])
    muStar[[metric]][2] <- 1/(nrow(trajectories[[1]])-nochangeIdFunc) * sum(abs(dfF[2]))
    sigma[[metric]][2]  <- sqrt(1/(nrow(trajectories[[1]])-nochangeIdFunc - 1) * sum((dfF[2] - mu[[metric]][2])^2)) 
  }
}

#Plot variance
plotlist <- list()

parameters2 <- c("population size", "id. init. function", "initial enz. concentration", "gene loss rate", "duplication rate", "max. nr. loci",
"mutation rate", "max. enzyme conc. change", "mean kinetic dist.", "mode kinetic dist.", "metabolite influx", "enzyme cost",
"max. fitness effect", "nr. of n.b. met.")

for (i in 1:4){
toPlot <- as.data.frame(cbind(parameters2, mu[[i]], muStar[[i]], sigma[[i]]))
toPlot[2] <- as.numeric(toPlot[2][[1]])
toPlot[3] <- as.numeric(toPlot[3][[1]])
toPlot[4] <- as.numeric(toPlot[4][[1]])
names(toPlot) <- c("parameter", expression(mu), "muStar", "sigma")

plotlist[[i]] <- ggplot(toPlot, aes(x = sigma, y = muStar
                     , shape = parameter
                   
))+
  geom_point()+
  labs(title = paste(metrics[i]),
       y = expression(mu*"*"),
       x = expression(sigma))+
  scale_shape_manual(values = c(1:14))+
  theme_classic()
}
ggarrange(plotlist = plotlist, common.legend = TRUE, legend = "bottom", ncol = 2, nrow = 2)

ggsave(paste0("morris_variance_", pos, ".pdf"), width = 22, height = 13, units = "cm")

#calculate savage scores and order factors according to the sum of  their scores
scores   <- rep(0, 14)

for (i in 1:4){
  smallbig <- rank(muStar[[i]])  #rank the variances of the parameters for the metric
  position <- 1
  for (j in smallbig){
    scores[position] <- scores[position]  + sum(1/((15-j):14)) #add savage score based on ranking in individual metric
    position <- position + 1
  }
}

final_order <- 15 - rank(scores)

#parameters in order of influence
order_table2 <- cbind(parameters2, scores, final_order)
```
