## Supplementary figures and images for "A metabolic model to investigate the evolution of chemodiversity"

### morris_variance_100.pdf

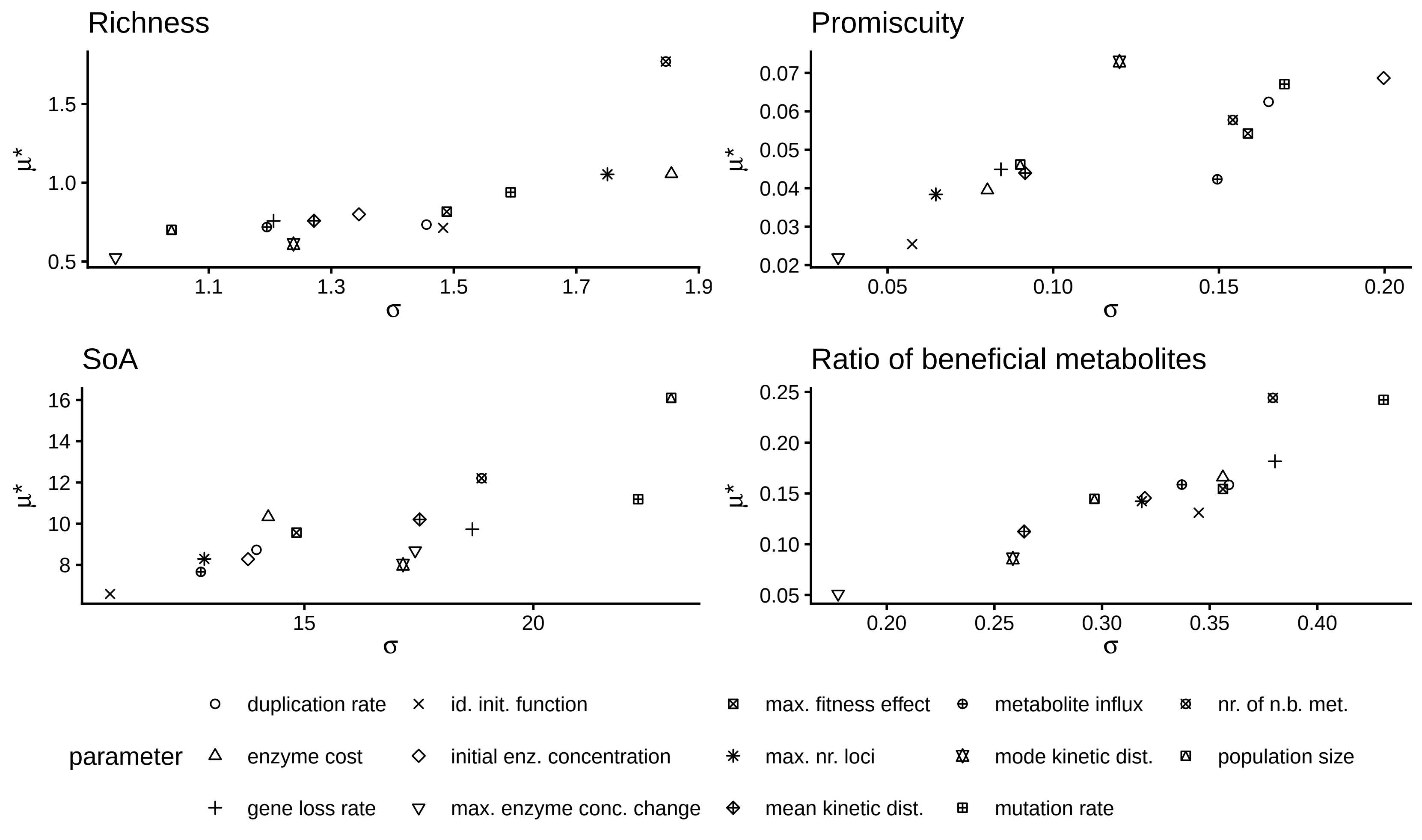
